# A Palette of Bridged Bicycle-Strengthened Fluorophores

**DOI:** 10.1101/2024.12.20.629585

**Authors:** Junwei Zhang, Kecheng Zhang, Kui Wang, Bo Wang, Siyan Zhu, Hongping Qian, Yumiao Ma, Mengling Zhang, Tianyan Liu, Peng Chen, Yuan Shen, Yunzhe Fu, Shilin Fang, Xinxin Zhang, Peng Zou, Wulan Deng, Yu Mu, Zhixing Chen

## Abstract

Organic fluorophores are the keystone of advanced biological imaging. The vast chemical space of fluorophores has been extensively explored in seek of molecules with ideal properties. However, within the current molecular constraints, there appears to be a trade-off between high brightness, robust photostability, and tunable biochemical properties. Herein we report a general strategy to systematically boost the performance of donor-acceptor-type fluorophores by leveraging SO_2_ and O-substituted azabicyclo[3.2.1] octane auxochromes. These bicyclic heterocycles give rise to a collection of ‘Bridged’ dyes (BD) spanning the UV and visible range with top-notch quantum efficiencies, enhanced water solubility, and tunable cell-permeability. Notably, these azabicyclic fluorophores showed remarkable photostability than its tetramethyl or azatidine analogue, at the same time completely resistant to oxidative photobluing rendered by the Bredt’s rule. Functionalized BD dyes are tailored for applications in single-molecule imaging, super-resolution imaging (STED and SIM) in fixed or live mammalian cells and plant cells, and live zebrafish imaging or chemigenetic voltage imaging. Synergizing with advanced imaging methods, the bridge bicycle dyes represent a versatile palette for biological researches.

## Introduction

In the pasted two decades, advanced fluorescence imaging techniques, featuring a spatial resolution beyond the Abbe diffraction limit and a temporal resolution near millisecond, have democratized as routine tools for biological research^1–4^. In parallel, the expanding palette of organic fluorophores^5–8^ and modern labeling strategies^9–12^ are supplanting fluorescent protein^13,14^ to meet the growing demand for 4D imaging on photon budget and multiplexed labeling. High brightness, superior photophysical, chemical, and spectral stability, tailored hydrophilicity, as well as excellent biocompatibility with and without excitation light, are the merit of an ideal fluorophore for contemporary imaging technologies. However, state-of-the-art fluorescent tools often have to compromise when trying to balance all the parameters. In other words, the quest for dyes with ideal properties calls for the exploration of wider biochemical space^15^.

The structural and electronic properties of auxochromes play a decisive role in the photophysical and chemical properties of the donor-acceptor-type (D-A type) fluorophores. Early designs generally improved the performance of fluorophores through structural rigidification and sulfonation (for example, **Alexa Fluor**^16^ and **ATTO Dyes**^17^). These classical pieces are still mainstays in the field of immunofluorescence. In parallel, while live-cell imaging and biorthogonal labeling strategies started to take the stage, a new trend of auxochrome engineering emerged to suppress the formation of twisted intramolecular charge transfer (TICT) to enhance the brightness (Fig. 1a)^18,19^. In 2008, the Foley group reported 7-azabicyclo[2.2.1]heptane as an auxochrome moiety with a small steric hindrance that can inhibit TICT state^20^. Moreover, this design deliberately leveraged bridgehead carbon, which cannot form double bonds, to eliminate the formation of photo-oxidation intermediates and completely avoid photobluing. More recently, the Lavis group systematically engineered azetidine substituted fluorophores, dubbed **JF** dyes^21–23^, which greatly improved brightness while preserving the compatibility with self-labeling protein tags^24,25^. JF dyes have now become a popular tool for live-cell and *in vivo* labeling. Yet, due to the high ring tensile force of azetidine and the presence of α-H atoms, their photobleaching (refers to photophysical stability) and photobluing(refers to spectral stability) are evident, particularly under light-intensive microscopes^26,27^. Subsequently, deuterated auxochromes were devised by the Lavis group^28^ and the Broichhagen group^29^ to improve photostability and spectral stability. Meanwhile, new strategies to prevent TICT state by adjusting electronic effects were reported by the Xiao ^30^ and Guo^31^ groups. Although these dyes have fluorescence quantum yields close to the theoretical limit, their photostability drastically decreased. At the same time, the Hell group reported triarylmethane fluorophores bearing auxochromes without alpha-H to effectively resistant oxidative photobluing, at the cost of installing a bulky yet hydrophobic tert-butyl group^32^. In addition to issues of brightness and photostability, the water solubility and biocompatibility of fluorophores are gradually receiving more attention. Gibbs group introduced a negatively charged sulfonate moiety to balance the cationic charge of rhodamine scaffolds to obtain a series of **ORFluor** with improved hydrophilicity while maintaining photophysical properties and cell-permeability^33^. Along the line of auxochrome engineering, our group leveraged the hydrophilic morpholino auxochromes^34,35^ to address the non-specific membrane staining issue of soluble Zinc indicators, which enabled multiplexed time-lapse imaging of orchestrated insulin-secretion^36^. While we are aware that the morpholino auxochromes compromise the brightness of rhodamine dyes, this initiative prompted us to further develop the next-generation molecular motifs that can generally render dyes with high brightness, high photostability, and spectral stability, yet customizable hydrophilicity for various applications.

**Fig. 1.**
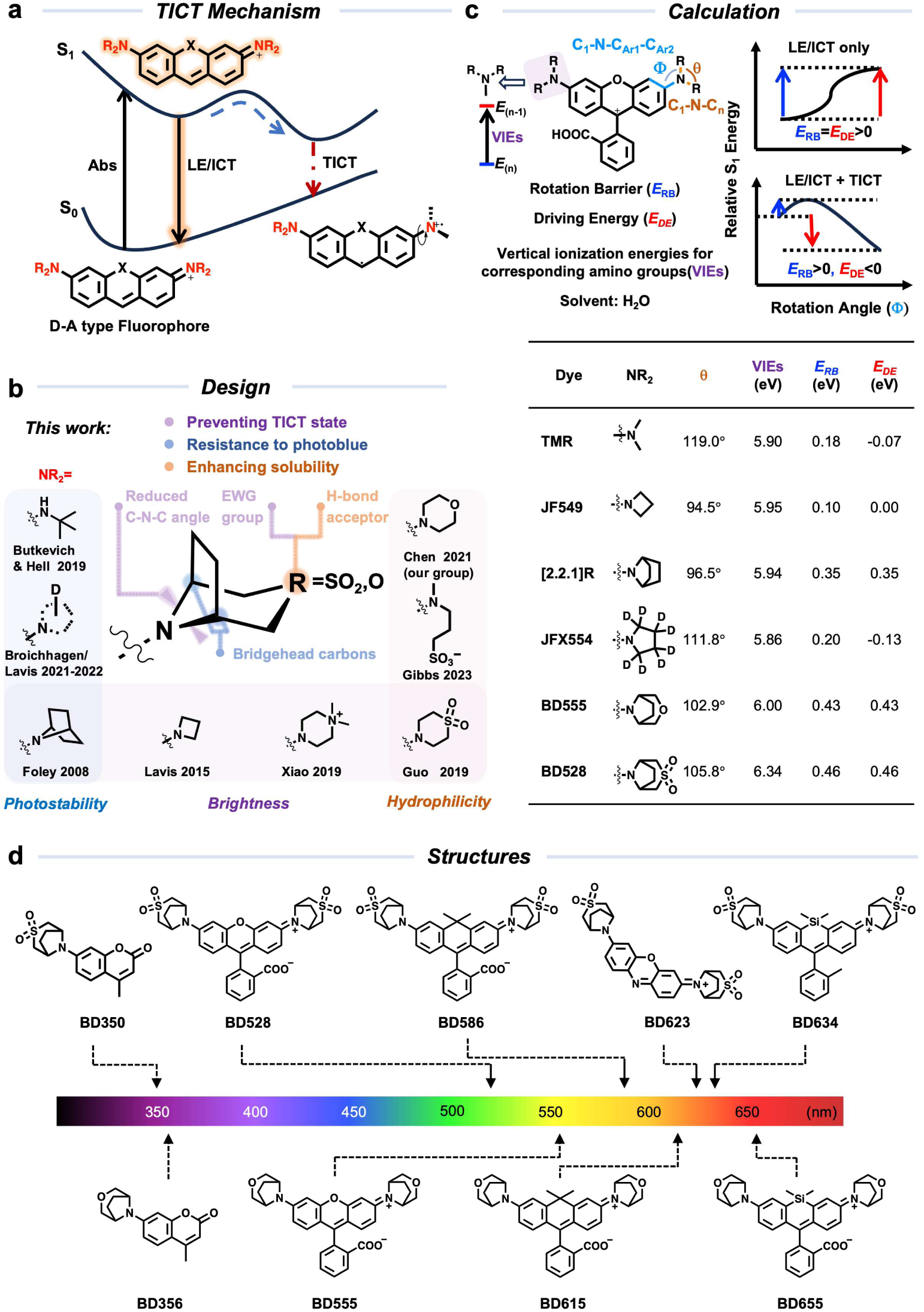
Development of fluorophores with bridged bicyclic auxochromes. (a) Twisted intramolecular charge transfer (TICT) mechanism, LE refers to local excited state while ICT refers to intermolecular charge transfer; (b) Selected auxochromes and their properties; (c) Theoretical models and calculation results;

Here we report a palette of bridged bicycle-strengthened fluorophores (**BriDye**, abbreviated to **“BD”**) with improved quantum efficiencies and uncompromised photo- and spectral stability that spans the UV and visible range. The **BD** family is composed of two series of fluorophores by using SO_2_ or O-substituted azabicyclo[3.2.1] octane auxochromes. Both dyes exhibit excellent photophysical properties while functionalized derivatives of **BD** dyes possess different hydrophilicity and membrane permeability to meet the demands of *in vitro* bioconjugation and live-cell protein labeling, respectively. By analyzing the crystal structure of the **BD** dye-protein binding complex, we have discovered new polar interactions between auxochromes and protein residues. We also showcase that **BD** dyes are privileged scaffolds for hybrid sensors and functional neuronal imaging. Furthermore, bioavailable **BD** dyes showed outstanding performance on general state-of-the-art imaging technologies (including but not limited to single-molecule imaging, stimulated emission depletion microscopy, or structure illumination microscopy) in fixed or live mammalian cells, plant cells, and live animals for structural imaging.

## Results

### Rational design of bridged bicycle-strengthened fluorophores

We seek to synergistically merge the biocompatibility of morpholino auxochrome, the brightness of azetidine or pyrrolidine auxochrome, and the photo/spectral stability of bridged 7-azabicyclo[2.2.1]heptane auxochrome. Along this line, we speculate that SO_2_ or O substituted azabicyclo[3.2.1] octane ^37^ are viable auxochromes with synthetic accessibility^37^ (Fig. 1b). These designs are expected to bring the following benefits: 1. The electron withdrawing groups substituted bridged bicyclic systems can optimize both steric hindrance^20,21^ and electronic effects^30,31^ to prevent the formation of the TICT state, thereby enhancing the brightness and photostability; 2. Following Bredt’s rule where a carbon double bond cannot occur at the branching position of the bridgeheads^38^, the radical cation and iminium intermediates will be energetically unfavored during photo-oxidation, inhibiting this major photobluing-photobleaching pathway; 3. Two types of auxochromes contain different hydrogen-bond acceptors will render the dyes customizable hydrophilicity and membrane permeability for various applications. Before the chemical synthesis and characterization, we first examined these two auxochromes *in silico* on rhodamine scaffold, which represents the most popular D-A type fluorophores in biological imaging. First, we characterized the steric hindrance effect of the auxochromes by the value of the C_1_-N-C_n_ bond angle, where the steric congestion of two [3.2.1] bridged bicyclic rings is lower than that of *N, N*-dimethyl auxochromes or deuterated pyrrolidine yet is higher than that of azatidine or azabicyclo[2.2.1]heptane moieties (Fig. 1c). Next, we evaluated their electron richness donating effects by calculating their vertical ionization energies (VIEs) (Fig. 1c). The results indicated that two [3.2.1] bridged bicyclic rings have higher VIEs than other moieties, implying that they are less electron-rich and harder to undergo potential single electron oxidation. Finally, we calculated the potential energy surfaces of **TMR/JF549/JFX554/[2.2.1]R** and **BD528/BD555** in water as a function of auxochrome rotation angle (Fig. 1c and S1). Our results suggested that the formation respectively) than that in **TMR/JF549/JFX554/[2.2.1]R** (0.18 eV, 0.10 eV, 0.20 eV and 0.35 eV respectively). In addition, the TICT state of **BD528/BD555** does not form a minimum on the excited state potential energy surface, resulting in a positive driving energy (*E*_DE_) which means that these two fluorophores are energetically prohibited from transitioning into their TICT state^39^, rendering the fluorescence emission their main pathway which translates into high fluorescence quantum yields.

### Photophysical properties of BD dyes in aqueous buffer

Encouraged by the calculation results, we installed the above auxochromes into different scaffolds including coumarin, rhodamine, carbon-rhodamine, oxazine, and silicon-rhodamine classes, forming a palette of **BD** dyes span the UV and visible range with different photophysical properties (Fig. 1d). **BD** dyes are generally synthesized in a modular manner. Starting from reported aryl triflates intermediates, we used the Buchwald-Hartwig cross-coupling approach to install the corresponding azabicyclo [3.2.1] octane in coumarin, rhodamine, carbon-rhodamine and oxazine fluorophores^35^ (Scheme S1 a-c). For silicon-rhodamine derivatives, the bicycles were introduced early in the synthetic route via the Buchwald-Hartwig reaction, followed by Friedel-Crafts type cyclization to give the silicon-rhodamine chromophore^40^ (Scheme S1 d-f).

We then measured the photophysical properties of these **BD** dyes against their *N, N*-dialkyl prototypes in aqueous buffer (Table 1 and Fig. S2). On coumarin scaffolds, **BD350** and **BD356** exhibited significantly increased fluorescence quantum yield (13.7-fold and 10.3-fold respectively), lifetime, and blue-shifted absorption spectrum compared to **coumarin 460**. On rhodamine scaffolds, **BD528** bearing SO_2_ substituted azabicyclo[3.2.1] octane auxochromes exhibit quantum yields close to the theoretical limit (98%), blue-shifted spectra (*λ*_Abs_/*λ*_Em_= 528/554 nm), and reduced equilibrium constant (*K*_L-Z_=0.090); **BD555** containing O substituted azabicyclo[3.2.1] octane auxochromes exhibited 1.8-fold increases in quantum yield and bathochromic shift in absorption maximum and emission maximum (*λ*_Abs_/*λ*_Em_= 555/587 nm). Compared to **[2.2.1]Rhod** and **TMR**, **BD555** and **BD528** also exhibited significantly improved water solubility (Fig. S3). The bridge auxochromes also showed a similar trend on the carbon-rhodamine scaffold. Notably, the lower apparent extinction coefficients and equilibrium constant *K*_L-Z_ of compounds **BD586** and **BD615** indicated a more pronounced trend toward their non-fluorescent lactone form in aqueous buffer. SO_2_ substituted azabicyclo[3.2.1] octane moiety on **BD623** for oxazine and **BD634** for monomethyl-substituted silicon-rhodamine scaffolds also exhibited markedly high fluorescence quantum yield (36% and 60%, respectively). Furthermore, O substituted azabicyclo[3.2.1] octane moiety on **BD655** for carboxyl-substituted silicon-rhodamine scaffolds showed a bathochromic shift in absorption and emission spectrum (*λ*_Abs_/*λ*_Em_= 655/680 nm) and smaller *K*_L-Z_ (0.0012 vs 0.0036) which suggested that it could be a better far-red fluorogenic labeling for protein tags. Overall, the bridge auxochromes generally elevate the fluorescence quantum yield of commonly used chromophores to a remarkably high level.

**Table 1.**
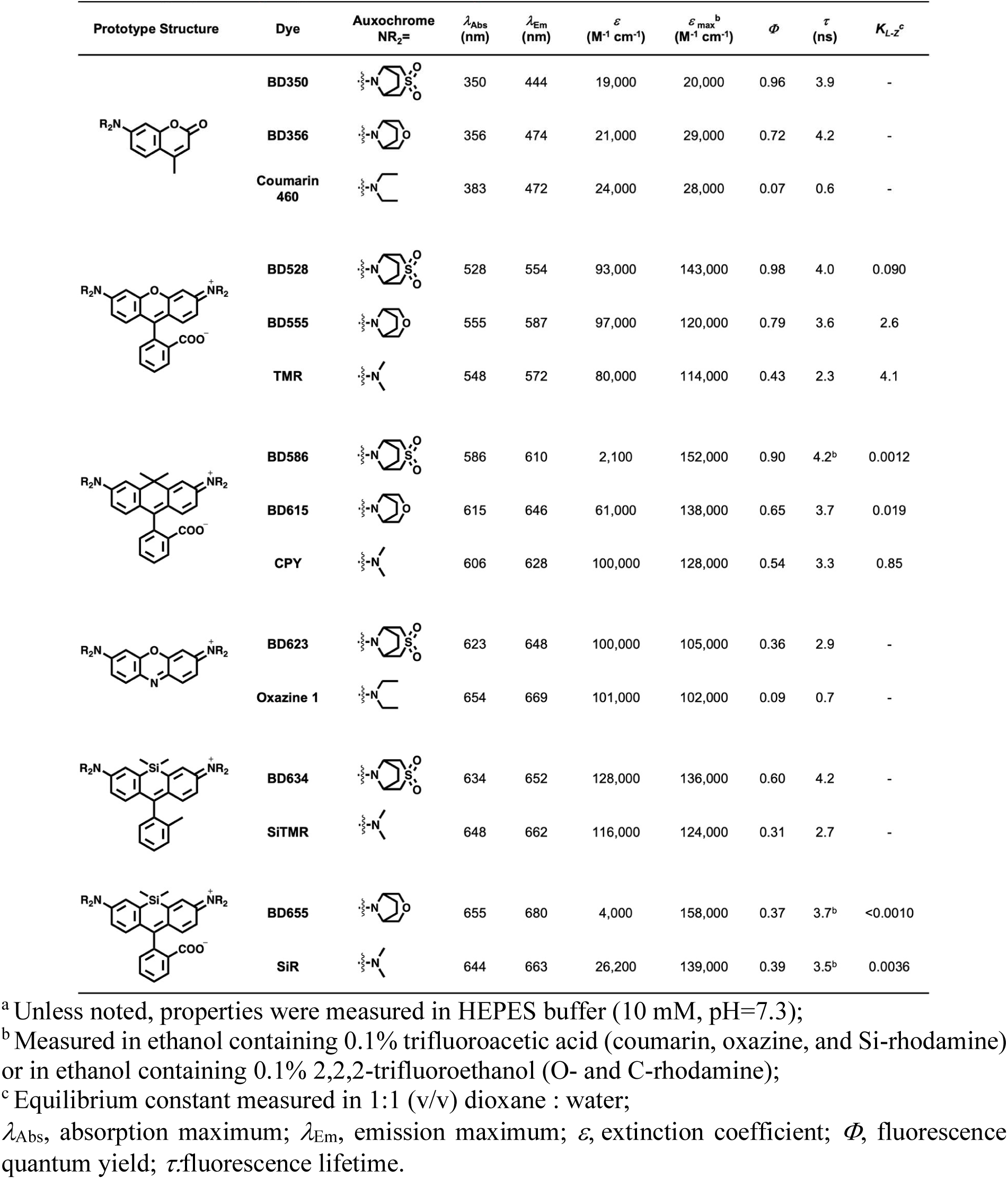
Photophysical properties of fluorophores^a^.

### Photostability of BD dyes in aqueous buffer

The primary known photobleaching pathway of rhodamine dyes is oxidative dealkylation at the N atom. Under irradiation and in oxygen-containing environments, the oxidative dealkylation of auxochromes successively generates radical cation, α-aminoalkyl radical, and iminium cation, which eventually hydrolyzed to give an aldehyde and a dealkylated dye (Fig. 2a). Each step of the dealkylation process will result in a blue shift of 10-15 nm in the absorption spectrum. These photodegradation reactions not only reduce the total photon budget but also cause spectral shift into shorter wavelength channels, complicating data analysis in multiplexed imaging^26,27,41^.

**Fig. 2.**
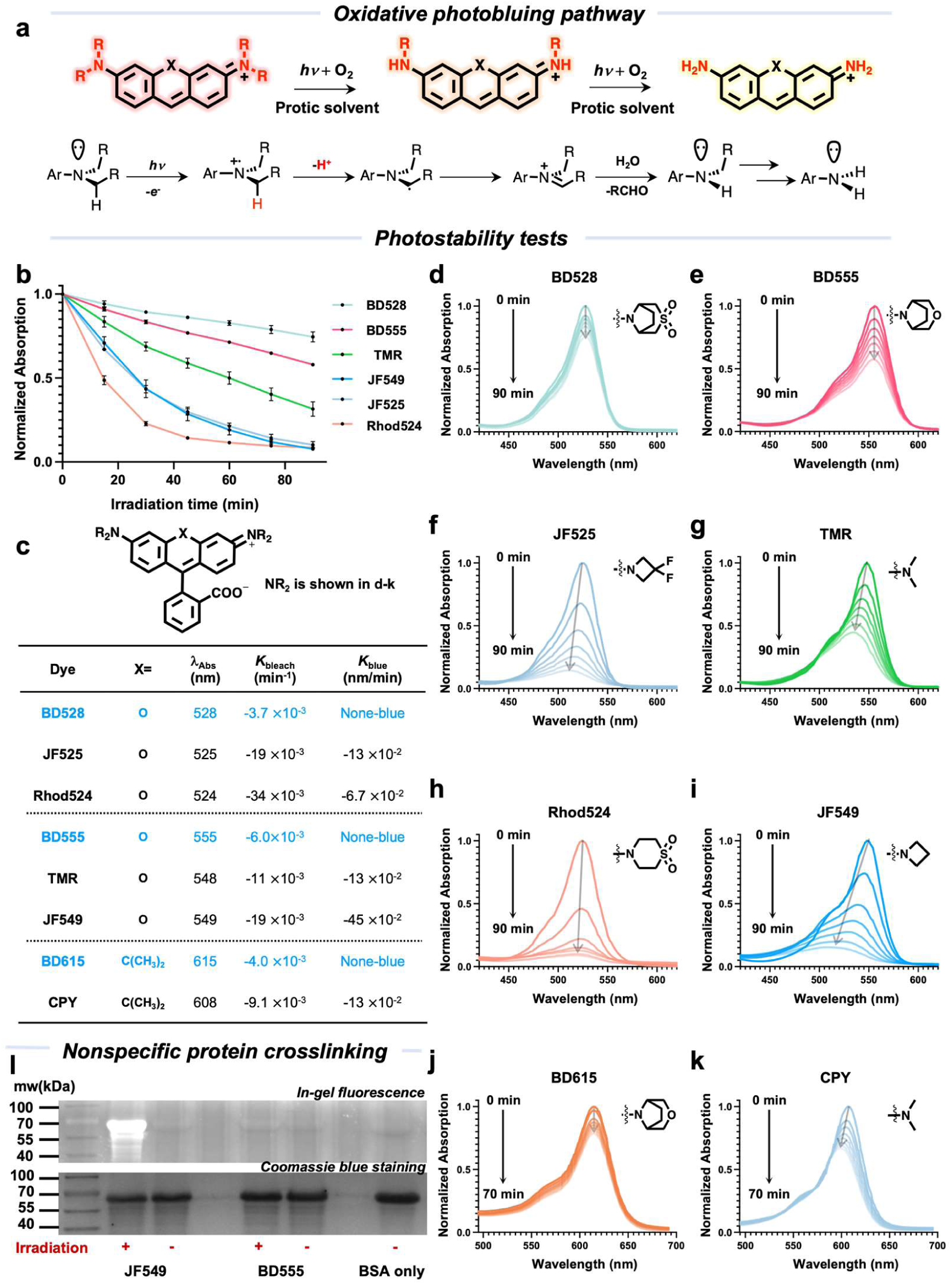
Photostability and photobluing properties of BD and benchmarking dyes. (a) General oxidative photobluing pathway for fluorescent dyes with (di)alkylamino auxochromic groups; (b) Absorptions at λ_max_ plotted against time.**BD528**, **BD555,** and various fluorophores (10 μM) with similar spectral properties were irradiated with an LED lamp (0.5 W/cm^2^, 520-530 nm) in HEPES buffer (10 mM, pH 7.3) (*n*=3, error bar show mean± S.D.); (c) Photobleaching and photobluing rates of selected fluorophores; (d) to (i) The absorption of dyes over time demonstrates the photobleaching of photooxidation-resistant, nonbluing fluorophores **BD528**, **BD555** (d-e) in comparison with photooxidation-prone fluorophores **TMR**, **JF549**, **JF525** and **Rhod524** (f-i). (j) to (k) The absorption of dyes over time demonstrates the photobleaching of photooxidation-resistant, nonbluing fluorophore **BD615** (j) in comparison with photooxidation-prone fluorophore **CPY** (k) under irradiation with an LED lamp (0.5 W/cm2, 590-600 nm) in HEPES buffer (10 mM, pH 7.3). (l) Photocrosslinking is a downstream side reaction of photooxidation. Solutions of fluorophores (10 μM in 0.1% (v/v) DMSO–PBS, pH 7.4, air-saturated) were irradiated in presence of BSA (1 mg/mL 25 °C, 30 min) with an LED lamp (0.5 W/cm^2^, 520-530 nm). The incubated samples were analyzed using in-gel fluorescence and coomassie blue staining under denaturing conditions (SDS-PAGE).

Thanks to the Bredt’s rule where double bonds are not favored at the bridgehead carbon, the bridged bicyclic auxochrome should prohibit the formation of the radical cation and iminium intermediate, blocking the dealkylative photobleaching and photobluing pathways. The photobleaching process of **BD** dyes as well as other state-of-the-art fluorophores in aqueous buffer was monitored using UV-Vis spectroscopy (Fig. 2b). The bridge bicycle **BD528** and **BD555** show the best photostability and spectral stability, with distinctive non-photobluing features (Fig. 2d and 2e). In contrast, the other selected rhodamine dyes undergo varying degrees of photobluing (Fig. 2f to 2i). The photobleaching rate of **BD 528** in aqueous solutions is slower than **JF525** and **Rhod524** (5.1-folds and 9.2-folds respectively) (Fig. 2c). The same strong effect on photostability also occurs between **BD555** to **TMR** and **JF549** (3.0-folds and 5.1-folds respectively) (Fig. 2c). Notably, electron-withdrawing groups significantly reduce the photobluing rate (**JF525** and **Rhod524** vs **JF549**), but this trend appears to be independent of the photobleaching process. On the red-shifted carbon-rhodamine scaffold, the bridge bicycle **BD615** also shows improved photostability and spectral-stability compared to **CPY** (Fig. 2c, 2j&2k). In addition to photostability, it is worth noting that the reactive aldehyde species derived from the oxidative photobluing byproducts will contribute to cross-linking reactivity with proteins. For a test *in vitro*, we irradiated bluing (**JF549**) and nonbluing (**BD555**) fluorophores in air-saturated PBS buffer with bovine serum albumin (BSA) for 30 min. SDS-PAGE analysis showed the formation of covalent dye-BSA conjugates with **JF549**, while **BD555** did not give this side-reaction (Fig. 2l). These experiments underscore the advantages of non-photoblue dyes in labeling biomacromolecules.

### BD derivatives in immunofluorescence imaging

Encouraged by the excellent photophysical properties of **BD** dyes, we tested their application in immunofluorescence. Other than brightness and photostability, a key factor for immunofluorescence is to achieve a high degree of labeling on secondary antibodies, which requires superb solubility and minimal aggregation in aqueous environments^42,43^. Traditionally, such solubilization was achieved by sulfonation. We envisioned that the **BD** dyes bearing sulfones in the bridged bicycles are ideal candidates, as the neutral sulfones have rendered them high solubility without introducing excessive charges which may hamper protein functions. To this end, the *ortho*-carboxyl groups on the pendant ring of **BD** rhodamines were functionalized with 4-(methylamino) butanoic acid as coupling handles (Fig. 3a, Scheme S2). Notably, such functionalization gives a bathochromic shift in the Abs/Em spectrum (~ 15 nm) and, at the same time locks the chromophore in the open form, as characterized by an increase in extinction coefficient (Fig. 3b and Fig. S4). In photostability tests (Fig. 3c and S5), **BD Yellow** showed 2.7-folds higher photostability than **ATTO 532**, with distinctive spectral stability (i.e. no photobluing) (Figure 3c and Figure S5 c-d). **BD Orange** and **ATTO 594** exhibited similar photostability (Figure 3d and Figure S5 e-f). **Atto 647N**, one of the most popular dye for single-molecule and super-resolution microscopy, showed significant blue shift after irradiation while **BD Red** was a none-photobluing dye with 8.0-folds higher photostability (Fig. 3e, and Fig. S5 g-h).

**Fig. 3.**
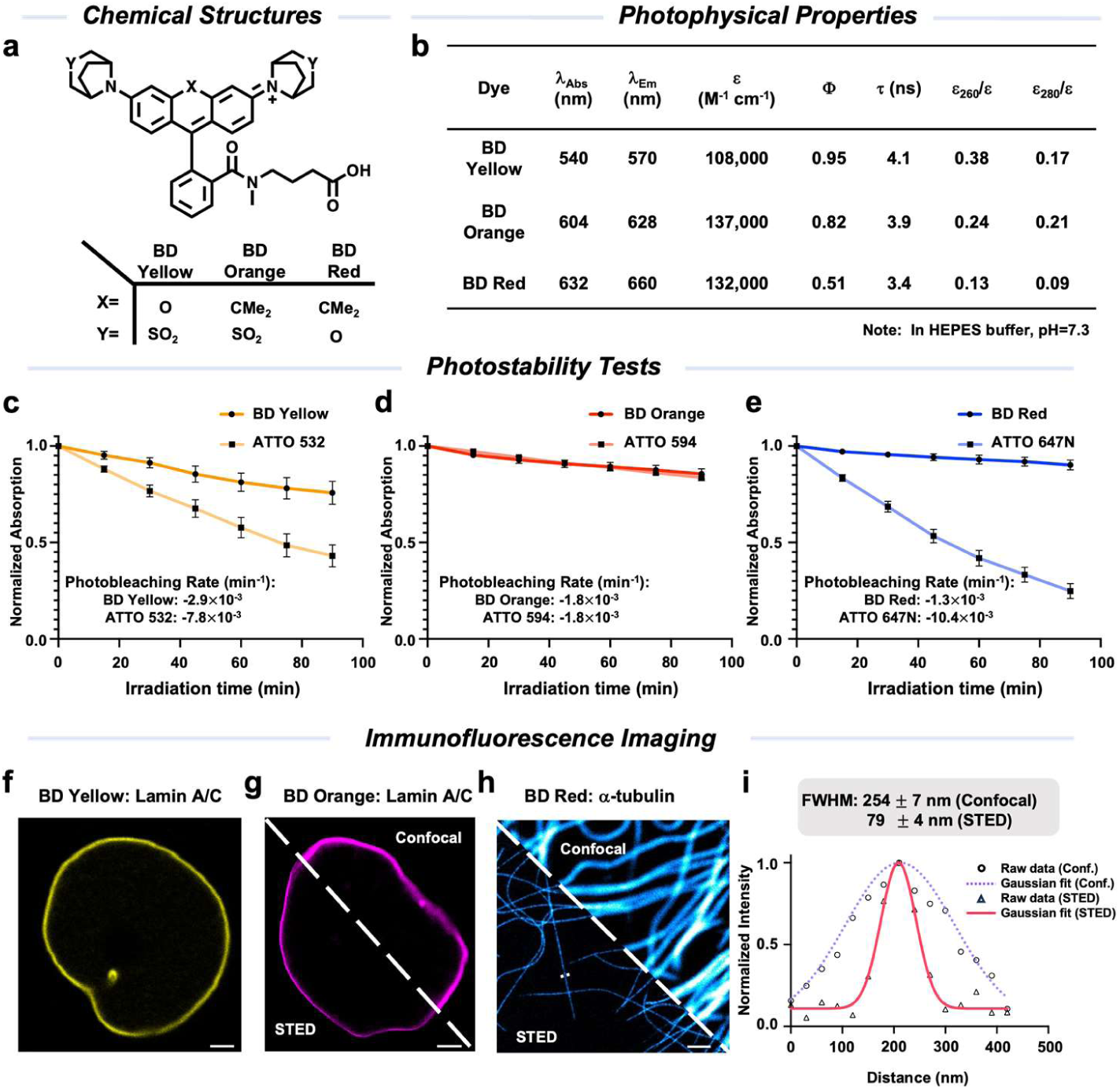
BD derivatives for antibody bioconjugation and immunofluorescence imaging. (a) Chemical structures of **BD** derivatives for antibody bioconjugation; (b) Photophysical properties of **BD** derivatives; (c) to (e) Photobleaching curves of BD derivatives in HEPES buffer (10 mM, pH=7.3), (*n*=3, error bar show mean± S.D.). (c) Absorption at λ_max_ of **BD Yellow** and **ATTO 532** (10 μM) were plotted as a function of irradiation time with an LED lamp (0.5 W/cm^2^) at 520-530 nm; (d) Absorption at λ_max_ of **BD Orange** and **ATTO 594** (10 μM) were plotted as a function of irradiation time with an LED lamp (0.5 W/cm^2^) at 590-600 nm; (e) Absorption at λ_max_ of **BD Red** and **ATTO 647N** (10 μM) were plotted as a function of irradiation time with an LED lamp (0.5 W/cm^2^) at 620-630 nm; (f) Confocal images of lamin A/C structures in fixed-HeLa cells labeled by indirect immunofluorescence with a secondary antibody bearing **BD Yellow** (DOL=3.27), 561 nm Ex./575-625 nm Em., scale bar=2 μm; (g) Confocal (right) and STED (left) images of lamin A/C structures in fixed-HeLa cells labeled by indirect immunofluorescence with a secondary antibody bearing **BD Orange** (DOL=2.63), 640 nm Ex./650-700 nm Em., STED at 775 nm, scale bar=2 μm; (h) Confocal (right) and STED (left) images of α-tubulin structures in fixed-HeLa cells labeled by indirect immunofluorescence with a secondary antibody bearing **BD Red** (DOL=2.42), 640 nm Ex./650-700 nm Em., STED at 775 nm (~140 mW), scale bar=1 μm; (i) Line-scan profile of fluorescence intensity at the dotted line in (h) and comparison of the FWHM achieved under confocal and STED conditions (n ≥ 15 filaments from 3 samples, mean S.D.).

Then, the amino-reactive *N*-hydroxysuccinimide (NHS) esters of **BD Yellow/Orange/Red** were prepared and reacted with goat anti-mouse secondary antibodies. The degree-of-labelings are 3.27, 2.63, and 2.42, respectively, showcasing their remarkable solubilities. **BriDyes** still outperform their **ATTO** counterparts in photostability when conjugated to antibodies (Fig. S6). Immunofluorescence imaging of Lamin A/C and alpha-tubulin using **BD Yellow**, **BD Orange**, and **BD Red** gives strong signals in confocal mode (Fig. 3f-3h). Thanks to their photostability, **BD Orange** and **BD Red** are compatible with stimulated emission depletion microscopy (STED) imaging using a 775 nm depletion laser (Fig. 3g & 3h). Single microtubules can be resolved with a resolution under 80 nm (~ 140 mW depletion laser, Fig. 3h & 3i), and its ultimate resolution can reach down to 30 nm under ~ 200 mW depletion laser (Fig. S7). Overall, bridged bicyclic rhodamines are new additions to the toolkit for STED immunofluorescence imaging.

### BD dyes synergize with HaloTag *in vitro* and *in cellulo*

To evaluate the compatibility of BD derivatives in live cell imaging, we coupled HaloTag ligands with **BD555, BD615,** and **BD655** as they exhibited moderate hydrophilicity and suitable *K*_L-Z_ ratios for crossing cell membrane (Fig. 4a and Scheme S3). Upon binding with purified HaloTag7 protein, three **BD_HTL_** showed a marked increase in both absorption and emission spectrum (Fig. 4b and Fig. S8), indicating a profound influence of protein environment on both the lactone-zwitterion equilibrium and the fluorescence properties. In contrast to their *N, N*-dimethyl or azetidine-containing analogs, these **BD_HTL_** derivatives showed a bathochromic shift in absorption spectra around 8-13 nm and emission spectra around 16-22 nm upon binding to HaloTag. So we renamed these HaloTag ligands based on their maximum absorption after binding to protein (for example **BD555** to **BD566_HTL_** and so on). **BD566_HTL_** and **BD626_HTL_** are among the brightest labels in their respective 561 and 640 channels, with extinction coefficients higher than 100,000 M^-1^cm^-1^ and quantum yields higher than 0.80 after binding to HaloTag7. **BD666_HTL_** showed 40× increase in absorption and 70× increase in emission (Fig. 4j and Fig. S8e and f) upon HaloTag binding, which is more fluorogenic than the widely used far-red dye **SiR_HTL_**^44^(4.8× increase in absorption and 15× increase in emission).

**Fig. 4.**
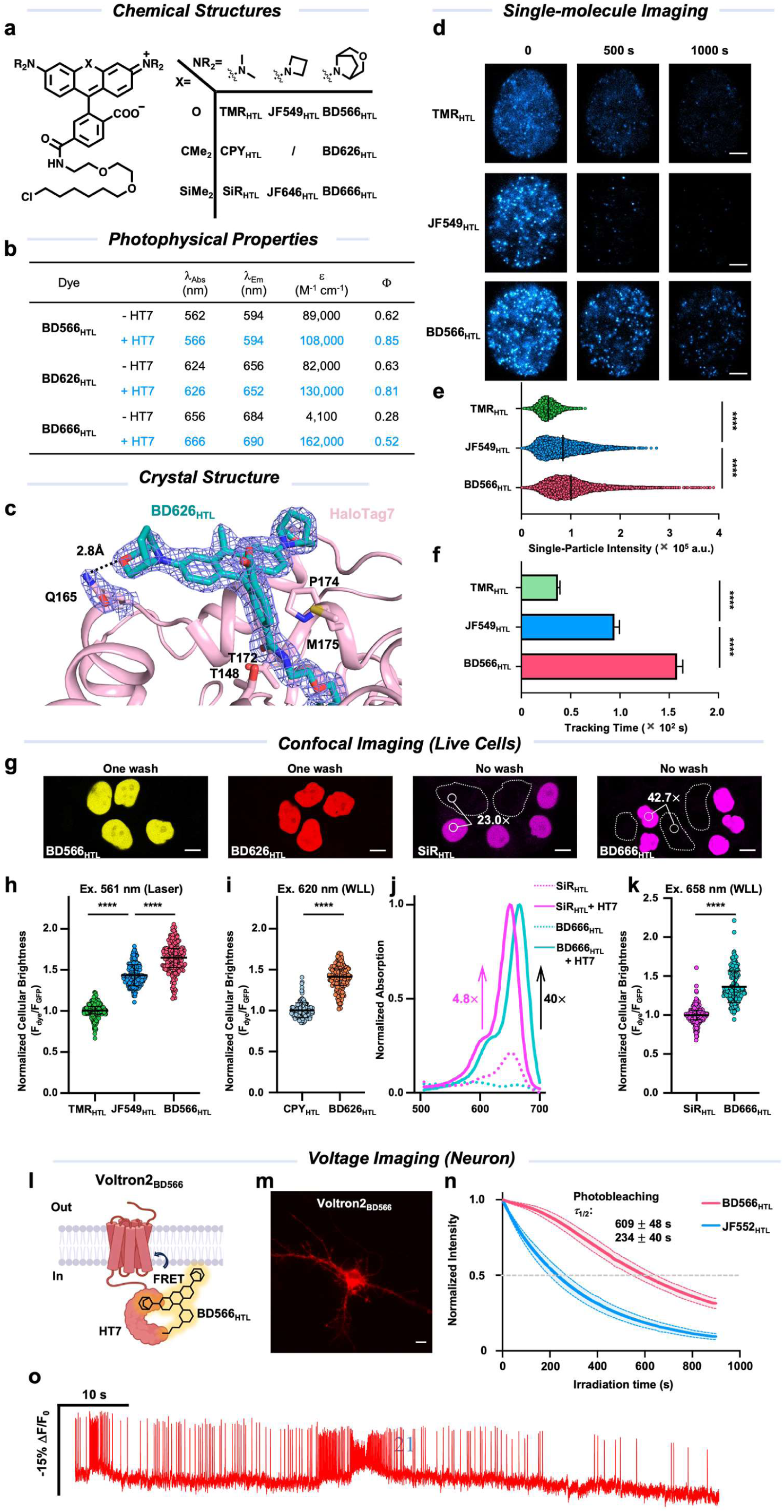
BD dyes with O-substituted azabicyclo[3.2.1] octane auxochromes are privileged HaloTag ligands for labeling and imaging cellular targets. (a) Chemical structures of BD derivatives and their dimethyl and azetidine counterparts with HaloTag ligands; (b) Photophysical properties of fluorescent HaloTag ligands; (c) A close-up crystal structure of **BD626_HTL_** in the pocket of HaloTag7 highlighting its polar interaction with Q165; 2*F_σ_*-*F_c_* omit electron density in the BD626HTL-HaloTag7 complex crystal structure contoured at 1.0*σ*; (d) Indicated time point of continuous time-lapse single-molecule imaging of fixed U2OS cells stably expressing H2B-HaloTag7 labeled with **TMR_HTL_**, **JF549_HTL_** and **BD566_HTL_** (2.5 pM, 30 min, 37°C, 2× wash), scale bar= 5 μm; (e) Fluorescence intensity per particle per frame for three fluorescent ligands: For **TMR_HTL_**, n= 4018; **JF549_HTL_**,n= 7464; **BD566_HTL_** n= 12067; p****< 0.0001, t-test; error bar show mean± S.E.M.; (f) Tracking time of tested dyes: For **TMR_HTL_**, n= 1924; **JF549_HTL_**,n= 1054; **BD566_HTL_** n= 1909; p****< 0.0001, t-test; error bar show mean± S.E.M.; (g) Live-cell confocal images of HeLa cells stably expressing H2B-HaloTag7-GFP labeled with **BD566_HTL_** (200 nM, 2 h, 37°C, one wash), **BD626_HTL_** (200 nM, 1 h, 37°C, one wash) or **SiR_HTL_** and **BD666_HTL_** (200 nM, 1 h, 37°C, no wash), mean of signal-over-background ratios (S/B) are indicated in figures (n≥100 areas in two independent experiments), scale bar= 10 μm; (h-i & k) Comparison of apparent cellular brightness obtained for different HaloTag ligands in 2 independent experiments. The fluorescence intensity from single nuclei was normalized by the cytosolic GFP fluorescence intensity of the same cell (expression control) to assess the cellular brightness, (error bar shows mean ± S.D.); For **TMR_HTL_**, n=189 cells, **JF549_HTL_**, n=194 cells, **BD566_HTL_**, n=170 cells; p****< 0.0001, t-test; For **CPY_HTL_**, *n*=187 cells, **BD626_HTL_**, *n*=175 cells; *p*\*\*\*\*< 0.0001, t-test; For **SiR_HTL_**, *n*=176 cells, for **BD666_HTL_**, *n*=191 cells; *p*\*\*\*\*< 0.0001, t-test. (j).Absorbance of **BD666_HTL_** and **SiR_HTL_** in the presence (+HT7) or absence (−HT7) of excess HaloTag7 protein (*n*=2). (l) A cartoon of chemigenetic voltage sensor Voltron2 labeled with a **BD566_HTL_**; (m) A representative wide-field image of a cultured rat hippocampal neuron expressing Voltron2 and labeled with **BD566_HTL_**, scale bar=10 μm; (n) Normalized fluorescence decay curves of Voltron2 labeled with **BD566_HTL_** and **JF552_HTL_** during time-lapse confocal imaging on fixed neurons, *n*=4 in two independent experiments; error bar show mean± S.D.; (o) Fluorescence trace of voltage imaging recorded from a neuron prepared in (m).

To better understand the interplay between **BD_HTL_** dyes and HaloTag protein, a single crystal of **BD626_HTL_** at the pocket of HaloTag7 was obtained and its structure was solved at 1.7 Å resolution (PDB entry 9JHA). Compared to its *N, N*-dimethyl analog **CPY_HTL_**-bound HT7 (PDB entry 6Y7B, at 3.1 Å resolution)^45,46^ the pockets in which the dyes reside are generally similar (Fig. S9a). In both **CPY_HTL_**- and **BD626_HTL_**-bound HT7, T172, and T148 engage with the oxygen and the nitrogen of the amide bond linking HaloTag ligands and fluorophore (Fig. S9b and 8c). Yet the positions and angles of chromophores on the HT7 protein are slightly different (Fig. S9d). Notably, the distance from one of the O-substituted azabicyclo[3.2.1] octane on **BD626_HTL_** to the amide side chain of Q165 is only 2.8 Å (Fig. 4c), indicating an active polar interaction (2.5 Å ~3.5 Å) which is absent in the **CPY_HTL_** counterpart. This polar interaction suggests that the hydrogen-bond acceptor on the auxochrome of fluorophore brings up new interaction sites with protein residues, giving new biochemical space for further protein engineering for advanced applications such as chemigenetic hybrid sensors^47,48^.

Next, we evaluated the performance of **BD_HTL_** at the single-molecule level in a cellular context. High dilution of HaloTag ligands (2.5 pM) was applied to live U2OS cells stably expressing H2B-HaloTag7 to ensure sparse labeling for single-molecule detection, followed by paraformaldehyde fixation. Continuous imaging of single molecules was acquired under total internal reflection fluorescence microscopy with Highly inclined thin illumination (HiLo) (Fig. 4d and Movie S1). **BD566_HTL_** labeling exhibited highest localization counts (Fig. S11a), featuring a 19% and 85% increase of single-molecule brightness compared to **JF549_HTL_** and **TMR_HTL_** labeling, respectively (Fig. 4e). Moreover, **BD566_HTL_** exhibited the best photostability among the three in single-molecule experiments (Fig. S11b), consistent with bulk *in vitro* measurement (Figure 2d). Thanks to the highest brightness and photostability of **BD566_HTL_**, the track lengths of single H2B molecules labeled by **BD566_HTL_** (158 ± 6 s) were 1.7 and 4.3 folds longer than those labeled by **JF549_HTL_** (95 ± 5 s) and **TMR_HTL_** (37 ± 2 s), respectively. These data demonstrated the outstanding performance of **BD566_HTL_** in cellular single-molecule imaging experiments.

We further assessed the performance of **BD_HTL_** on bulk imaging experiments using HeLa cells stably expressing a nuclear-localized H2B-HaloTag7-GFP fusion protein (Fig. 4g). **BD566_HTL_** showed 14% and 64% higher apparent brightness than **JF549_HTL_** and **TMR_HTL_** after 2 h incubation under 200 nmol concentration with living HeLa cells (Fig. 4h). The labeling kinetics of nuclear HaloTag7 with **BD566_HTL_** was about 50% slower than **TMR_HTL_** due to its slightly lower permeability rendered by higher hydrophilicity (signal saturate within 30 min vs 20 min, Fig. S10a). Due to the difference between **CPY_HTL_** and **BD626_HTL_** in excitation spectra, a white light laser (WLL) is employed to excite them at 620 nm for a comparable excitation efficiency. **BD626_HTL_** showed 40% higher brightness than **CPY_HTL_** under this condition (Fig. 4i). Similarly, **BD666_HTL_** exhibited better signal to background ratio (S/B) than **SiR_HTL_** (42.7 v.s. 23.0) ^44^under no wash condition and 36% higher brightness under 658 nm excitation laser at which channel the excitation efficiencies are the same (Fig. 4k). There is no significant difference in cell permeability between **CPY_HTL_** and **BD626_HTL_** (Fig. S10b), both fluorescence signals saturate within 2.5 min, as well as **SiR_HTL_** and **BD666_HTL_** (Fig. S10c)

When imaged at single-molecule level using a 642-nm excitation laser, **BD626_HTL_** labeled single H2B molecules showed a 1.30 and 2.18 fold of brightness than **SiR_HTL_** and **BD666_HTL_** respectively (Fig. S12a). In bulk experiments using a 640-nm excitation laser, **BD626_HTL_** showed 352% higher brightness than **CPY_HTL_** (Fig. S12b) yet **BD666_HTL_** showed 49% lower brightness than **SiR_HTL_** (Fig. S12c). The differences in the above data are likely caused by the mismatch in excitation efficiency and cannot fairly compare the performance of **BD_HTL_** with other dyes. However, these data are valuable reference due to the wide use of 640/642-nm lasers on fluorescence microscopy.

Encouraged by the outstanding performance of **BD_HTL_** in structural imaging in live cells, we further applied **BD566_HTL_** to functional voltage imaging in cultured rat hippocampal neurons. We chose the chemigenetic hybrid voltage sensor, Voltron2 as a model, which contains a self-labeling HaloTag protein fused to the C-terminus of a voltage sensitive rhodopsin (Fig. 4l) ^50^. Due to the requirement of high-speed camera frame rate for voltage imaging, which typically exceeds 500 Hz, Voltron2-based sensors are very demanding in brightness and photostability for time-lapse data acquisition. Hippocampal neurons expressing Voltron2 can be readily labeled with **BD566_HTL_,** giving a bright signal at the plasma membrane (Fig. 4m). Voltron2_BD566_ is capable of recording spontaneous neuronal action potentials with a sensitivity of approximately 11.32± 0.69% (Fig. 4o). And there is no significant difference in voltage sensitivities of **BD566_HTL_** and **JF552_HTL_** on Voltron2 (−23.8% v.s. −25.3%) in HEK293T cells (Fig. S13). Compared to the spectrally similar Voltron2_JF552_, Voltron2_BD566_ is 2.6 times more photostable as measured by half-bleaching time in fixed neurons (τ_1/2_:609 s v.s. 234 s) (Figure 4n). Together, the above voltage imaging data demonstrate that the brightness and photostability of Voltron2_BD566_ would make it an ideal sensor for the time-lapse recording of neuronal activity.

### BD_HTL_ are powerful tools for general imaging in cells, plants, and animals

Under confocal imaging, **BD566_HTL_** and **BD626_HTL_** exhibited high brightness, ideal permeability, and high photo- and spectral-stability, making them privileged candidates for super-resolution imaging with boosted illumination schemes. In live-cell time-lapse structured illumination microscopy (SIM) imaging, the signal of **BD566_HTL_** labeled TOMM20-HaloTag7 remain more than 75% fluorescence after 100 frames (Fig. S15 a-b). In live-cell STED microscopy, **BD626_HTL_** labeled microtubule-binding protein Cep41-HaloTag7 (Fig. 5a) showed markedly nanoscopic resolutions (41 ± 2 nm) under 775 nm depletion laser (~160 mW). The resolution is comparable to **SiR_HTL_** (50 ± 5 nm, Fig. S15). Live-cell STED time-lapse imaging further captured the dynamics of mitochondria (Fig. 5d). For **BD626_HTL_** in live-cell time-lapse STED imaging, the frame numbers with >50% original intensity (τ_1/2_) was around 2 times higher than **SiR_HTL_** and 3 times higher than **JF646_HTL_** (Fig. 5d&e). Furthermore, **BD626_HTL_** enabled 3D STED imaging which was prohibitively challenging due to the repeated photobleaching during z-stacking. Practically, **BD626_HTL_** can offer > 25 z-frames for 3D reconstruction, enabling the reconstruction of complete 3D structure of mitochondrial outer membrane (Figure 5b) or the distribution of the cytoskeleton (Cep41) throughout the entire cell (Figure 5c). **BD566_HTL_** and **BD626_HTL_** are therefore recommended reagents for live-cell time-lapse nanoscopic and 3D imaging in red and far-red channels.

**Fig. 5.**
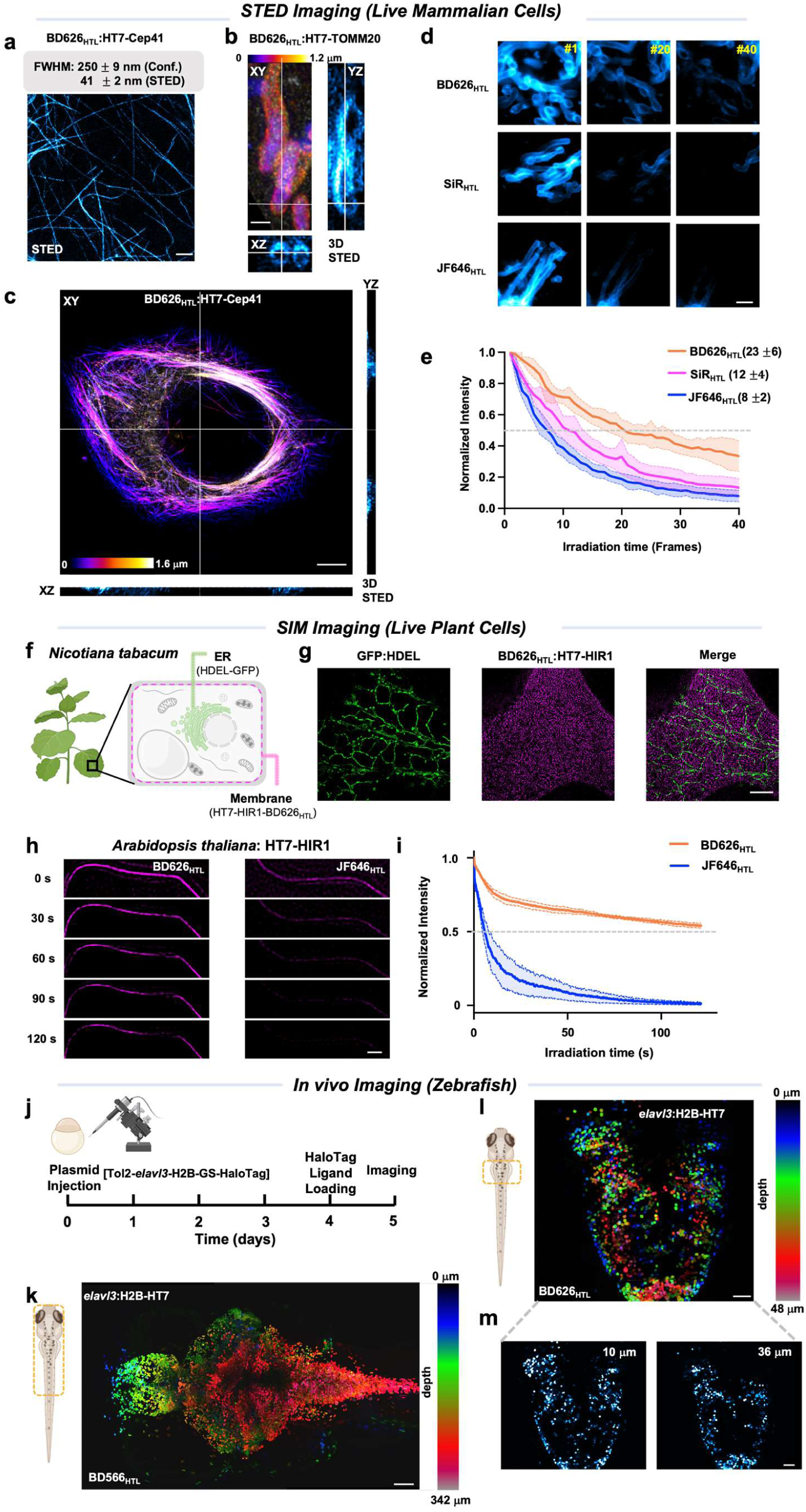
BDHTLs are versatile tools for imaging in cells, plants, and animals. (a) 2D-STED images of live HeLa cells expressing Cep41-HT7 labeled with **BD626_HTL_** (500 nM, 30 min, 37°C, one wash), and comparison of the FWHM achieved under confocal and STED (at 775 nm, ~160 mW) conditions (n ≥12 filaments from 3 samples, error bar show mean ± S.D.); scale bar= 1 μm; (b) Orthogonal views of 3D-STED images of HeLa cells expressing TOMM20-HT7 labeled with **BD626_HTL_** (500 nM, 30 min, 37°C, one wash, fixed). A volume of 3.3 × 8.5× 1.2 μm (*xyz*) was recorded using 30 z-stack images with a z-step of 40 nm; scale bar= 1 μm. (c) Orthogonal views of 3D-STED images of HeLa cells expressing Cpe41-HT7 labeled with **BD626_HTL_** (500 nM, 30 min, 37°C, one wash, fixed). An area of 48.5 × 50.0× 1.6 μm (*xyz*) was recorded using 25 z-stack images with a z-step of 65 nm; scale bar= 1 μm. (d) Time-lapse STED images of live HeLa cells expressing TOMM20-HT7 labeled with **BD626_HTL_**, **SiR_HTL_** and **JF646_HTL_** (500 nM, 30 min, 37°C, one wash). Frame numbers indicated at the top right corner (20 s/frame). Scale bar: 1 μm. (e) Normalized fluorescence decay curves of samples in (d) with half-bleaching frame numbers τ given in brackets; *n*=9 in 2 independent experiments; error bars show mean± S.D.; (f) Cartoon demonstrating the labeling strategies for ER and membrane proteins in the leaves of *N. benthamiana*; (g) Two-color TIRF-SIM images of live *N. benthamiana* cells expressing HDEL-GFP and HIR1-HT7 labeled with **BD626_HTL_** (200 nM, 1 h, 37°C, 3×wash), scale bar= 5 μm; (h) Time-lapse TIRF-SIM images of live *Arabidopsis thalianas* cells expressing HT7-HIR1 labeled with **BD626_HTL_** or **JF646_HTL_** (200 nM, 1 h, 37°C, 3×wash), scale bar=1 μm; (i) Normalized fluorescence decay curves of samples in (g), *n*=4 in 2 independent experiments; error bar show mean± S.D.; (j) Experimental scheme of larval zebrafish neuronal labeling and imaging experiments; (k-m) Z-stack confocal images of live zebrafishes expressing *elavl3*:H2B-HT7 and labeled with **BD566_HTL_** or **BD626_HTL_** (3.3 μM, 1 h, 37°C, 2 h wash). For (k), the color code image shows the whole-body 3D reconstruction from 172 individual z-section images, with z-step size 2 μm. Scale bar, 100 μm; For (l), the color code image shows the 3D reconstruction of the hindbrain from 17 individual z-section images, with z-step size 3 μm. The dashed red square on larval zebrafish outlines the imaged brain region. Scale bar, 30 μm; (m) Two z-section images at the depths of 10 and 36 μm in (l). Scale bar, 30 μm.

We then challenge whether the biocompatibility of BD dyes can enable super-resolution imaging in plants with cell walls, which are important but relatively less explored with fluorescent dyes. His-Asp-Glu-Leu (HDEL) - the endoplasmic reticulum retention signal is important to ensure the correct folding and transportation of proteins^51,52^ while membrane hypersensitive induced reaction (HIR) protein plays a significant role on plant immunity^53^. The leaves of *Nicotiana benthamiana* transiently expressed HDEL-GFP and HIR1-HaloTag7, and were incubated with **BD626_HTL_**. Two color total internal reflection SIM (TIRF-SIM) images show clear and specific signals of ER (HDEL) and membrane protein (HIR1) (Fig. 5f). We also expressed HIR1-HaloTag7 in another model plant cell: *Arabidopsis thalianas*. Live-cell time-lapse recording showed that **BD626_HTL_** (Fig. 5g, left column) is ~20 times photostable than **JF646_HTL_** (Fig. 5g, right column) at the same channel (τ_1/2_:121.2 s v.s. 6.2 s) (Fig. 5h, Movie S2).

Finally, we test whether **BD** dyes can freely diffuse into live zebrafish for *in vivo* imaging. Such assay is challenging because the dye has to be sufficiently soluble to reach a high concentration for effective staining of the animal, yet highly permeable to freely diffuse across multiple membranes. Zebrafish larvae have transparent bodies and sophisticated nervous systems, making them ideal model animals for evaluating *in vivo* performance using fluorescence imaging. To this end, *elavl3:*H2B-GS-HaloTag plasmids were injected into fertilized zebrafish eggs, and labeling and imaging were conducted at the larvae on 4 and 5 days post fertilization, respectively (Fig. 5i). **BD566_HTL_** and **BD626_HTL_** (3.3 μmol, 1 h) efficiently labeled the nuclear regions in the brain and spinal cord of live larvae under confocal microscopy. The abundant fluorescence signal allowed us to reconstruct the whole body neural system in 3D (Fig. 5j-5l, Movie S3). These whole-animal imaging experiments clearly demonstrate the *in vivo* labeling capability of **BD_HTL_**s and their potential to empower neurobiological research, from cell models to live animals, and from 3D structural imaging to long-time functional recordings.

## Discussion and conclusion

Here we establish SO_2_ and O substituted azabicyclo[3.2.1] octane auxochromes as general molecular motifs to systematically upgrade fluorophores towards high brightness, supreme photo-spectral stability, and tailored biochemical properties. From a chemistry perspective, bridged bicycle-strengthened fluorophores represent a novel, modular, and integrated solution for probe design. The compact motif achieves 5 desired properties all at once: a) suppressing the formation of TICT states from both a steric hindrance and electronic effects, giving ideal fluorescence quantum yields, b) blocking the photooxidation pathway to form blue shifted compounds, completely eliminating spectral artifacts in advanced imaging, c) fine-tuning the water solubility, cell-permeability, *K*_L-Z_ values and spectrum of fluorophores by introducing different hydrogen-bond acceptor into azabicyclo[3.2.1] octane system, bypassing traditional modifications such as sulfonation, d) pre-installing hydrogen-bond acceptors which interact with HaloTag and future engineered protein machinery, and e) streamling synthetic chemistry for facile preparations. We envision this class of auxochromes would open new chemical space for fluorophore engineering^54,55^.

From the perspective of bioimaging applications, the **BD** dye palette enables applications spanning antibody conjugation and immunofluorescence, live mammalian/plant cell labeling, chemigenetic sensors, and zebrafish labeling *in vivo*. Their superb optical properties are well suited for single-molecule imaging and time-lapse super-resolution (SIM/STED) imaging, with rigorous neck-to-neck comparisons with state-of-the-art fluorophores that are widely used. With additional tuning and functionalization methods to further optimize the BD dye palette, such as fine-tuning the spectra^22^, balancing the lactone-zwitterion equilibrium^56,57^, and alleviating phototoxicity^58^, we regard these bridged-bicyclic fluorophores would serve as infrastructures for advanced imaging applications such as multiplexed single-molecule tracking, volumetric super-resolution imaging, and chemigenetic sensing.

## Acknowledgments

This project was supported by funds from National Key R&D Program of China (2021YFF0502904 to Z.C.; 2020YFA0509502 to W.D.), the Beijing Municipal Science & Technology Commission (Project: Z221100003422013 to Z.C.), and National Natural Science Foundation of China (32170566 to W.D.). We thank Prof. Lei Chen from Peking University for the guidance in analyzing the crystal structure. We thank Prof. Jinxing Lin for the assistance on plant cell imaging. We thank the Tsinghua University Branch of China National Center for Protein Sciences (Beijing) and Tsinghua University Technology Center for Protein Research for the X-ray crystallography facility support. We thank Dr. Shilong Fan, Dr, Lele Wang and Dr. Min Li from X-ray crystallography platform of National Protein Science Facility, Tsinghua University, Beijing, China for their technical help. We thank the Metabolic Mass Spectrometry Platform of IMM, the analytical instrumentation center of Peking University, the NMR facility of the National Center for Protein Sciences at Peking University, and the National Center for Protein Sciences at Peking University.

## Data availability

All data associated with this study are presented in the article or supplementary information. The data generated during the study are also available from the corresponding author upon reasonable requese.

## Supplementary Information

Captions for Movie S1 to S3

Figs S1 to S15

Methods

Table S1

Synthesis and Characterization of New Compounds

NMR Spectra

References for Supplementary Information

