## Supplementary Information for "A Palette of Bridged Bicycle-Strengthened Fluorophores"

**Table of Contents:**

Captions for Movie S1 to S3------------------------------------------------------------------------------S2
Figs S1 to S15-----------------------------------------------------------------------------------------S3
Methods----------------------------------------------------------------------------------------------- S13
Table S1---------------------------------------------------------------------------------------------- S21
Synthesis and Characterization of New Compounds------------------------------------------------ S22
NMR Spectra------------------------------------------------------------------------------------------- S39
References for Supplementary Information--------------------------------------------------------- S63

**Captions for Movie S1 to S3**

**Movie S1** Time-lapse single-molecule images of fixed U-2 OS cells stably expressing H2B-HaloTag7 labeled with **TMR$_{HTL}$**, **JF549$_{HTL}$** and **BD566$_{HTL}$**, scale bar= 5 μm.

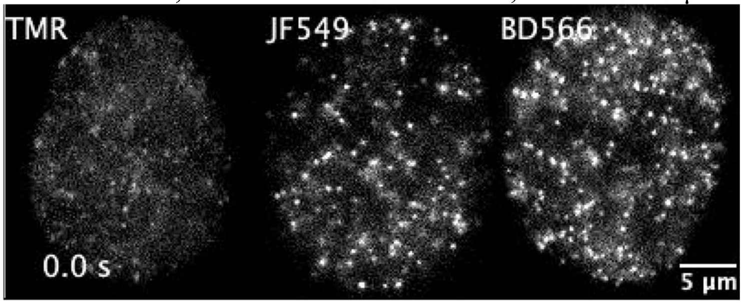


**Movie S2** Time-lapse TIRF-SIM images of live *Arabidopsis thalianas* cells expressing HIR1-HaloTag7 labeled with **BD626$_{HTL}$** or **JF646$_{HTL}$**, scale bar=1 μm.

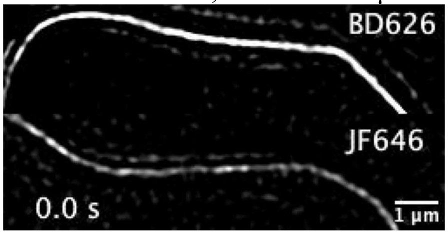


**Movie S3** Z-stack confocal images and 3D reconstruction of a live zebrafish expressing *elavl3*:H2B-HT7 and labeled with **BD566$_{HTL}$**.

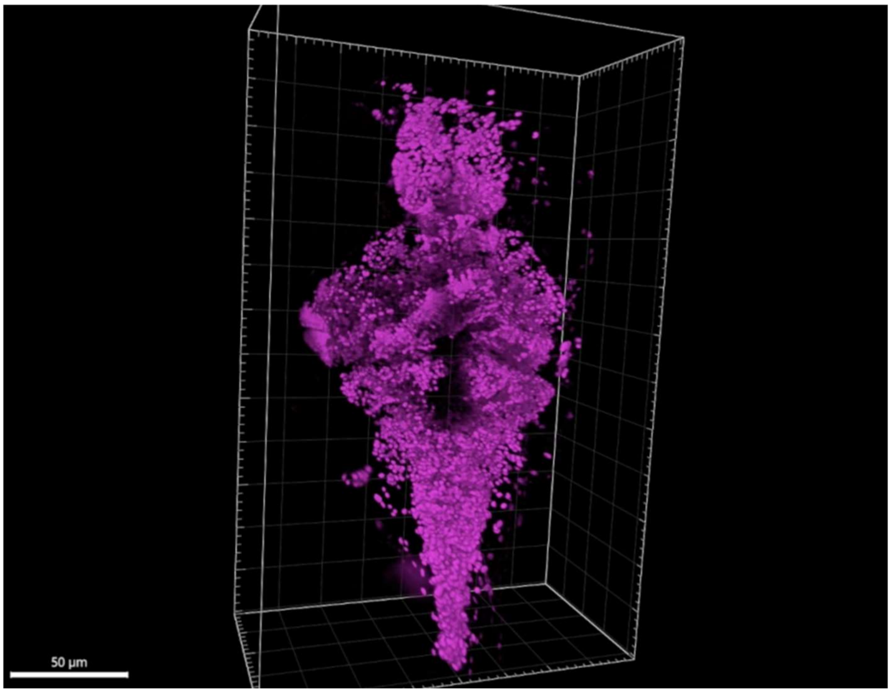

**Figs S1 to S11**

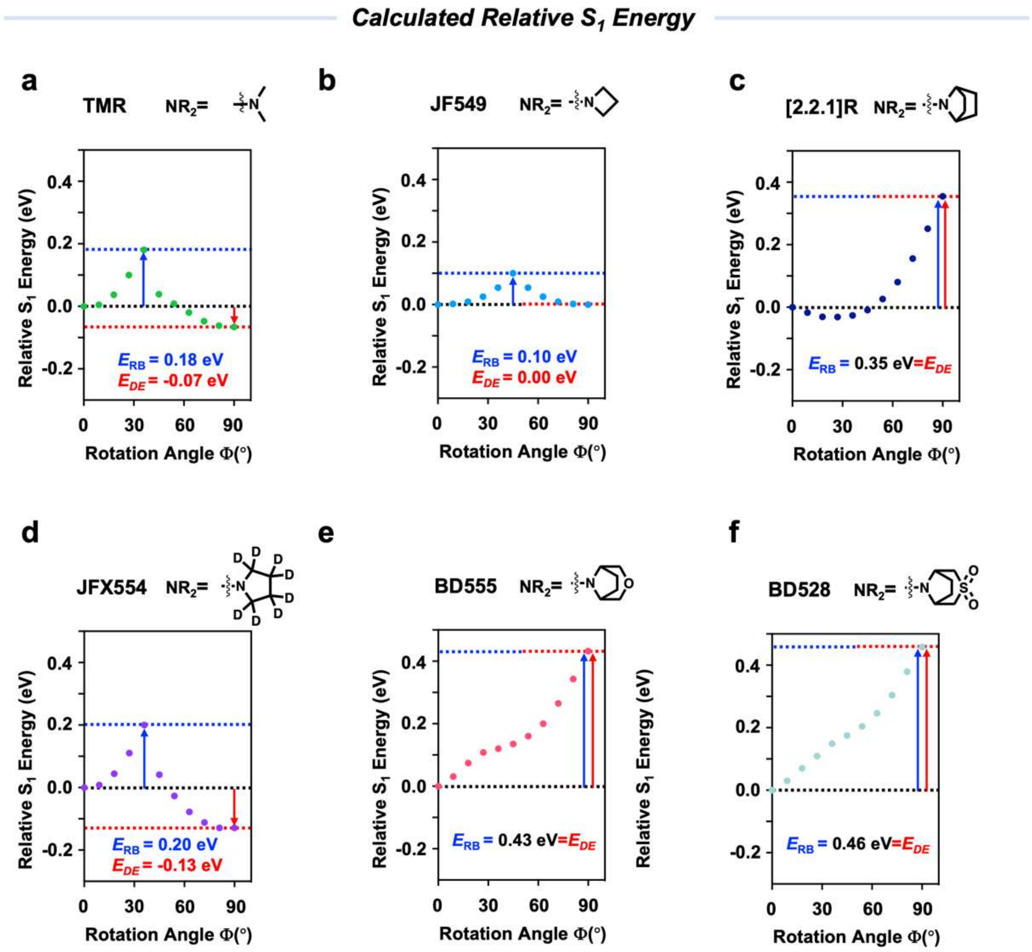


**Fig.S1 Calculated relative $S_1$ energy for O-rhodamine with various auxochromes in water.** (a) **TMR**; (b) **JF549**; (c) **[2.2.1]R**; (d) **JFX554**; (e) **BD555**; (f) **BD528**.

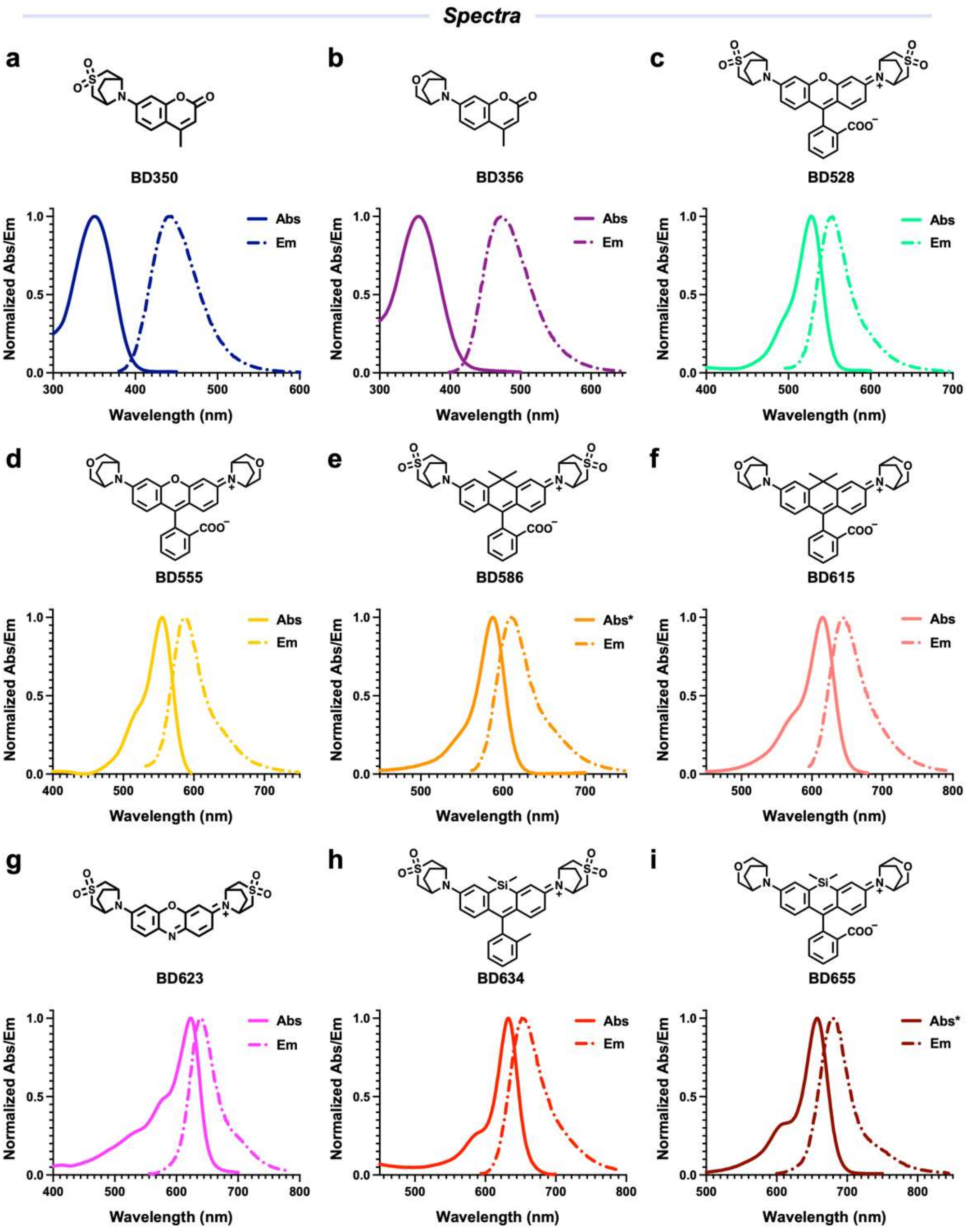


**Fig.S2 Spectra for BD dyes.**

Normalized absorption (Abs) and fluorescence emission (Em) spectra for (a) **BD350**; (b) **BD356**; (c) **BD528**; (d) **BD555**; (e) **BD586**; (f) **BD615**; (g) **BD623**; (h) **BD634**; (i) **BD655** in HEPES buffer, (10 mM, pH=7.3) (*n*=2).

*Absorption of **BD586** in (e) and **BD655** in (i) were measured in HEPES buffer (10 mM, pH=7.3) with 0.1% SDS to enforce the open-zwitterionic form.

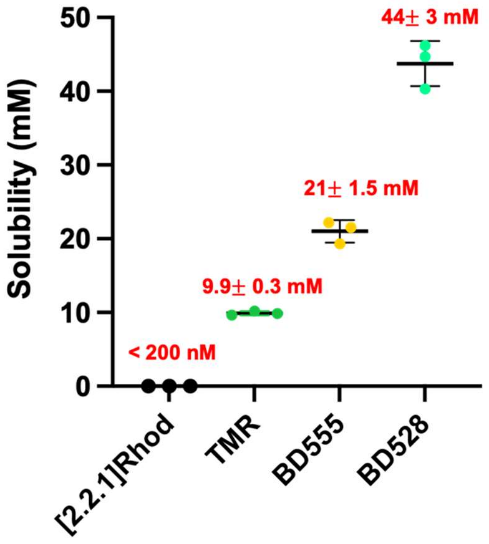


**Fig.S3 Water solubility of [2.2.1]Rhod, TMR, BD555 and BD528.**
Each dot represents an independent experiment; n = 3; data are presented as mean ± S.D.

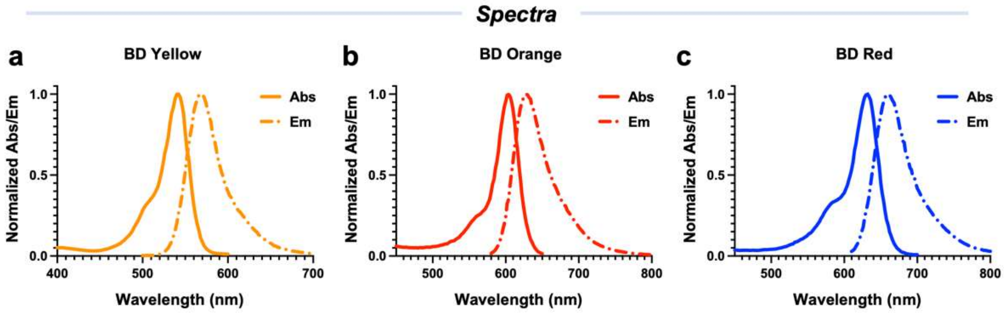


**Fig.S4 Spectra for BD derivatives for antibody bioconjugation.**
Normalized absorption (Abs) and fluorescence emission (Em) spectra for (a) **BD Yellow**; (b) **BD Orange**; (c) **BD Red** in HEPES buffer (10 mM, pH=7.3) (*n*=2).

*Photostability Tests of free dyes*

a

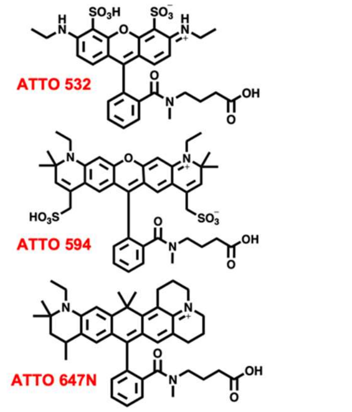


b

| Dye | $\lambda_{Abs}$(nm) | $k_{bleach}$(min$^{-1}$) | $k_{blue}$(nm/min) |
|---|---|---|---|
| BD Yellow | 540 | $-2.9 \times 10^{-3}$ | None-blue |
| ATTO 532 | 532 | $-7.8 \times 10^{-3}$ | $-6.7 \times 10^{-2}$ |
| BD Orange | 604 | $-1.8 \times 10^{-3}$ | None-boue |
| ATTO 594 | 604 | $-1.8 \times 10^{-3}$ | $-2.2 \times 10^{-2}$ |
| BD Red | 630 | $-1.3 \times 10^{-3}$ | None-blue |
| ATTO 647N | 646 | $-10 \times 10^{-3}$ | $-16 \times 10^{-2}$ |

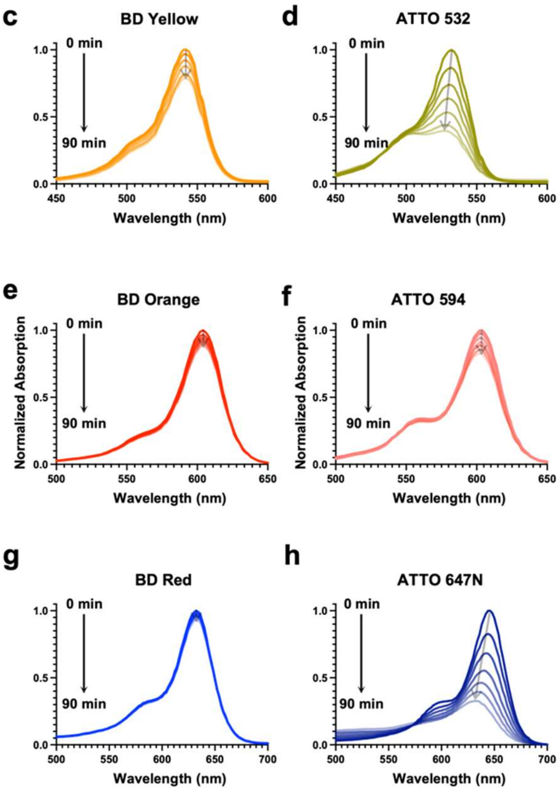


**Fig.S5 Photostability and photoblue properties of free BD derivatives for antibody bioconjugation and various ATTO dyes.**

(a) Chemical structures of various **ATTO** dyes;

(b) Photobleaching and photobluing rates of selected fluorophores in HEPES buffer (10 mM, pH 7.3) (*n*=3);

(c) to (d) The absorption of dyes over time demonstrates the photobleaching of photooxidation-resistant, nonbluing fluorophore **BD Yellow** (c) in comparison with photooxidation-prone fluorophore **ATTO 532** (d) under irradiation with an LED lamp (0.5 W/cm$^2$, 520-530 nm).

(e) to (f) The absorption of dyes over time demonstrates the photobleaching of photooxidation-resistant, nonbluing fluorophore **BD Orange** (e) in comparison with photooxidation-prone fluorophore **ATTO 594** (d) under irradiation with an LED lamp (0.5 W/cm$^2$, 590-600 nm).

(g) to (h) The absorption of dyes over time demonstrates the photobleaching of photooxidation-resistant, nonbluing fluorophore **BD Red** (g) in comparison with photooxidation-prone fluorophore **ATTO 647N** (h) under irradiation with an LED lamp (0.5 W/cm$^2$, 620-630 nm).

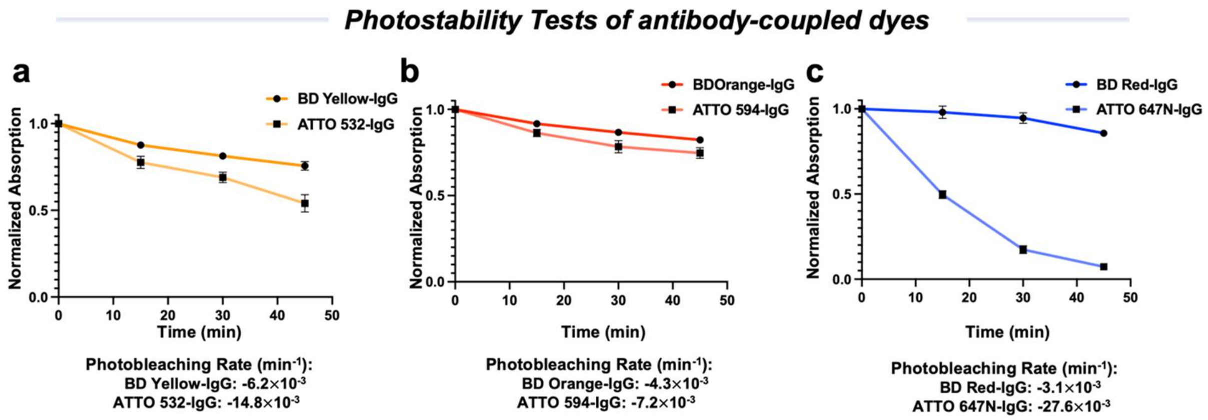


**Fig.S6 Photostability of antibody-coupled BD derivatives and their ATTO counterparts.**
(a) to (c) Photobleaching curves of antibody-coupled dyes in HEPES buffer (10 mM, pH=7.3), (n=3, error bar show mean± S.D.).
(a) Absorption at $\lambda_{max}$ of **BD Yellow** (DOL=3.27) and **ATTO 532** (DOL=3.47) were plotted as a function of irradiation time with an LED lamp (0.5 W/cm$^2$) at 520-530 nm;
(b) Absorption at $\lambda_{max}$ of **BD Orange** (DOL=2.63) and **ATTO 594** (DOL=2.50) were plotted as a function of irradiation time with an LED lamp (0.5 W/cm$^2$) at 590-600 nm;
(c) Absorption at λmax of **BD Red** (DOL=2.42) and **ATTO 647N** (DOL=2.02) were plotted as a function of irradiation time with an LED lamp (0.5 W/cm$^2$) at 620-630 nm.

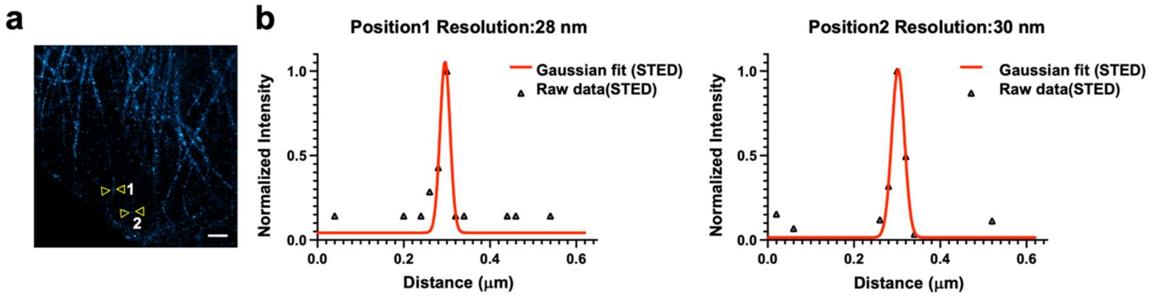


**Fig.S7 Immunofluorescence STED imaging under high-intensity depletion laser using BD Red.**
(a) STED images of a-tubulin structures in fixed-HeLa cells labeled by indirect immunofluorescence with a secondary antibody labeled with **BD Red** (DOL=2.42), 640 nm Ex./650-700 nm Em., STED at 775 nm (~200 mW), scale bar=1 μm;
(b) Line-scan profile of fluorescence intensity at the yellow arrow in (a) under high resolution STED conditions.

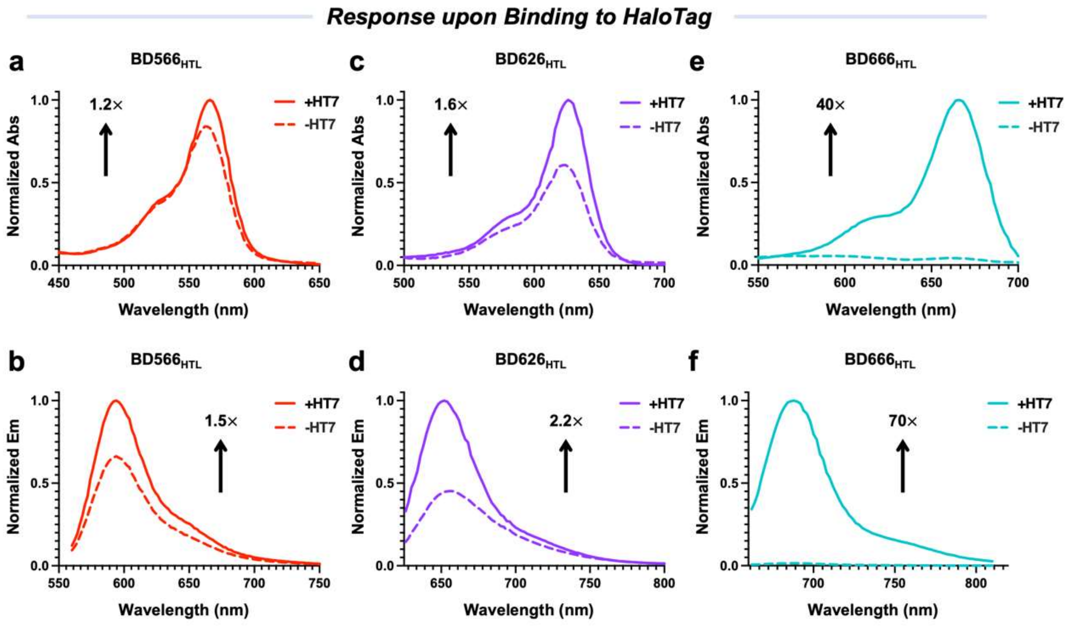


**Fig.S8 Response of BDHTLs to HaloTag.**

Absorption (a, c, e) and fluorescence emission (b, d, f) spectra of 2 μM of **BD566HTL** (a-b), **BD626HTL** (c-d), **BD666HTL** (e-f) measured in the absence (dashed line) and presence of HaloTag (5 μM, solid line) after 2 h incubation in HEPES buffer (10 mM, pH 7.3) (*n*=3). The numbers indicate the ratio of absorption maximum or emission maximum of each **BDHTL**s in the presence and absence of HaloTag.

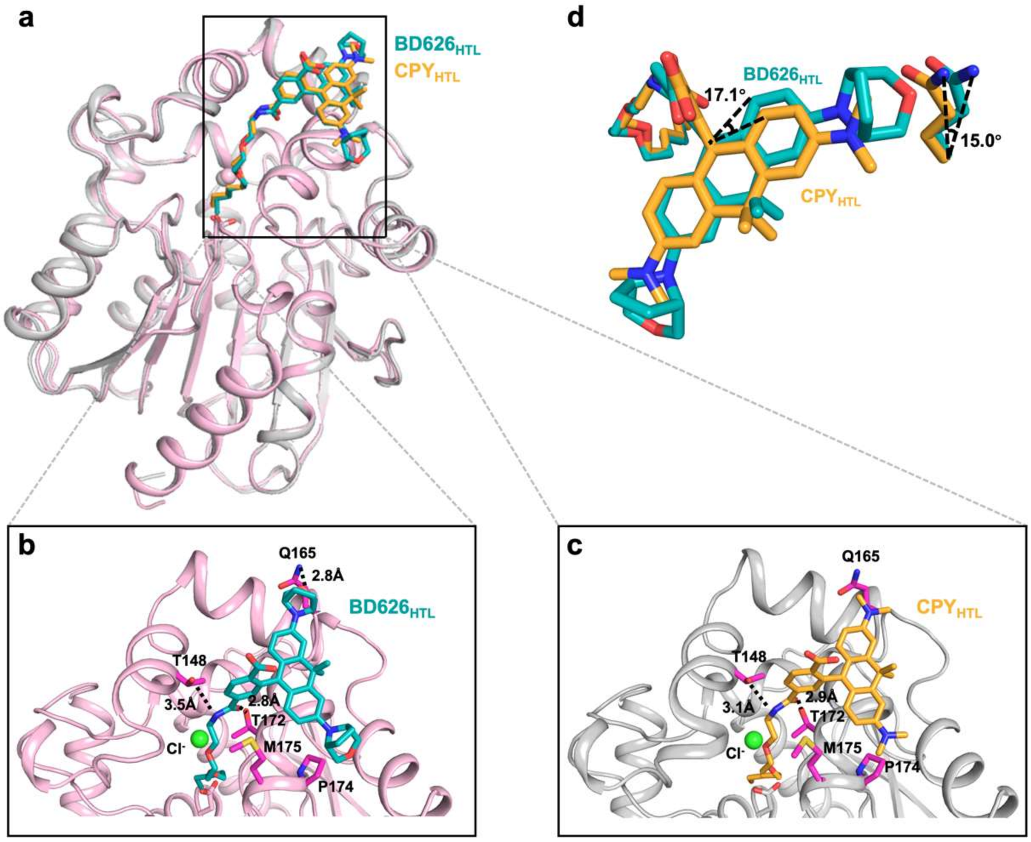


**Fig.S9 A crystal structure of BD626$_{HTL}$ at the HaloTag7 shows additional interactions between dye and protein**.

(a) Structural comparison between **HT7-CPY** (cyan, PDB 6Y7B) and **HT7-BD626** (yellow, PDB 9JHA);

(b-c) Close-ups of the substrate binding sites. Proteins are represented as pink or gray cartoons. Fluorophores and residues are shown as sticks. A polar interaction is identified between the bridged auxochrome and Q165 and shown as dashed lines with annotated distances;

(d) A overlaid comparison of **CPY$_{HTL}$** and **BD626$_{HTL}$** in the pocket of HT7.

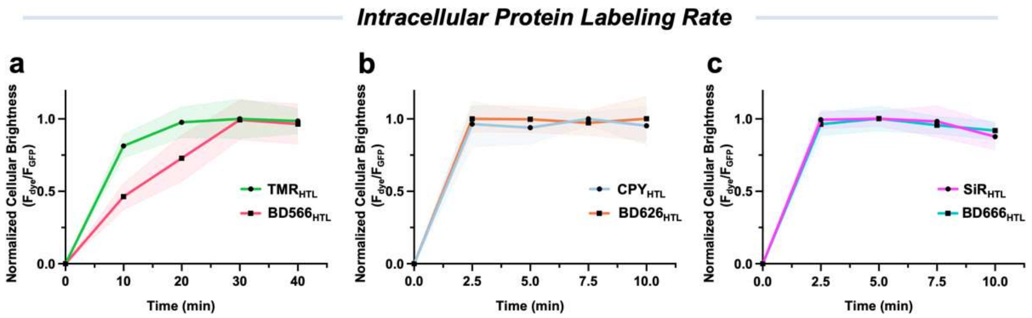


**Fig.S10 Intracellular protein labeling rate of BDHTLs and control dyes.**

Normalized ratio of **BDHTL**s and control ligands (200 nM each) fluorescence to GFP fluorescence at different time point in HeLa cells stably expressing H2B-HaloTag7-GFP; error bar show mean± S.D.; n=100 cells in each group were examined in 2 independent experiments.

**(a) BD566HTL** v. s. **TMRHTL**; (b) **BD626HTL** v. s. **CPYHTL**; (c) **BD666HTL** v. s. **SiRHTL**.

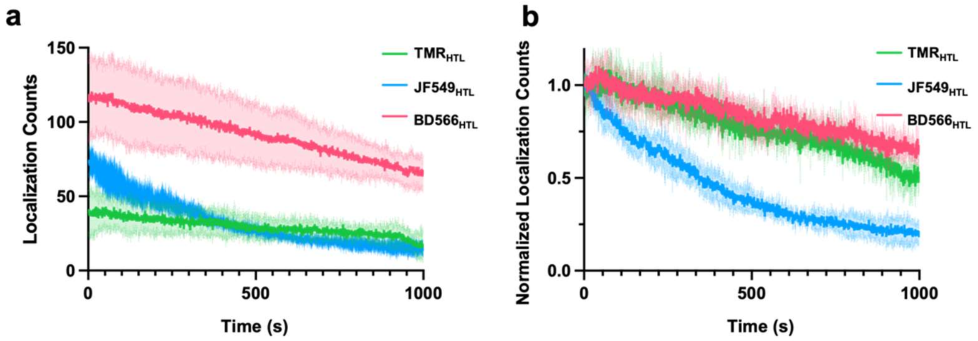


**Fig. S11 Single-molecule localization number per nuclei over time.** (a) Original data; (b) Normalized data, with the initial number as 1. Error bar show mean ± S.D.

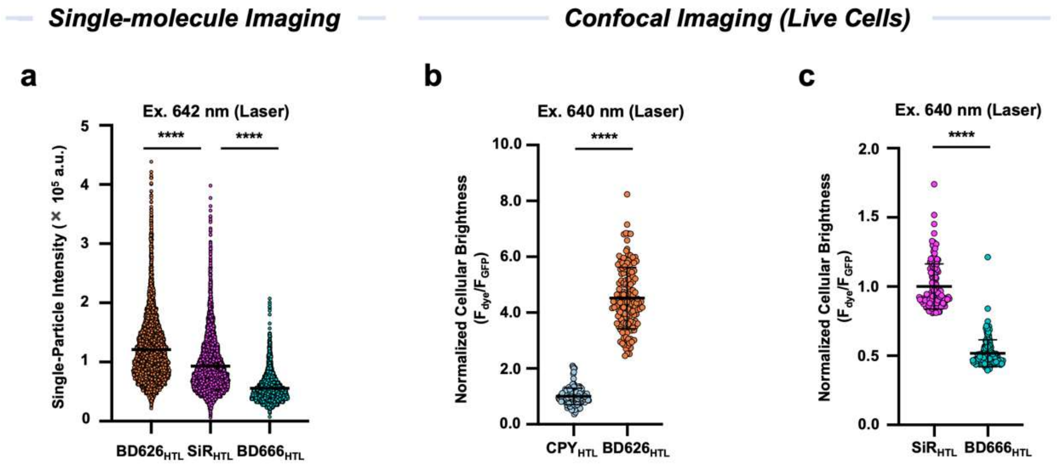


**Fig.S12 Brightness of BDHTLs and control ligands on mammalian cells under 633/640-nm lasers**.

(a) Comparison of brightness on single-molecule level in fixed U-2 OS cells stably expressing H2B-HaloTag7 labeled with **BD626HTL**, **SiRHTL** and **BD666HTL** (2.5 pM, 30 min, 37°C, one wash) under 642-nm laser;

For **BD626HTL**, n= 5158; **SiRHTL**,n= 5591; **BD666HTL** n= 4476; p****< 0.0001, t-test; error bar show mean± S.E.M.

(b-c) Comparison of apparent cellular brightness in live HeLa cells stably expressing H2B-HaloTag7-GFP labeled with different HaloTag ligands (200 nM, 1 h, 37°C, one wash) under 640-nm laser. The fluorescence intensity from single nuclei was normalized by the cytosolic GFP fluorescence intensity of the same cell (expression control) to assess the cellular brightness (error bar show mean ± S.D.);

For **CPYHTL**, $n$=154 cells, **BD626HTL**, $n$=149 cells; $p^{****}$< 0.0001, t-test;

For **SiRHTL**, $n$=150 cells, for **BD666HTL**, $n$=158 cells; $p^{****}$< 0.0001, t-test.

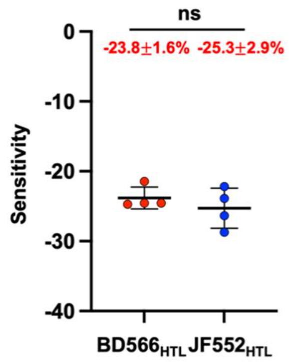


**Fig.S13 Comparison of the voltage sensitivities between BD566HTL and JF552HTL on Voltron2 in HEK293T cells.** The membrane potential was controlled via whole-cell voltage clamp, and step waveforms were applied from -70 mV to + 30 mV. n=4, error bar show mean ± S.D.

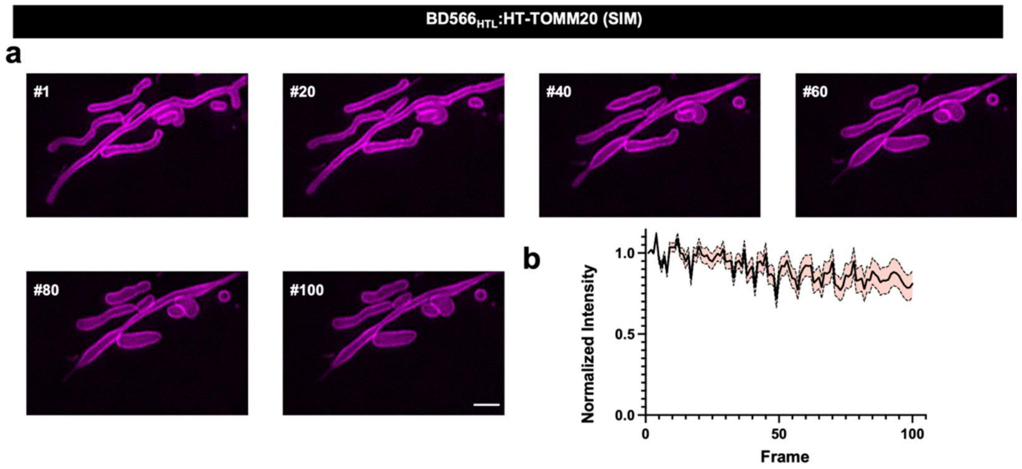


**Fig.S14 BD566HTL enables time-lapse SIM imaging.**
(a) Time-lapse SIM images of live COS-7 cells stably expressing TOMM20-HT7 labeled with **BD566HTL** (500 nM, 1 h, 37°C, one wash), scale bar=2 μm;
(b) Normalized fluorescence decay curves of samples in (a), *n*=4 in 2 independent experiments; error bar show± S.D.

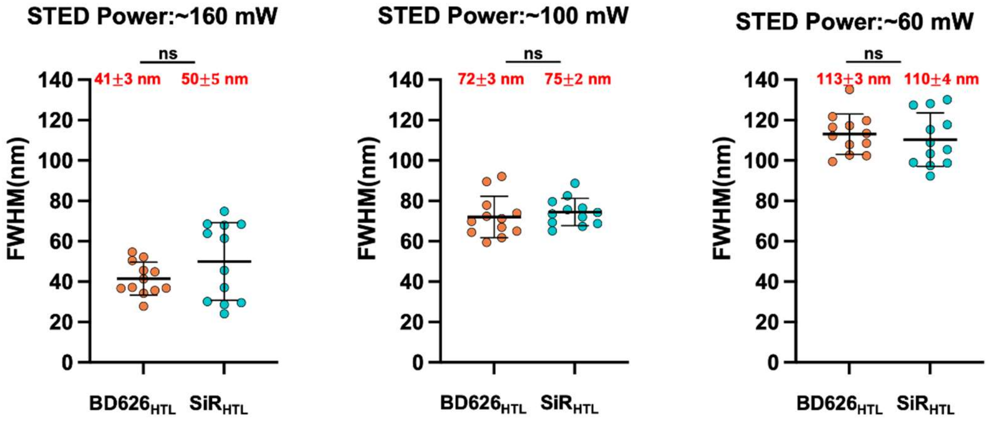


**Fig.S15 Comparison of the FWHM achieved under different STED microscopy conditions of live HeLa cells expressing HT7-Cep41 and stain with BD626HTL or SiRHTL (500 nM, 30 min, 37°C, one wash).** (n>12 filements from 3 samples).

### Methods

#### Optical spectroscopy

UV-vis absorption and fluorescence

Fluorescent dyes were prepared as stock solutions in DMSO at concentration of 1 mM and diluted such that the DMSO concentration did not exceed 0.5% (vol/vol). All measurements were taken at ambient temperature (23 $\pm$ 2 °C) in 10 mM air-saturated HEPES buffer, pH=7.3 unless otherwise noted. UV-vis absorption spectra of dye solutions were measured using a Deutta spectrometer (HORIBA Instruments, Japan) in a 3.5 mL square quartz cuvette (1 cm path length, Starna Cells). Emission spectra were measured using a Shimadzu RF-5301PC spectrofluorometer in the same cuvette. Normalized spectra are shown for clarity.

Extinction coefficient ($\varepsilon$ and $\varepsilon_{max}$)

Fluorescent dyes were quantified by qNMR using a dioxane standard (2.0 μL). A dilution series of 0.5, 1.0, 1.5, 2.0 and 2.5 μM of fluorophore were prepared in HEPES buffer (10 mM, pH=7.3) for $\varepsilon$ (extinction coefficient). The $\varepsilon_{max}$ (maximal extinction coefficient) was measured in 0.1% (vol/vol) trifluoroacetic acid (TFA) in 2,2,2-trifluoroethanol (TFE) for compounds **BD528, BD555, TMR, BD586, BD615** and **CPY** or 0.1% TFA in ethanol for compounds **BD350, BD356, coumarin 460, BD623, oxazine 1, BD634, SiTMR, BD655** and **SiR**. Absorbance spectra were recorded on a Deutta spectrometer (HORIBA Instruments, Japan) and the absorbance maxima were plotted against the concentration. A linear function was fitted to the data using Origin 2023 and extinction coefficients were calculated from slope following the Lambert-Beer's Law. Reported values for $\varepsilon$ and $\varepsilon_{max}$ are averages (*n*=3).

Quantum yield ($\Phi$)

Absolute fluorescence quantum yield ($\Phi$) were measured in HEPES buffer (10 mM, pH=7.3) using dilute samples ($A$ < 0.1) on a FLS 980 lifetime and steady state spectrometer (Edinburgh Instruments, UK) which contained an integrating sphere to determine photons absorbed and emitted. Self-absorption corrections were performed using the instrument software. Reported values for $\Phi$ are averages (*n*=3).

Fluorescence Lifetime ($\tau$)

Fluorescence lifetimes ($\tau$) were measured on a FLS 980 lifetime and steady state spectrometer (Edinburgh Instruments, UK). Measurements were carried out in HEPES buffer (10 mM, pH=7.3) for **BD350, BD356, coumarin 460, BD528, BD555, TMR, BD615, CPY, BD623, oxazine 1, BD634** and **SiTMR** and 0.1% TFA in ethanol for **BD586**, **BD655** and **SiR**. Single exponential fitting were performed using the instrument software to get $\tau$ values. Reported values for $\tau$ are averages (*n*=3).

The lactone-zwitterion equilibrium constant ($K_{L-Z}$)

The lactone-zwitterion equilibrium constant $K_{L-Z}$ were calculated by equation (1) as reported[1]:

$$K_{L-Z} = \frac{\varepsilon_{dw}/\varepsilon_{max}}{1-\varepsilon_{dw}/\varepsilon_{max}} \qquad (1)$$

$\varepsilon_{dw}$ is the extinction coefficient of a fluorophore in a 1:1 spectral-grade dioxane (Sigma-Aldrich): milliQ $H_2O$ (vol/vol) solvent mixture.
$\varepsilon_{max}$ (maximal extinction coefficient) were measured in 0.1% (vol/vol) trifluoroacetic acid (TFA) in 2,2,2-trifluoroethanol (TFE) for **BD528, BD555, TMR, BD586, BD615** and **CPY** or 0.1% TFA in ethanol for **BD655** and **SiR**.

**Water solubility measurements**
**[2.2.1]Rhod**, **TMR**, **BD528** and **BD555** was dissolved in 1 mL MeOH with final stock concentration ranging from 10 to 50 mM. The solvent was removed in vacuo. Afterthat, 250 μL dd$H_2O$ was added. The mixture was vortexed and sonicated in an ultrasonic bath for 45 min at 95 °C. The undissolved pellet was separated via centrifugation at 15,000 rpm for 5 min. The concentration of dye in the supernatant was measured by using a Deutta spectrometer (HORIBA Instruments, Japan) and then calculated by Lambert-Beer's Law.

**Bulk Photobleaching**
Fluorescent dyes were diluted with air-saturated HEPES buffer (10 mM, pH=7.3) to 10 μM and followed by irradiation with an LED lamp (520-530 nm, 0.5 W/cm$^2$ for **BD528**, **BD555**, **TMR**, **JF549**, **JF525**, **Rhod524**, **BD Yellow** and **ATTO 532**; 590-600 nm 0.5 W/cm$^2$ for **BD615, CPY, BD Orange and ATTO 594**; 620-630 nm, 0.5 W/cm$^2$ for **BD Red and ATTO 647N**). UV-Vis spectra were recorded on a microplate reader (TECAN Infinite M Nano$^+$, Switzerland) in 96-well plates with optical bottom (Costar) every 15 min over the course of 90 min. This experiment was independently repeated three times.

**Nonspecific labeling of bovine serum albumin (BSA) with dye photo-byproducts**
**JF549/BD555** (10 μM) and BSA (5 mg/mL) were dissolved in 1 mL air-saturated PBS buffer (pH=7.4). The mixture was irradiated under an LED lamp (0.5 W/cm$^2$, 520-530 nm) for 30 min with continuous shaking. Afterwards, the samples were mixed with 5× loading buffer (LBP6510-10, COFITT) supplemented with 2-mercaptoethanol to 10% final concentration. Samples were heated for 5 min at 95 °C, and 10 μL aliquot of each sample was loaded into the gradient 4-15% Mini-PROTEAN TGX™ precast protein gels (Bio-Rad). SDS-PAGE was performed in Mini-PROTEAN Tetra Cell (Bio-Rad) apparatus with voltage set to 125 V until the loading dye had run out of the gel. In-gel fluorescence was acquired with Bio-Rad Image System (Cy3 channel). Afterwards, the gels were stained with Coomassie Brilliant Blue G-250 for protein quantification. Image quantification was performed using Fiji software region measurement tool.

**Computational chemistry**
All calculations were carried out with Gaussian 16 software[2]. The B3LYP functional[3] was adopted for all calculations in combination with the D3BJ dispersion correction[4]. For geometry optimization, frequency calculations and TD-DFT calculations, the 6-311G(d) basis set[5] was used. The vertical ionization energies were further derived from single point calculations using the 6-311+G(d,p) basis set[6]. The SMD solvation model[7] was used to take account of the solvation effect of water for all the calculations.

**Cell Biology**
<u>Cell culture</u>

HeLa, COS-7, and U-2 OS EF1α-H2B-HaloTag7-expressing cells were cultured in Dulbecco's modified Eagle medium (DMEM, 4.5g/L glucose, Macgene #CM10017) supplemented with 10% (vol/vol) fetal bovine serum (FBS, ThermoFisher 10500064) and 1% (vol/vol) Penicillin-Streptomycin Solution (10 IU/ml penicillin / 0.1 mg/ml streptomycin, Macgene # CC004) in a humidified 5% $CO_2$ incubator at 37 °C. To prepare cells stably expressing histone H2B-HaloTag7-GFP, HeLa cells were integrated with an expressing plasmid via the piggyback transposase. Cells were split every 2-3 days or at confluency, and were regularly tested for mycoplasma contamination.

Transfection of cells

HeLa and COS-7 cells were transfected 24-48 h prior to imaging using lipofectamine™ 3000 (L3000015; Thermo Fisher Scientific). Transfection was carried out according to the manufacturer's protocols. After 6-12 h, the residual transfection reagent was removed and washed with DMEM. The cells were maintained in a fresh medium for another 18-24 h before staining with dyes.

Fixation for immunofluorescence labeling

HeLa cells were grown for 48-72 h on glass coverslips and washed three-times with PBS buffer (pH=7.4) before fixation. Cells were pre-fixed in 4% (vol/vol) formaldehyde (FA) in PBS for 45 s, permeabilized in 0.4% (vol/vol) Triton X-100 in PBS for 3 min and fixed in 4% (vol/vol) PFA and 0.1% (vol/vol) glutaric dialdehyde (GA) in PBS for another 15 min at 37°C. PFA and GA was quenched by 100 mM $NH_4Cl$ and 100 mM glycine in PBS for 10 min. After washing twice for 10 min in PBS, the cells were labeled with corresponding antibody.

**Antibody labeling using dye-NHS ester**

**BD Yellow/Orange/Red** NHS-esters (500 nmol) were dissolved in 20 μL DMSO. Then the stock solutions were mixed with 1 mg secondary antibody (in PBS buffer, pH=8.3, adjusted by $NaHCO_3$ solution) in a proportion of 20-25 equivalents (dye/protein). After that, the mixture was incubated at room temperature for 2.5 h in the dark and the mixture was then purified by the gel filtration column (Sephadex G-25, Sigma-Aldrich). UV-Vis measurements were performed in a NanoDrop One small volume spectrometer (Thermo Scientific) to determine the degree of labeling (DOL, dye/protein).

**UV-Vis and fluorescence spectra of dye-HaloTag conjugates**

HaloTag7 protein was recombinantly expressed and purified following an established protocol (REF) and stored asa 100 μM solution in 50 mM HEPES buffer containing 50 mM NaCl, pH 7.4. Compounds **22-28** were diluted to 2 μM in 10 mM HEPES, pH 7.3. Then an aliquot of HaloTag7 protein (3.0 equiv.) was added and the resulting mixture was incubated at 37°C for 4 h. UV-Vis and emission spectra measurements were recorded on a microplate reader (TECAN Infinite M Nano$^+$, Switzerland) in 96-well plates with optical bottom (Costar). Reported values are averages (n = 3 independent experiments).

**Cloning, Protein Expression, and Purification**

HaloTag7 was cloned in a pET21b(+) vector (Zoman) for production in Escherichia coli, featuring an N-terminal His7 tag and a PreScission Protease (PP) cleavage site. Cloning was performed by Gibson assembly using E.coli DH5α cells (Beijing Zoman Biotechnology # ZC101). Proteins were expressed in E. coli strain BL21(DE3) (Beijing Zoman Biotechnology # ZC121). Lysogeny broth (LB) cultures were grown at 37 °C to an optical density at 600 nm (OD600) of 0.8. Protein expression was induced by the addition of 1 mM isopropyl β-D-thiogalactopyranoside (IPTG), and cells were grown at 37 °C for 4 hours and 18°C overnight. The cells were centrifuged at 4,000 g, 4 °C for 20 min. The pellet was collected and sonicated on ice in standard TBS buffer (20 mM Tris-HCl, 150 mM NaCl, pH 8.0) with 2 mM β-mercaptoethanol, and 1 mM phenylmethanesulfonylfluoride (PMSF). Then, the cell debris was removed by centrifugation at 40,000g (JA-20, Beckman) at 4 °C for 30 min. The supernatant was loaded onto the TALON column (Clontech) and washed with 20 mM Tris-HCl (pH 8.0) and 500 mM NaCl, 2 mM β-mercaptoethanol, followed by TBS buffer with 10 mM imidazole. The column was eluted with 250 mM imidazole in TBS. The eluted protein was concentrated using Amicon Ultra 15-mL 10-kDa centrifugal filters (MilliporeSigma), and loaded onto a Superdex200 10/300 GL column (Cytiva) in TBS buffer, followed by purification with ÄktaPure chromatography system (Cytiva). N-terminal His7 tag was removed by overnight cleavage with self-purified GST tagged PreScission Protease at 4 °C as previously described. The proteins were further purified by affinity-tag purification using a 5 mL HisTrap FF crude column (Cytiva) and a GSTrap HP Column (Cytiva) on an ÄktaPure chromatography system (Cytiva), where the flow-through fractions were collected. Proteins were further separated by size exclusion chromatography (Superdex200 10/300 GL colum, Cytiva) in a mobile phase of standard TBS buffer and concentrated using Amicon Ultra 15-mL 10-kDa centrifugal filters (MilliporeSigma). The protein purity were verified via SDS−PAGE. Proteins were aliquoted and stored at −80 °C after being flash-frozen in liquid nitrogen.

**Protein Labeling and Crystallization**

HaloTag protein labeling was performed 2 hours at 37°C at the final concentration of 10 μM (**BD626**$_{\mathbf{HTL}}$) in the presence of 5 μM HaloTag7 protein. The protein was concentrated with Amicon Ultra 15-mL 10-kDa centrifugal filters (MilliporeSigma) and the excess of fluorophore substrate was removed by a 5 mL HiTrap Desalting column (Cytiva). Dye-protein conjugate was further purified by size exclusion chromatography (Superdex200 10/300 GL column, Cytiva) in a mobile phase of crystallization buffer (10 mM Tris-HCl, 100 mM NaCl, pH 8.0) and concentrated using Amicon Ultra 15-mL 10-kDa centrifugal filters (MilliporeSigma) to a final concentration of 30 mg/mL. Dye-protein conjugate was submitted to crystallization trials using different commercial screens via mixing in a 200 nL final volume protein solution/crystallization solution (1:1) using a NT8 Drop Setter for Protein Crystallization (Formulatrix).

**BD626**$_{\mathbf{HTL}}$**-HaloTag7 Crystallization**

Crystallization was performed at 20 °C using the vapor diffusion method. Crystals of HaloTag7 labeled with **BD626**$_{\mathbf{HTL}}$ fluorophore substrate was grown by mixing equal volumes of a 30 mg/mL protein solution in crystallization buffer (10 mM Tris-HCl, 100 mM NaCl, pH 8.0) and a reservoir solution containing 0.1 M sodium cacodylate (pH 7.3), 0.2 M ammonium sulfate, and 32% (m/v) PEG 8000. The crystals were briefly washed in a cryoprotectant solution consisting of the reservoir solution with glycol added to a final concentration of 20% (v/v), prior to flash-cooling in liquid nitrogen. Single-crystal X-ray diffraction data were collected at 100 K in the X-ray crystallography facility at Tsinghua University (XtaLAB Synergy Custom FRX and a hybrid photon counting detector HyPix6000, Rigaku, Japan). The structure of HaloTag7 labeled with **CPY**$_{\mathbf{HTL}}$ was determined by molecular replacement (MR) using Phaser[8] and the coordinates of Protein Data Bank (PDB) entry 6Y7A as a search model. Geometrical restraints for **BD626**$_{\mathbf{HTL}}$ were generated using JLigand[9] program in CCP4[10]. The final models were optimized in iterative cycles of manual rebuilding using Coot[11,12] and refinement using phenix.refine[13]. Data collection and refinement statistics are summarized in Table S2, and the model quality was validated with MolProbity[14] as implemented in PHENIX[15].

### Confocal Imaging

Confocal imaging was performed on a Leica SP8 FALCON microscope (Leica Microsystems) equipped with a Leica TCS SP8 X scanhead; a SuperK white light laser (WLL), Leica HyD SMD detectors and a 100×/1.40 oil objective. For Fig. 4g and 4i, **CPY$_{HTL}$ and BD626$_{HTL}$** were excited at 620 nm (WLL, 10% intensity, Em:625-657 nm). For Fig. 4g and 4k, **SiR$_{HTL}$ and BD666$_{HTL}$** were excited at 658 nm (WLL, 10% intensity, Em:663-713 nm). Other selected images are acquired with an Abberior STEDYCON microscope (Abberior Instruments GmbH) equipped with 488(4.4 µW, Em:500-550 nm)/561(5.0 µW, Em:575-625 nm)/640 (8.5 µW, Em:650-750 nm) laser and a 100×/1.45 oil objective. The Leica microscope was equipped with a live cell incubator (Life Imaging Services, 37 °C).

Live-cell fluorescence signal, labeling kinetics, and signal-over-background ratio

HeLa cell lines stably expressing histone H2B-SNAP-eDHFR:L28C-HaloTag7-GFP were seeded. For **TMR$_{HTL}$, JF549$_{HTL}$ and BD566$_{HTL}$**, cells were labeled with 200 nM dyes for 2 h at 37°C and washed once with culture medium (DMEM) before imaging. For **CPY$_{HTL}$ and BD626$_{HTL}$**, cells were labeled with 200 nM dyes for 1 h at 37°C and washed once before imaging. For **SiR$_{HTL}$ and BD666$_{HTL}$** cells were labeled with 200 nM dyes for 1 h at 37°C and directly used for imaging without washing step. Cells were focused in the 488 channel and signals from the 561 or 640 channels were collected. The fluorescence signal intensity was measured from 12 fields of view (FOVs) in 3 repetitive experiments ($n \geq 150$ cells) and normalized to the GFP signal to correct for expression levels.

Live-cell labeling kinetic experiments were performed using 200 nM dye in the staining media. The cells are imaged every 10 min for **TMR$_{HTL}$** and **BD566$_{HTL}$** and 2.5 min for **CPY$_{HTL}$, BD626$_{HTL}$, SiR$_{HTL}$ and BD666$_{HTL}$**. The nuclear fluorescence intensity was quantified from >100 cells in total overtime from 3 individual experiments, averaged and normalized to the initial and maximal fluorescence signal.

To determine signal-over-background ratio, live HeLa cells stably expressing H2B-SNAP-eDHFR:L28C-HaloTag7-GFP and wild-type HeLa cells were co-seeded (1:1) at 37 °C in a humidified 5% (v/v) $CO_2$ environment for 1-2 days. The cells were labeled with 200 nM of **SiR$_{HTL}$** and **BD666$_{HTL}$** and directly imaged after 60 min. The white dashed lines (Fig 4g) indicate the wild-type HeLa cells, which were located based on bright-field images. The fluorescence signal in nucleus (HeLa H2B-SNAP-eDHFR:L28C-HaloTag7-GFP cells) and cytoplasma (wild-type HeLa cells) was corrected by subtracting the average density of adjacent background regions. The $F_{nuc}/F_{cyt}$ values were determined based on 150 cells from 3 repetitive experiments.

### STED Imaging

STED imaging was performed on an Abberior Facility microscope (Abberior Instruments GmbH) equipped with 640 nm laser and a 100×/1.4 oil objective. Pixel sizes of 20 nm used for single-frame super-resulution imaging. **BD Orange**, **BD Red** and **BD626$_{HTL}$** and control dyes were excited at 640 nm (8.5 µW, Em:650-750 nm) and STED was performed using a pulsed depletion laser at 775 nm (~60 to 200 mW) with gating of 1-7 ns and dwell time of 10 µs.

For 3D STED imaging, **BD626HTL** were excited at 640 nm (10 μW, Em:650-750 nm) and STED was performed using a pulsed depletion laser at 775 nm (~80 mW, 80% to 2D and 20% to 3D) with pixel sizes of 40 nm and dwell time of 8 μs.
For time-lapse STED imaging, **BD626HTL** and control dyes were excited at 640 nm (5 μW, Em:650-750 nm) and STED was performed using a pulsed depletion laser at 775 nm (~60 mW) with gating of 1-7 ns and dwell time of 10 μs, 20s/frame.

**SIM Imaging**
SIM imaging was performed on the Polar-SIM system from Airy Technology. The SIM is equipped with a wide-field objective lens (CFI SR HP Apo TIRF 100× 1.49 NA, Nikon). **BD566HTL** were excited at 561 nm (200 mW, 25%, Em:575-625 nm ) The original images were captured using an sCMOS camera (Hamamatsu, Orca Flash 4.0 version 3). These images were then reconstructed using Airy SIM's specialized processing software.

**Single-molecule Imaging**
Microscopy setup
All single-molecule tracking (SMT) experiments were performed on a custom-built Nikon Ti2 microscope as before[16]. It is equipped with two EM-CCD cameras (Andor, iXon Ultra 897), a 100x/NA 1.49 oil-immersion TIRF objective (Nikon apochromat CFI Apo TIRF 100x oil), a perfect focusing system (Applied Scientific Instrumentation), and a stable-top chamber for maintaining 37°C and 5% $CO_2$ (Tokai Hit). The HILO illumination was achieved by a Nikon TIRF module, multiple lasers (405 nm: maximum 140 mW, OBIS, Coherent; 560 nm: maximum 1 W, MPB; 642 nm: maximum 1.5 W, MPB) controlled by the AOTF system (AA Opto-Electronic, AOTFnC-VIS-TN), a multi-band dichroic mirror (405/488/561/633 nm quad-band, Semrock) and emission filters (**TMRHTL/JF549HTL/BD566HTL**: Semrock 593/40 nm band-pass filter; **BD626HTL/SiRHTL/BD666HTL**: Semrock 676/37 nm bandpass filter).

Cell labeling for SMT
U2OS stably expressing histone H2B-HaloTag fusion proteins were generated by co-expressing the PiggyBac EF1a-H2B-HaloTag-IRES-Neo vector with the PiggyBac transposase, followed by G418 selection (400 μg/ml) for two weeks and verified by FACS sorting.U2OS cells were plated on 8-well Cellvis chambers (Cellvis # C8-1.5H-N) with #1.5 high performance cover glass, and were cultured in the complete DMEM medium (Dulbecco's Modified Eagle Medium (Gibco, C11995500BT) supplemented with 10% fetal bovine serum (VisTech, SE100-011) and 1% penicillin/streptomycin (Hyclone, SV30010)). To label Halo fusion protein in living cells, Halo ligand conjugated fluorescent dyes at indicated final concentration were added to culture medium (2.5 pM for **TMRHTL**, **JF549HTL**, **BD566HTL**, **BD626HTL**, **SiRHTL** and **BD666HTL**) and incubate at cell incubator for 30 min. The medium was then removed and two rounds of incubation with fresh medium for 30 minutes were performed to remove free dyes and fixed cells with 500 μL 4% paraformaldehyde after 30 min. Medium used in each step was pre-warmed and caution was taken to avoid cells to detach from the glass bottom.

Acquisition and analysis of SMT data
The slowSMT experiments used a long exposure time (500 ms) and continuous illumination with low laser intensity to capture the position of slow mobile molecules. Generally, each cell was captured for 2000 frames with 500 ms exposure time at 10% of maximal excitation power. The

microscope, laser lines, and camera integration were controlled using the Nikon NIS-Elements software. The slowSMT movies were analyzed with custom-made MATLAB scripts (https://gitlab.com/tjian-darzacq-lab/SPT_LocAndTrack) implementing a multiple-target tracking (MTT) algorithm[17] to generate single-molecule trajectories. The settings used in the MATLAB scripts for slowSMT were as follows: localization error, $10^{-6.25}$; deflation loops, 0; Blinking (frames), 1; maximum competitors, 3; maximum D ($\mu m^2\ s^{-1}$), 0.7. For SPT brightness, intensity values from the first frame of the imaging experiment were analyzed. For SPT duration, the single-molecule trajectories of each cell were combined and a survival curve was generated based on the 1-cumulative distribution function (1-CDF) histogram.

**Voltage Imaging**

Cell culture and transfection

HEK293T cells were incubated in Dulbecco's modified Eagle medium (DMEM, Gibco) containing 10% vol/vol fetal bovine serum (FBS, Gibco) at 37 °C with 5% $CO_2$. Cells were seeded in a 24-well plate and grown to 70–90% confluent for transfection. 500 ng plasmid of Voltron2 and 1 µl Lipofectamine 2000 reagent were mixed in Opti-MEM medium and added to the culture cell in DMEM for 4 -8h. After that, cells were digested by trypsin-EDTA (0.25%, Gibco), reseeded on a sterile 14-mm glass coverslip pre-treated with matrigel matrix, and incubated in complete medium for 24 h before **BD566$_{HTL}$** labelling.

Primary rat hippocampal neurons were digested from isolated from rat brains at postnatal day 0 (P0) and seeded on 12-mm glass coverslips pre-coated with 20 µg/mL poly-D-lysine (Sigma) and 10 µg/mL laminin (Gibco). Neurons were incubated at 37 °C with 5% $CO_2$ in neuronal culture medium (Neurobasal medium, B-27 supplement, GlutaMAX supplement and penicillin-streptomycin). Neurons were transfected on DIV8 (8 days *in vitro*). For each well of a 24-well plate, 500 ng plasmid of Voltron2 and 1 µL Lipofectamine 3000 reagent (Invitrogen) were mixed in Neurobasal medium and then incubated neurons with the mixture for 45 min at 37 °C with 5% $CO_2$. Transfected neurons were labeled and imaged after 3-10 days.

Voltage imaging

Neurons expressing Voltron2 were labeled with **BD566$_{HTL}$** (100 nM in DMEM) at 37 °C with 5% $CO_2$ for 30 min. After that, neurons were gently rinsed three times with DMEM and were transferred to Tyrode's buffer before imaging. Voltage imaging were conducted on an inverted microscope (Nikon-TiE) equipped with a objective CFl PlanFluor 40x oi, NA 1.3 (Nikon, Tokyo, Japan), one laser line (561 nm, Coherent OBlS), and one scientificCMOS camera (Hamamatsu ORCA-Flash 4.0 v2). The microscope, lasers and cameras were controlled by a custom-built software written in LabVIEW (National Instruments, 15.0 version). In order to record action potentials, fluorescence images were captured at a frame rate of 400 Hz, under ~1.6 W/cm$^2$ 561 nm illumination. Image analysis was performed in MATLAB (version R2018b) and ImageJ/Fiji (version 1.53t).

To investigate photobleaching of **BD566$_{HTL}$** and **JF552$_{HTL}$** in cells, Voltron2 was transfected in HEK293T cells. Transfected cells were labeled with **BD566$_{HTL}$/JF552$_{HTL}$** and fixed cells with 500 µL 4% paraformaldehyde after 24 h. Transferred cells to Tyrode's buffer and then illuminated cells with 561 nm (~2 W/cm$^2$) lasers.

**Imaging of Plant Cells**

*Arabidopsis thaliana* ecotype Columbia-0 (Col-0) and Nicotiana benthamiana were employed as the wild-type (WT) control. Constructs were introduced into plants via the floral-dip method of Agrobacterium tumefaciens-mediated transformation (10.1385/1-59259-827-7:091). Seeds were surface-sterilized by using a mixture of ethanol and $H_2O_2$ (4:1,70% ethanol:30% $H_2O_2$), stratified at 4 °C for 24 h in the dark. Arabidopsis was grown for 4 days on 1/2 MS medium supplemented with 1% sucrose under a 16-h light/8-h dark (23°C–25°C) photoperiod before imaging. The leaves of 6-week-old *N.benthamiana* plants were co-infiltrated with equal volumes of different Agrobacterium tumefaciens strain combinations, and the images were taken after being maintained at a temperature of 23°C–25°C) for 2 days.
Arabidopsis seedlings and N. benthamiana leaves expressing the HaloTag vector were transferred into centrifuge tubes and infiltrated with HaloTag ligand solution (200nM **JF646$_{\text{HTL}}$**, 200nM **BD626$_{\text{HTL}}$**), which was incubated for 1 hour in the dark at 23-6 °C. Careful washing for a minimum of 4 hours or up to overnight in water was essential to reduce non-specific background (10.1093/jxb/erl065). For imaging, a 4-day-old seedling was used and placed on a 35-mm-diameter, 170-μm-thick cover glass slide. Live-cell imaging was carried out on a GE Healthcare Delta Vision OMX SR imaging system furnished with a high NA 60× 1.42 objective. The GFP fluorophore was excited at 488 nm (200 mW, 20%, Em: 500-550 nm). The **JF646$_{\text{HTL}}$** and **BD626$_{\text{HTL}}$** was excited at excited at 640 nm (200 mW, 5%, Em: 650-700 nm). Reconstruction and image analysis of the TIRF-SIM images were executed using SoftWoRx v5.9 (GE).

***In vivo* confocal imaging in larval zebrafish**

For zebrafish experiments, larvae at 4-6 days post-fertilization (dpf) were used in this study. Zebrafish adults, embryos, and larvae were maintained at 28°C in system water on a 14-hour light and 10-hour dark cycle. All procedures were approved by the Institute of Neuroscience, Chinese Academy of Sciences.
The *elavl3*:H2B-HaloTag plasmid (25 ng/μL), mixed with Tol2 transposase mRNA (25 ng/μL), was injected into fertilized embryos with a Nacre background at the one-cell stage to generate chimeric transgenic fish. Before imaging, 4 dpf larvae were soaked in the dye solution (3.3 μM in system water) for 1 hour. Afterward, they were placed back in clean system water for 2 hours to wash off the surface dye, and then imaging was performed. An FV3000 confocal microscope (Olympus) equipped with a 20x (NA 1.0) water-immersion objective was used on fish embedded in 2% agarose gel to obtain 1024 x 1024 pixel z-stack images. The **BD566$_{\text{HTL}}$** was excited at excited at 561 nm (1 W, 5%, Em: 593/40 nm); The **BD626$_{\text{HTL}}$** was excited at excited at 633 nm (1.5 W, 5%, Em: 676/37 nm).

**Table S1 Data collection and refinement statistics for the crystal structure of BD626$_{HTL}$-HaloTag7. Values in parentheses are for the highest resolution shell.**

| | |
|---|---|
| **PDB code** | |
| **Data-collection statistics** | |
| Diffraction Source | XtaLAB Synergy Custom FRX |
| Wavelength (Å) | 1.54178 |
| Temperature (K) | 100 |
| Resolution range (Å) | 31.35 - 1.698 (1.759 - 1.698) |
| Space group | $P4_32_12$ |
| Unit cell parameters | |
| *a, b, c* (Å) | 62.696, 62.696, 164.386 |
| *α, β, γ* (°) | 90, 90, 90 |
| Total reflections | 1004475 |
| Unique reflections | 37009 (3540) |
| Multiplicity | 27.1(4.9) |
| Completeness (%) | 99.55 (97.84) |
| Mean I/σ (*I*) | 39.47(1.70) |
| Wilson *B* factor (Å$^2$) | 21.37 |
| $R_{meas}$ | 0.126(0.604) |
| $R_{p.i.m.}$ | 0.020(0.256) |
| $CC_{1/2}$ | 0.998(0.859) |
| **Structure-refinement statistics** | |
| Reflections used in refinement | 36979 (3540) |
| Reflections used for $R_{free}$ | 1841 (170) |
| $R_{work}$ | 0.1912 (0.2334) |
| $R_{free}$ | 0.2404 (0.2723) |
| No. of protein chains in asymmetric unit | 1 |
| No. of non-H atoms | 2857 |
| Protein residues | 299 |
| R.m.s.d., bond lengths (Å) | 0.047 |
| R.m.s.d., bond angles (°) | 0.9 |
| Ramachandran statistics (%) | |
| Favored | 95.53 |
| Allowed | 4.47 |
| Outliers | 0 |
| Rotamer outliers (%) | 0 |
| Clashscore | 1.28 |
| Average *B* factor (Å$^2$) | |
| Overall | 24.46 |
| Macromolecules | 22.27 |
| Ligands | 35.42 |
| Solvent | 34.26 |

**Synthesis and Characterization of New Compounds**

**General Information**

Unless otherwise mentioned, all reactions were carried out under a nitrogen atmosphere with dry solvents under anhydrous conditions. All the chemicals were purchased at the highest commercial quality and used without further purification unless otherwise stated. Reactions were conducted in round-bottomed flasks or septum-capped crimp-top vials containing Teflon-coated magnetic stir bars. The heating of reactions were accomplished with a silicon oil bath on a stirring hotplate equipped with an electronic contact thermometer to maintain the indicated temperatures. Anhydrous N, N-dimethylformamide (DMF) and tetrahydrofuran (THF) was purchased from Innochem (China). 3-thia-8-azabicyclo [3.2.1] octane 3,3-dioxide hydrochloride, 3-thia-8-azabicyclo [3.2.1] octane 3,3-dioxide hydrochloride and 3-thia-8-azabicyclo [3.2.1] octane hydrochloride were perchased from PharmaBlock (China).

Reactions were monitored by Thin Layer Chromatography on plates (GF254) supplied by Yantai Chemicals (China) using UV light as a visualizing agent and an ethanolic solution of phosphomolybdic acid and cerium sulfate, and heat as developing agents or by LC/MS (4.6 mm × 150 mm 5 μm C18 column; 2 μL injection; 5-100% $CH_3CN/H_2O$, linear-gradient, with constant 0.1% v/v TFA additive; 6 min run; 0.6 mL/min flow; ESI; positive ion mode; UV detection at 254 nm with ACQUITY PDA). If not specially mentioned, flash column chromatography uses silica gel (200-300 mesh) supplied by Tsingtao Haiyang Chemicals (China). Preparative HPLC separations were performed using Teledyne Isco EZ Prep UV-Vis and a RediSep Prep C18 column (100 Å, 5 μm, 20 × 150 mm).

NMR spectra were recorded on Brüker Advance 400 ($^1H$ 400 MHz, $^{13}C$ 101 MHz) and are calibrated using residual undeuterated solvent ($CDCl_3$ at 7.26 ppm $^1H$ NMR, 77.16 ppm $^{13}C$ NMR; $CD_3OD$ at 3.31 ppm $^1H$ NMR, 49.00 ppm $^{13}C$ NMR; $CD_3CN$ at 1.94 ppm $^1H$ NMR, 1.32 and 118.26 ppm $^{13}C$ NMR; DMSO-$d_6$ at 2.50 ppm $^1H$ NMR, 39.52 ppm $^{13}C$ NMR). Data for $^1H$ NMR spectra are reported as follows: chemical shift (δ ppm), multiplicity (s = singlet, d = doublet, t = triplet, q = quartet, dd = doublet of doublets, dt = triplet of doublets, m=multiplet, br=broad), coupling constant (Hz), integration. Data for $^{13}C$ NMR are reported by chemical shift (δ ppm). High-resolution mass spectrometric data were obtained using Acquity I class UPLC synapt G2-SI using ESI (electrospray ionization).

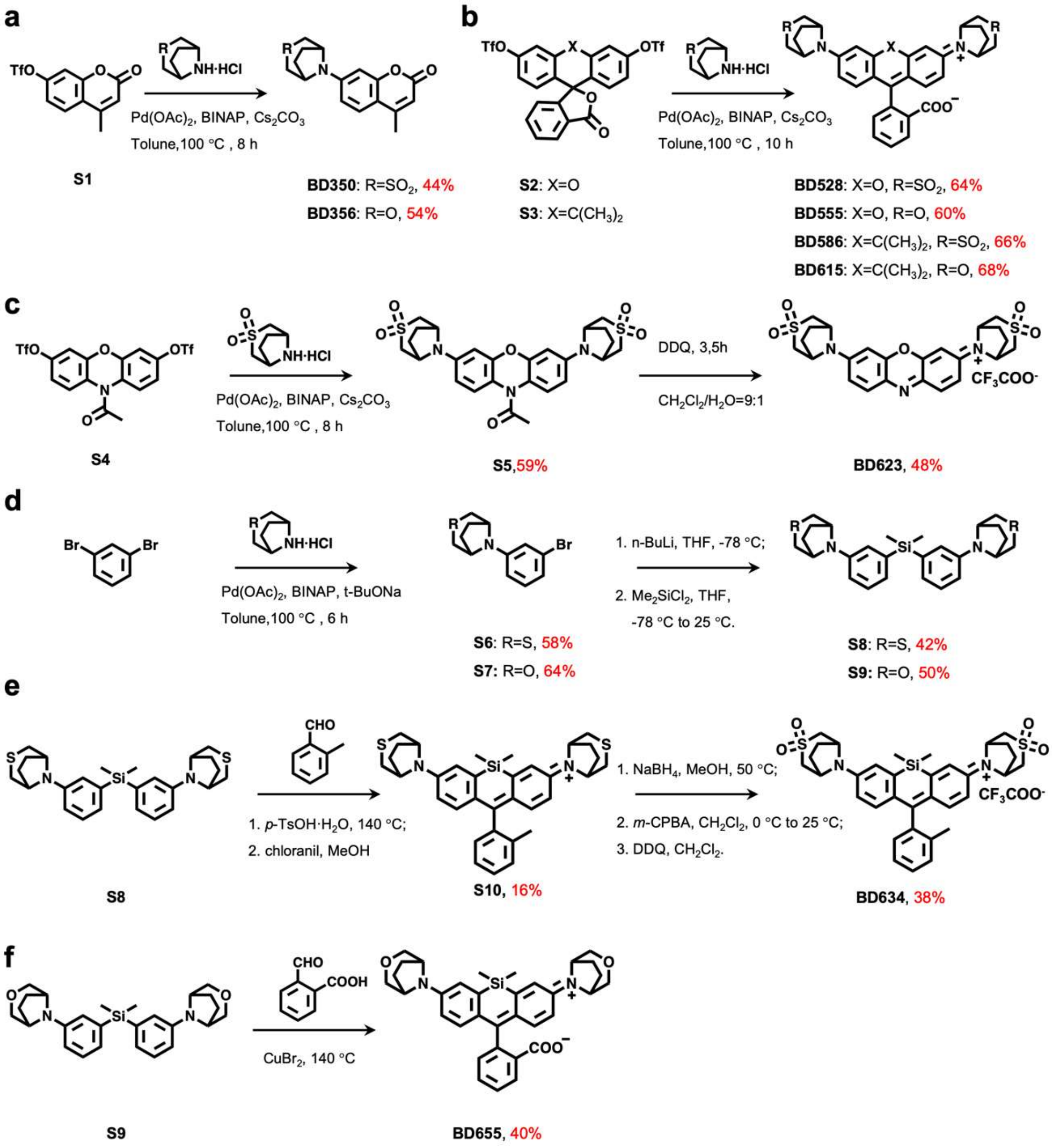


**Scheme S1.** Synthesis of bridged-bicycle strengthened fluorophores.

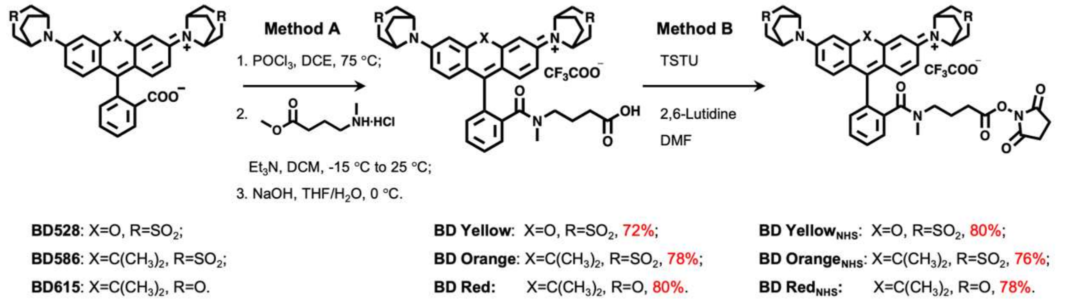


**Scheme S2.** Synthesis of BD derivatives for bioconjugation.

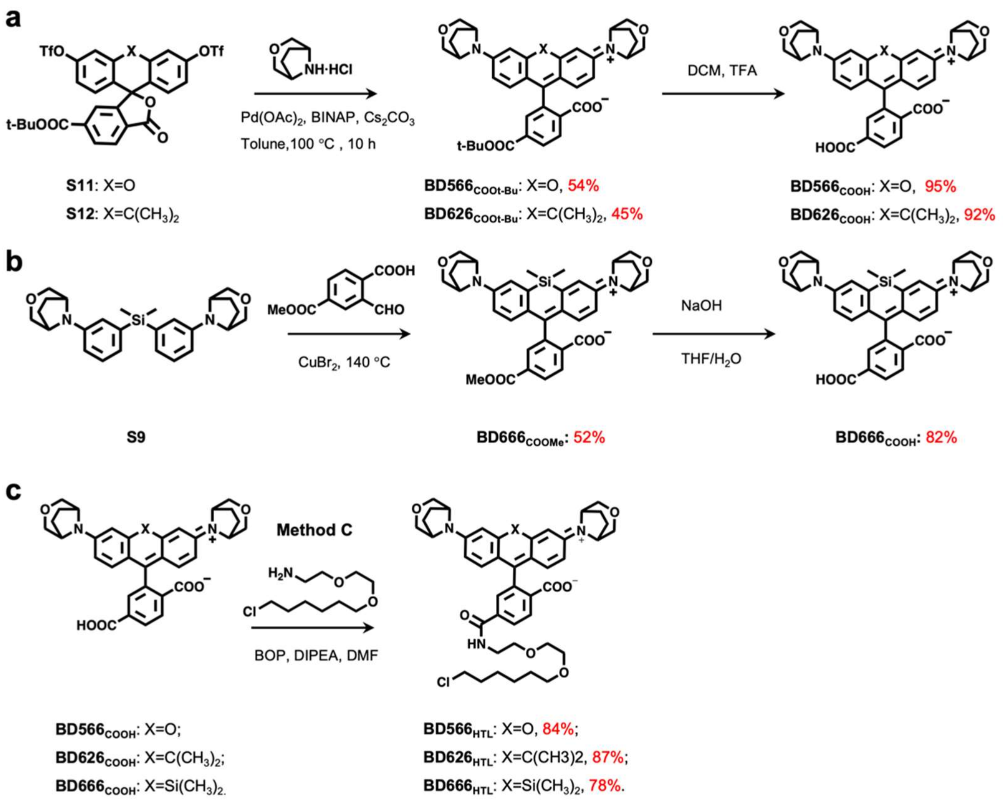


**Scheme S3.** Synthesis of cell-permeant BD derivatives with HaloTag ligands

### Synthetic Procedures and Characterizations

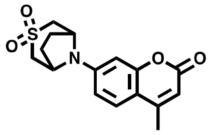

**BD 350: S1** is prepared according to a published protocol[18]. A vial was charged with 4-methylumbelliferone triflate **S1** (120 mg, 390 μmol, 1.0 eq.), $Pd(OAc)_2$ (18 mg, 78 μmol, 0.2 eq.), BINAP (72 mg, 117 μmol, 0.3 eq.), $Cs_2CO_3$ (356 mg, 1092 μmol, 2.8 eq.) and 3-thia-8-azabicyclo [3.2.1] octane 3,3-dioxide hydrochloride (323 mg, 1640 μmol, 4.2 eq.). The vial was sealed and evacuated/backfilled with $N_2$ for three times. Toluene (3 mL) was added, and the reaction was flushed again with $N_2$ for three times. The reaction was stirred at 100 °C for 8 h. After cooling to room temperature, water (10 mL) was added and $CH_2Cl_2$ (10 mL × 3) was used to extract organic compounds. The combined organic phase was dried over $Na_2SO_4$, filtered and concentrated *in vacuo*. Purification of the residue by silica gel chromatography (PE/EA=1/1, v/v) provided **BD 350** (55 mg, 44% yield) as a white solid.

**$^1$H NMR** (400 MHz, DMSO-*d*$_6$) δ 7.63 (d, *J* = 8.8 Hz, 1H), 6.98 (dd, *J* = 8.8, 2.5 Hz, 1H), 6.94 (d, J = 2.4 Hz, 1H), 6.11 – 6.07 (m, 1H), 4.93 – 4.87 (m, 2H), 3.31 – 3.28 (m, 2H), 3.27 – 3.20 (m, 2H), 2.37 (d, *J* = 1.2 Hz, 3H), 2.36 – 2.33 (m, 2H), 2.18 – 2.13 (m, 2H).

**$^{13}$C NMR** (101 MHz, DMSO-*d*$_6$) δ 160.4, 155.3, 153.5, 146.9, 126.9, 111.6, 110.9, 109.8, 101.2, 55.0, 53.1, 26.3, 18.0.

**HRMS** (ESI) calcd for $C_{16}H_{18}NO_4S$ $[M+H]^+$ 320.0957, found 320.0962.

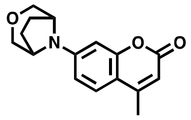

**BD 356: S1** is prepared according to a published protocol[18]. A vial was charged with 4-methylumbelliferone triflate **S1** (120 mg, 390 μmol, 1.0 eq.), $Pd(OAc)_2$ (18 mg, 78 μmol, 0.2 eq.), BINAP (72 mg, 117 μmol, 0.3 eq.), $Cs_2CO_3$ (356 mg, 1092 μmol, 2.8 eq.) and 3-oxa-8-azabicyclo[3.2.1]octane hydrochloride (245 mg, 1640 μmol, 4.2 eq.). The vial was sealed and evacuated/backfilled with $N_2$ for three times. Toluene (3 mL) was added, and the reaction was flushed again with $N_2$ for three times. The reaction was stirred at 100 °C for 8 h. After cooling to room temperature, water (10 mL) was added and $CH_2Cl_2$ (10 mL × 3) was used to extract organic compounds. The combined organic phase was dried over $Na_2SO_4$, filtered and concentrated *in vacuo*. Purification of the residue by silica gel chromatography (PE/EA=1/1, v/v) provided **BD 356** (57 mg, 54% yield) as a white solid.

**$^1$H NMR** (400 MHz, $CDCl_3$) δ 7.43 (d, *J* = 8.8 Hz, 1H), 6.67 (dd, *J* = 8.8, 2.4 Hz, 1H), 6.58 (d, *J* = 2.4 Hz, 1H), 6.07 – 5.96 (m, 1H), 4.17 – 4.11 (m, 2H), 3.89 – 3.81 (m, 2H), 3.59 – 3.52 (m, 2H), 2.36 (d, *J* = 1.2 Hz, 3H), 2.20 – 2.03 (m, 4H).

**$^{13}$C NMR** (101 MHz, $CDCl_3$) δ 162.0, 156.0, 152.9, 149.8, 126.1, 111.7, 111.0, 110.2, 101.6, 69.6, 57.0, 27.1, 18.6.

**HRMS** (ESI) calcd for $C_{16}H_{18}NO_3$ $[M+H]^+$ 272.1287, found 272.1280.

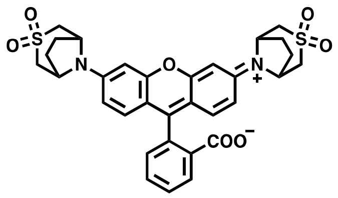

**BD 528: S2** is prepared according to a published protocol[18]. A vial was charged with fluorescein ditriflate **S2** (100 mg, 168 μmol, 1.0 eq.), $Pd(OAc)_2$ (7.5 mg, 34 μmol, 0.2 eq.), BINAP (31 mg, 50 μmol, 0.3 eq.), $Cs_2CO_3$ (153 mg, 469 μmol, 2.8 eq.) and 3-thia-8-azabicyclo [3.2.1] octane 3,3-dioxide hydrochloride (139 mg, 705 μmol, 4.2 eq.). The vial was sealed and evacuated/backfilled with $N_2$ for three times. Toluene (2 mL) was added, and the reaction was flushed again with $N_2$ for three times. The reaction was stirred at 100 °C for 10 h. It was subsequently cooled to room temperature, diluted with MeOH, filtered and concentrated *in vacuo*. Purification of the residue by reverse phase HPLC (eluent, a 30-min linear gradient, from 20% to 95% solvent B; flow rate, 5.0 mL/min; detection wavelength, 530 nm; eluent A ($ddH_2O$ containing 0.1% TFA (v/v)) and eluent B ($CH_3CN$)) provided **BD 528** (66 mg, 64% yield）as a red solid.

**$^1$H NMR** (400 MHz, $CDCl_3$) δ 8.03 (d, $J$ = 7.6 Hz, 1H), 7.71 (td, $J$ = 7.5, 1.2 Hz, 1H), 7.63 (td, $J$ = 7.4, 1.2 Hz, 1H), 7.28 (d, $J$ = 7.6 Hz, 1H), 6.72 (d, $J$ = 8.7 Hz, 2H), 6.55 (d, $J$ = 2.4 Hz, 2H), 6.44 (dd, $J$ = 8.8, 2.4 Hz, 2H), 4.71 – 4.54 (m, 4H), 3.57 – 3.36 (m, 4H), 3.17 – 3.00 (m, 4H), 2.64 – 2.48 (m, 4H), 2.34 – 2.15 (m, 4H).

**$^{13}$C NMR** (101 MHz, DMSO-$d_6$) δ 168.7, 152.4, 152.1, 145.6, 135.6, 130.1, 129.3, 126.4, 124.6, 124.2, 111.5, 108.5, 101.4, 83.3, 69.8, 54.7, 54.6, 53.0, 52.9, 26.4.

**HRMS** (ESI) calcd for $C_{32}H_{31}N_2O_7S_2$ $[M+H]^+$ 619.1567, found 619.1573.

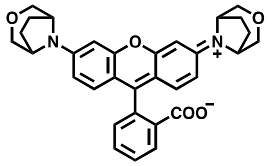

**BD 555: S2** is prepared according to a published protocol[18]. A vial was charged with fluorescein ditriflate **S2** (100 mg, 168 μmol, 1.0 eq.), $Pd(OAc)_2$ (7.5 mg, 34 μmol, 0.2 eq.), BINAP (31 mg, 50 μmol, 0.3 eq.), $Cs_2CO_3$ (153 mg, 469 μmol, 2.8 eq.) and 3-oxa-8-azabicyclo[3.2.1]octane hydrochloride (105 mg, 704 μmol, 4.2 eq.). The vial was sealed and evacuated/backfilled with $N_2$ for three times. Toluene (2 mL) was added, and the reaction was flushed again with $N_2$ for three times. The reaction was stirred at 100 °C for 10 h. It was subsequently cooled to room temperature, diluted with MeOH, filtered and concentrated *in vacuo*. Purification of the residue by reverse phase HPLC (eluent, a 30-min linear gradient, from 20% to 95% solvent B; flow rate, 5.0 mL/min; detection wavelength, 555 nm; eluent A ($ddH_2O$ containing 0.1% TFA (v/v)) and eluent B ($CH_3CN$)) provided **BD 555** (53 mg, 60% yield) as a purple solid.

**$^1$H NMR** (400 MHz, $CDCl_3$) δ 8.03 (d, $J$ = 7.6 Hz, 1H), 7.66 (td, $J$ = 7.5, 1.3 Hz, 1H), 7.60 (td, $J$ = 7.4, 1.1 Hz, 1H), 7.21 (d, $J$ = 7.6 Hz, 1H), 6.64 (d, $J$ = 8.8 Hz, 2H), 6.56 (d, $J$ = 2.4 Hz, 2H), 6.45 (dd, $J$ = 8.8, 2.4 Hz, 2H), 4.15 – 4.02 (m, 4H), 3.93 – 3.81 (m, 4H), 3.59 – 3.45 (m, 4H), 2.19 – 1.89 (m, 8H).

**$^{13}$C NMR** (101 MHz, $CD_3OD$) δ 168.1, 161.2, 159.6, 154.7, 135.3, 133.9, 132.9, 132.6, 132.2, 131.5, 131.5, 116.7, 115.5, 98.9, 72.7, 72.7, 59.2, 27.5.

**HRMS** (ESI) calcd for $C_{32}H_{31}N_2O_5$ $[M+H]^+$ 523.2227, found 523.2256.

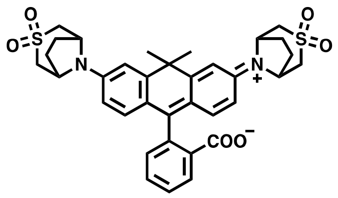

**BD 586: S3** is prepared according to a published protocol[18]. A vial was charged with carbofluorescein ditriflate **S3** (100 mg, 161 µmol, 1.0 eq.), $Pd(OAc)_2$ (7.2 mg, 32 µmol, 0.2 equiv), BINAP (30 mg, 48 µmol, 0.3 equiv), $Cs_2CO_3$ (147 mg, 450 µmol, 2.8 equiv) and 3-thia-8-azabicyclo [3.2.1] octane 3,3-dioxide hydrochloride (133 mg, 676 µmol, 4.2 equiv). The vial was sealed and evacuated/backfilled with $N_2$ for three times. Toluene (2 mL) was added, and the reaction was flushed again with $N_2$ for three times. The reaction was stirred at 100 °C for 10 h. It was subsequently cooled to room temperature, diluted with MeOH, filtered and concentrated *in vacuo*. Purification of the residue by reverse phase HPLC (eluent, a 30-min linear gradient, from 30% to 95% solvent B; flow rate, 5.0 mL/min; detection wavelength, 590 nm; eluent A ($ddH_2O$ containing 0.1% TFA (v/v)) and eluent B ($CH_3CN$)) provided **BD 586** (69 mg, 66% yield) as an off-white solid.

**$^1$H NMR** (400 MHz, $CDCl_3$) δ 8.01 (d, *J* = 7.6 Hz, 1H), 7.65 (td, *J* = 7.5, 1.2 Hz, 1H), 7.58 (td, *J* = 7.4, 1.2 Hz, 1H), 7.13 (d, *J* = 7.6 Hz, 1H), 6.91 (d, *J* = 2.5 Hz, 2H), 6.70 (d, *J* = 8.7 Hz, 2H), 6.55 (dd, *J* = 8.8, 2.4 Hz, 2H), 4.77 – 4.63 (m, 4H), 3.54 – 3.35 (m, 4H), 3.16 – 2.98 (m, 4H), 2.62 – 2.49 (m, 4H), 2.30 – 2.18 (m, 4H), 1.83 (s, 3H), 1.75 (s, 3H).

**$^{13}$C NMR** (101 MHz, $CDCl_3$) δ 170.8, 155.3, 147.1, 143.82, 135.0, 130.3, 129.3, 126.5, 125.1, 124.2, 122.1, 114.0, 111.7, 86.5, 54.7, 54.6, 53.5, 53.4, 38.4, 35.3, 33.5, 27.1, 27.0.

**HRMS** (ESI) calcd for $C_{35}H_{37}N_2O_6S_2$ $[M+H]^+$ 645.2088, found 645.2048.

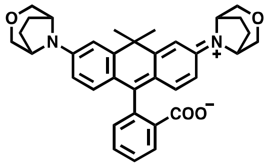

**BD 615: S3** is prepared according to a published protocol[18]. A vial was charged with carbofluorescein ditriflate **S3** (100 mg, 161 µmol, 1.0 eq.), $Pd(OAc)_2$ (7.2 mg, 32 µmol, 0.2 equiv), BINAP (30 mg, 48 µmol, 0.3 equiv), $Cs_2CO_3$ (147 mg, 450 µmol, 2.8 equiv) and 3-oxa-8-azabicyclo[3.2.1]octane hydrochloride (101 mg, 676 µmol, 4.2 equiv). The vial was sealed and evacuated/backfilled with $N_2$ for three times. Toluene (2 mL) was added, and the reaction was flushed again with $N_2$ for three times. The reaction was stirred at 100 °C for 10 h. It was subsequently cooled to room temperature, diluted with MeOH, filtered and concentrated *in vacuo*. Purification of the residue by reverse phase HPLC (eluent, a 30-min linear gradient, from 30% to 95% solvent B; flow rate, 5.0 mL/min; detection wavelength, 620 nm; eluent A ($ddH_2O$ containing 0.1% TFA (v/v)) and eluent B ($CH_3CN$)) provided **BD 615** (60 mg, 68% yield) as a blue solid.

**$^1$H NMR** (400 MHz, $CDCl_3$) δ 7.99 (dd, *J* = 7.0, 1.4 Hz, 1H), 7.60 (td, *J* = 7.4, 1.4 Hz, 1H), 7.55 (td, *J* = 7.4, 1.2 Hz, 1H), 7.08 (d, *J* = 7.6 Hz, 1H), 6.97 (d, *J* = 2.3 Hz, 2H), 6.60 (d, *J* = 8.7 Hz, 2H), 6.54 (dd, *J* = 8.8, 2.4 Hz, 2H), 4.11 – 4.06 (m, 4H), 3.93 – 3.86 (m, 4H), 3.56 – 3.50 (m, 4H), 2.12 – 1.96 (m, 8H), 1.82 (s, 3H), 1.73 (s, 3H).

**$^{13}$C NMR** (101 MHz, $CDCl_3$) δ 170.8, 155.3, 147.8, 147.2, 134.6, 129.5, 129.0, 127.1, 125.0, 124.0, 121.3, 114.4, 112.3, 87.7, 69.8, 69.7, 57.1, 38.5, 35.6, 32.6, 26.9.

**HRMS** (ESI) calcd for $C_{35}H_{37}N_2O_4$ $[M+H]^+$ 549.2748, found 549.2761.

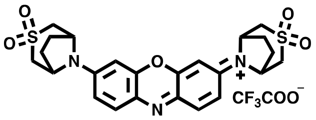


**BD 623: S4** is prepared according to a published protocol[18]. A vial was charged with ditriflate **S4** (150 mg, 288 μmol, 1.0 eq.), $Pd(OAc)_2$ (13 mg, 57.6 μmol, 0.2 equiv), BINAP (54 mg, 86.4 μmol, 0.3 equiv), $Cs_2CO_3$ (263 mg, 806 μmol, 2.8 eq.) and 3-thia-8-azabicyclo [3.2.1] octane 3,3-dioxide hydrochloride (238 mg, 1210 μmol, 4.2 eq.). The vial was sealed and evacuated/backfilled with $N_2$ for three times. Toluene (3 mL) was added, and the reaction was flushed again with $N_2$ for three times. The reaction was stirred at 100 °C for 8 h. It was subsequently cooled to room temperature, diluted with dichloromethane. The combined organic phase was washed with brine, dried over $Na_2SO_4$, filtered and concentrated *in vacuo*. Purification by silica gel chromatography (PE/ EA =2/3, v/v) afforded the *N*-acetyl leuco-dye **S5** as a colorless solid (93 mg, 59%). The intermediate **S5** (93 mg, 171 μmol) was taken up in a mixture of dichloromethane (10.0 mL) and water (1.1 mL) and cooled to 0 °C. DDQ (43 mg, 188 μmol) was added, and the reaction was stirred at room temperature for 3 h. A second portion of DDQ (21 mg, 94 μmol) was added, and the reaction was stirred for an additional 0.5 h. The mixture was evaporated, redissolved in $CH_3CN$, filtered and concentrated to dryness. Purification of the residue by reverse phase HPLC (eluent, a 30-min linear gradient, from 10% to 95% solvent B; flow rate, 5.0 mL/min; detection wavelength, 630 nm; eluent A ($ddH_2O$ containing 0.1% TFA (v/v)) and eluent B ($CH_3CN$)) provided **BD 623** (41 mg, 48% yield, TFA salt) as a blue solid.

**$^1$H NMR** (400 MHz, DMSO-$d_6$) δ 8.01 (d, *J* = 9.4 Hz, 2H), 7.67 (dd, *J* = 9.5, 2.4 Hz, 2H), 7.34 (d, *J* = 2.5 Hz, 2H), 5.38 – 5.34 (m, 4H), 3.65 – 3.57 (m, 4H), 3.53 – 3.45 (m, 4H), 2.53 – 2.40 (m, 4H), 2.29 – 2.21 (m, 4H).

**$^{13}$C NMR** (101 MHz, DMSO-$d_6$) δ 153.5, 149.3, 135.2, 134.7, 119.7, 98.4, 58.2, 55.1, 25.8.

**HRMS** (ESI) calcd for $C_{24}H_{26}N_3O_5S_2$ $[M]^+$ 500.1308, found 500.1318.

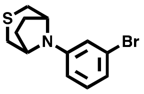

**S6:**A vial was charged with 1,3-dibromobenzene (7.5 g, 32 mmol, 1.0 eq.), $Pd(OAc)_2$ (1.4 g, 6.4 mmol, 0.2 eq.), BINAP (6.0 g, 9.6 mmol, 0.3 eq.), t-BuONa (6.2 g, 64 mmol, 2.0 eq.) and 3-thia-8-azabicyclo [3.2.1] octane hydrochloride (6.3 g, 38 mmol, 1.2 equiv). The vial was sealed and evacuated/backfilled with $N_2$ for three times. Toluene (30 mL) was added, and the reaction was flushed again with $N_2$ for three times. The reaction mixture was stirred at 100 °C for 6 h. After cooling to room temperature, water (20 mL) was added and $CH_2Cl_2$ (20 mL × 3) was used to extract organic compounds. The combined organic phase was dried over $Na_2SO_4$, filtered and concentrated *in vacuo*. Purification of the residue by silica gel chromatography (PE/EA=50/1, v/v) provided compound **S6** (5.3 g, 58% yield) as a white solid.

**$^1$H NMR** (400 MHz, $CDCl_3$) δ 7.08 (t, *J* = 8.1 Hz, 1H), 6.87 – 6.78 (m, 2H), 6.62 (dd, *J* = 8.3, 2.4 Hz, 1H), 4.44 – 4.32 (m, 2H), 3.41 – 3.30 (m, 2H), 2.31 – 2.12 (m, 4H), 2.00 – 1.84 (m, 2H).

**$^{13}$C NMR** (101 MHz, $CDCl_3$) δ 146.6, 131.1, 124.0, 119.7, 117.6, 113.4, 54.2, 28.8, 27.4.

**HRMS** (ESI) calcd for $C_{12}H_{15}BrNS$ $[M+H]^+$ 284.0103, found 284.0105.

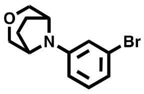

**S7:**A vial was charged with 1,3-dibromobenzene (7.5 g, 32 mmol, 1.0 eq.), $Pd(OAc)_2$ (1.4 g, 6.4 mmol, 0.2 eq.), BINAP (6.0 g, 9.6 mmol, 0.3 eq.), t-BuONa (6.2 g, 64 mmol, 2.0 eq.) and 3-oxa-8-azabicyclo[3.2.1]octane hydrochloride (5.7 g, 38 mmol, 1.2 equiv). The vial was sealed and evacuated/backfilled with $N_2$ for three times. Toluene (30 mL) was added, and the reaction was flushed again with $N_2$ for three times. The reaction mixture was stirred at 100 °C for 6 h. After cooling to room temperature, water (20 mL) was added and $CH_2Cl_2$ (20 mL × 3) was used to extract organic compounds. The combined organic phase was dried over $Na_2SO_4$, filtered and concentrated *in vacuo*. Purification of the residue by silica gel chromatography (PE/EA=50/1, v/v) provided compound **S7** (5.5 g, 64% yield) as a white solid.
**$^1$H NMR** (400 MHz, $CDCl_3$) δ 7.08 (t, *J* = 8.1 Hz, 1H), 6.92 – 6.84 (m, 2H), 6.68 (dd, *J* = 8.4, 2.4 Hz, 1H), 4.04 – 3.97 (m, 2H), 3.92 – 3.84 (m, 2H), 3.57 – 3.47 (m, 2H), 2.14 – 1.94 (m, 4H).
**$^{13}$C NMR** (101 MHz, $CDCl_3$) δ 148.7, 131.0, 123.8, 120.9, 118.1, 113.9, 69.8, 57.8, 26.9.
**HRMS** (ESI) calcd for $C_{12}H_{15}BrNO$ $[M+H]^+$ 268.0332, found 268.0338.

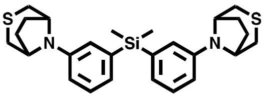

**S8**:To a flame-dried flask flushed with nitrogen, compound **S6** (3.7 g, 13 mmol, 1.0 equiv) and anhydrous THF (9 mL) were added. The solution was cooled to -78 °C, n-BuLi (8.8 mL of 1.6 M solution in hexane, 14 mmol, 1.1 equiv) was injected with a syringe quickly dropwise, and the mixture was stirred at -78 °C for 20 min. At the same temperature, $SiMe_2Cl_2$ (0.78 mL, 6.5 mmol, 0.5 equiv) was injected with a syringe dropwisely, and the mixture was warmed to room temperature, then stirred for 30 min. It was subsequently quenched with saturated $NH_4Cl$, diluted with water, and extracted with EtOAc. The organic phase was collected and washed with brine, dried over $Na_2SO_4$, filtered, and concentrated *in vacuo*. The residue was purified by silica gel column chromatography (PE / EA = 20/1, v/v) to afford product **S8** as a white solid (1.3 g, 42%).
**$^1$H NMR** (400 MHz, CDCl3) δ 7.28 – 7.23 (m, 2H), 7.03 – 6.81 (m, 4H), 6.83 – 6.58 (m, 2H), 4.46 – 4.39 (m, 4H), 3.47 – 3.31 (m, 4H), 2.30 – 2.12 (m, 8H), 1.98 – 1.77 (m, 4H), 0.52 (s, 6H).
**$^{13}$C NMR** (101 MHz, CDCl3) δ 144.6, 139.8, 129.3, 123.0, 120.8, 116.0, 53.9, 28.8, 27.0,-2.1.
HRMS (ESI) calcd for $C_{26}H_{35}N_2S_2Si$ [M+H]+ 467.2005, found 467.2019.

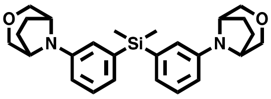

**S9**:To a flame-dried flask flushed with nitrogen, compound **S7** (3.5 g, 13 mmol, 1.0 equiv) and anhydrous THF (9 mL) were added. The solution was cooled to -78 °C, n-BuLi (8.8 mL of 1.6 M solution in hexane, 14 mmol, 1.1 equiv) was injected with a syringe quickly dropwise, and the mixture was stirred at -78 °C for 20 min. At the same temperature, $SiMe_2Cl_2$ (0.78 mL, 6.5 mmol, 0.5 equiv) was injected with a syringe dropwisely, and the mixture was warmed to room temperature, then stirred for 30 min. It was subsequently quenched with saturated $NH_4Cl$, diluted with water, and extracted with EtOAc. The organic phase was collected and washed with brine,

dried over $Na_2SO_4$, filtered, and concentrated *in vacuo*. The residue was purified by silica gel column chromatography (PE / EA = 20/1, v/v) to afford product **S9** as a white solid (1.3 g, 42%).
**$^1$H NMR** (400 MHz, $CDCl_3$) δ 7.26 – 7.22 (m, 2H), 7.01 – 6.90 (m, 4H), 6.88 – 6.77 (m, 2H), 4.09 – 3.98 (m, 4H), 3.98 – 3.83 (m, 4H), 3.56 – 3.42 (m, 4H), 2.10 – 2.01 (m, 4H), 1.99 – 1.91 (m, 4H), 0.51 (s, 6H).
**$^{13}$C NMR** (101 MHz, $CDCl_3$) δ 146.8, 139.6, 129.1, 124.4, 121.2,116.4, 70.0, 57.3, 26.7,-2.1.
**HRMS** (ESI) calcd for $C_{26}H_{35}N_2S_2Si$ $[M+H]^+$ 435.2462, found 435.2477.

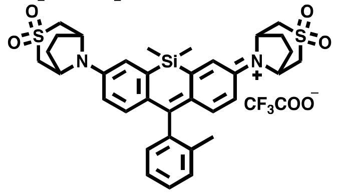


**BD 634**: Compound **S8** (150 mg, 0.32 mmol, 1.0 eq.), *o*-tolualdehyde (190 mg, 1.6 mmol, 5.0 eq.) and *p*-TsOH·$H_2O$ (61 mg, 0.32 mmol, 1.0 eq.) were mixed in a sealable pressure tube. The tube was sealed tightly and heated at 140 °C for 8 h. After cooling to room temperature, the mixture was diluted with MeOH (3 mL), then added chloranil (79 mg, 0.32 mmol, 1.0 eq.) and stirred for 2 h. After filtration and removal of the solvent, the residue was purified by silica gel column chromatography ($CH_2Cl_2$/MeOH=15/1, v/v, with constant 1% v/v AcOH additive) to afford intermediate **S10** as blue-green solid (39 mg, 16%).

The intermediate **S10** (39 mg, 69 mmol) was taken up in MeOH (5 mL) and added $NaBH_4$ (5.2 mg, 140 mmol). The reaction mixture was stirred for 0.5 h at 50°C. After cooling to room temperature, the mixture was diluted with $H_2O$ and then extracted with $CH_2Cl_2$ (3×). The combined organic layers were dried over $Na_2SO_4$, concentrated *in vacuo* and directly used for the next reaction. The residue was dissolved in $CH_2Cl_2$ (2 mL) and a solution of 3-chloroperozybenzoic acid (126 mg, 550 mmol) in $CH_2Cl_2$ (2 mL) was added dropwisely at 0 °C. The reaction mixture was stirred for 2 h at 0 °C, then allowed to warm to room temperature for an additional 2h. After that, the mixture was washed with aq. $NaHSO_3$ and extracted with $CH_2Cl_2$. The collected organic layers were dried over $Na_2SO_4$ and concentrated *in vacuo*. The residue was again dissolved in $CH_2Cl_2$ (2 mL), and DDQ (15 mg, 69 mmol) was added in one portion. The mixture was stirred for 2 h, and then washed with brine, dried over $Na_2SO_4$, filtered, and concentrated *in vacuo*. The resulting residue was redissolved in MeOH (1 mL), and AcOH (10 μL) was added (immediate blue color). After stirring the blue solution at room temperature for 10 min, the mixture was concentrated to dryness and purified by HPLC (eluent, a 30-min linear gradient, from 10% to 95% solvent B; flow rate, 5.0 mL/min; detection wavelength, 630 nm; eluent A ($ddH_2O$ containing 0.1% TFA (v/v)) and eluent B ($CH_3CN$)) provided **BD 634** (16 mg, 38% yield, TFA salt) as a blue solid.
**$^1$H NMR** (400 MHz, DMF-$d_7$) δ 7.96 (d, $J$ = 2.5 Hz, 2H), 7.61 – 7.45 (m, 3H), 7.30 (dd, $J$ = 7.6, 1.3 Hz, 1H), 7.25 – 7.15 (m, 4H), 5.65 – 5.30 (m, 4H), 3.67 – 3.53 (m, 8H), 2.61 – 2.52 (m, 4H), 2.37 – 2.27 (m, 4H), 2.10 (s, 3H), 0.70 (s, 3H), 0.69 (s, 3H).
**$^{13}$C NMR** (101 MHz, $CD_3CN$) δ 173.5, 151.5, 151.1, 143.8, 139.2, 136.6, 131.2, 130.2, 130.0, 129.7, 126.6, 123.5, 116.8, 59.0, 55.3, 26.8, 19.5, -1.4, -1.5.
**HRMS** (ESI) calcd for $C_{34}H_{39}N_2O_4S_2Si$ $[M]^+$ 631.2115, found 631.2107.

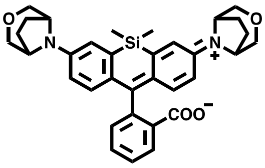

**BD 655**: Compound **S9** (139 mg, 0.32 mmol, 1.0 eq.), 2-formylbenzoic acid (240 mg, 1.6 mmol, 5.0 eq.) and $CuBr_2$ (7 mg, 0.03 mmol, 0.1 eq.) were mixed in a sealable pressure tube. The tube was sealed tightly and heated at 140 °C for 4 h. After cooling to room temperature, the reaction mixture was dissolved in MeOH (5 mL), filtered, and concentrated *in vacuo*. The residue was purified by silica gel column chromatography (PE / EA = 5/2, v/v) to afford product **BD 655** as a blue-green solid (72 mg, 40%).
**$^1$H NMR** (400 MHz, $CDCl_3$) δ 7.97 (d, *J* = 7.6 Hz, 1H), 7.66 (td, *J* = 7.5, 1.2 Hz, 1H), 7.56 (td, *J* = 7.5, 1.0 Hz, 1H), 7.34 (d, *J* = 7.6 Hz, 1H), 7.04 (d, *J* = 2.7 Hz, 2H), 6.79 (d, *J* = 8.7 Hz, 2H), 6.59 (dd, *J* = 8.9, 2.8 Hz, 2H), 4.11 – 4.05 (m, 4H), 3.91 – 3.84 (m, 4H), 3.56 – 3.46 (m, 4H), 2.12 – 1.99 (m, 4H), 2.02 – 1.93 (m, 4H), 0.61 (s, 3H), 0.60 (s, 3H).
**$^{13}$C NMR** (101 MHz, $CDCl_3$) δ 170.7, 154.0, 146.5, 137.7, 133.8, 133.7, 129.0, 128.7, 127.1, 125.9, 124.8, 119.9, 116.2, 91.6, 69.9, 69.8, 57.0, 56.9, 26.9, 26.9, 0.5, -1.5.
**HRMS** (ESI) calcd for $C_{34}H_{37}N_2O_4Si$ $[M+H]^+$ 565.2517, found 565.2528.

**General Method A**: **Functionalization with carboxylic acid**
Representative Procedure for **BD Yellow**:

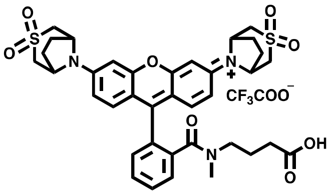

**BD Yellow**: To a flame-dried flask flushed with nitrogen, **BD528** (24 mg, 0.038 mmol, 1.0 eq.) and anhydrous 1,2-dichloroethane (2 mL) were added. Then $POCl_3$ (0.18 mL, 1.9 mmol, 50.0 eq.) was added by a syringe. The reaction was stirred at 75 °C for 2 h. After cooling to room temperature, the mixture was concentrated *in vacuo*. $CH_2Cl_2$ (2 mL) was added and removed by rotary evaporation, and the mixture was placed under high vacuum (25 mmHg) for 0.5 h. To the round-bottomed flask, anhydrous $CH_2Cl_2$ (1 mL) was added by a syringe and the reaction was placed in an ice bath (–15 °C). Meanwhile, methyl 4-(methylamino)butanoate hydrochloride (63 mg, 0.38 mmol, 10.0 eq.) and triethylamine (0.05 mL) was dissolved in $CH_2Cl_2$ (1 mL), giving amine solution. After the reaction mixture had stirred for 10 minutes in the external bath, the free-based amine was added via syringe in one addition. The reaction was stirred at the same temperature for another 20 minutes and then allowed to warm to 25 °C and stirred for an additional 20 minutes. 1 N HCl (2 mL) was added by syringe and the two layers separated. The aqueous layer was further extracted with $CH_2Cl_2$ (2 x 3 mL). The organic layers were combined, washed with brine and dried over $Na_2SO_4$. The solvent was removed by rotary evaporation and the crude methyl ester was taken directly to the next reaction. The crude methyl ester **24** (0.038 mmol) was added into a 25 mL flask. THF (5 mL) was added by a syringe followed by dd$H_2O$ (2 mL). The reaction was stirred at 0 °C for 10 min. A solution of 1 N NaOH (0.19 mL, 0.19 mmol, 5.0 eq.) was added and the reaction was stirred at this same temperature for 2 h before acidified with glacial AcOH (0.5 mL, 8.75 mmol, 10.0 eq). The reaction was concentrated *in vacuo*. Purification of the residue by HPLC (eluent, a 30-min linear gradient, from 10% to 90% solvent B; flow rate, 5.0 mL/min;

detection wavelength, 540 nm; eluent A (dd$H_2O$ containing 0.1% TFA (v/v)) and eluent B ($CH_3CN$)) provided **BD Yellow** (19 mg, 72% yield) as a red solid.

**$^1$H NMR** (400 MHz, $CD_3CN$) δ 7.77 – 7.60 (m, 3H), 7.51 – 7.29 (m, 3H), 7.18 – 7.03 (m, 4H), 5.07 – 5.01 (m, 4H), 3.75 – 3.30 (m, 8H), 3.24 – 3.13 (m, 2H), 2.91 (s, 3H), 2.58 – 2.48 (m, 4H), 2.37 – 2.16 (m, 4H), 1.78 – 1.69 (m, 2H), 1.38 – 1.25 (m, 2H).

**$^{13}$C NMR** (101 MHz, $CD_3CN$) δ 174.6, 169.0, 159.3, 154.1, 137.7, 134.4, 134.0, 131.1, 131.0, 130.5, 128.4, 126.7, 116.7, 116.3, 99.7, 58.8, 55.6, 55.6, 46.7, 37.7, 31.0, 27.1, 22.3.

**HRMS** (ESI) calcd for $C_{37}H_{40}N_3O_8S_2$ $[M]^+$ 718.2251, found 718.2253.

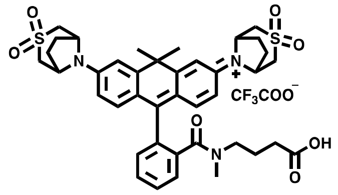


**BD Orange**: The title compound (25 mg, 78%, purple solid) was prepared from **BD 586** according to general method A.

**$^1$H NMR** (400 MHz, Acetonitrile-$d_3$) δ 7.72 – 7.65 (m, 2H), 7.63 – 7.58 (m, 1H), 7.46 – 7.39 (m, 1H), 7.32 – 7.23 (m, 4H), 6.90 (dd, $J$ = 9.3, 2.4 Hz, 2H), 5.11 (s, 4H), 3.49 – 3.30 (m, 8H), 3.24 – 3.16 (m, 2H), 2.92 (s, 3H), 2.58 – 2.45 (m, 4H), 2.30 – 2.22 (m, 4H), 1.83 (s, 3H), 1.81 – 1.73 (m, 2H), 1.71 (s, 3H), 1.41 – 1.30 (m, 2H).

$^{13}$C NMR (101 MHz, $CD_3CN$) δ 174.6, 169.3, 167.3, 159.7, 153.1, 140.1, 137.3, 135.1, 131.1, 130.3, 130.1, 128.1, 123.2, 115.2, 113.8, 59.1, 59.0, 55.4, 46.9, 43.1, 37.7, 34.8, 32.4, 31.2, 27.0, 26.9, 22.5.

**HRMS** (ESI) calcd for $C_{40}H_{46}N_3O_7S_2$ $[M]^+$ 744.2772, found 744.2790.

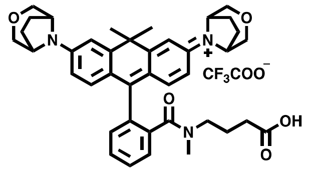


**BD Red**: The title compound (28 mg, 80%, blue solid) was prepared from **BD 615** according to general method A.

**$^1$H NMR** (400 MHz, $CD_3CN$) δ 7.67 – 7.62 (m, 2H), 7.59 – 7.49 (m, 1H), 7.42 – 7.34 (m, 1H), 7.17 – 7.14 (m, 2H), 7.14 – 7.08 (m, 2H), 6.79 – 6.72 (m, 2H), 4.65 – 4.62 (m, 4H), 3.72 – 3.65 (m, 8H), 3.26 – 3.14 (m, 2H), 2.88 (s, 3H), 2.18 – 2.10 (m, 4H), 2.12 – 2.04 (m, 4H), 1.83 – 1.78 (m, 2H), 1.77 (s, 3H), 1.66 (s, 3H), 1.45 – 1.37 (m, 2H).

**$^{13}$C NMR** (101 MHz, $CD_3CN$) δ 174.4, 169.5, 163.0, 158.2, 152.8, 139.1, 137.3, 135.4, 131.2, 130.0, 129.9, 127.9, 121.9, 114.6, 113.4, 72.8, 58.8, 58.7, 46.9, 42.6, 37.8, 31.4, 27.1, 27.0, 22.6.

**HRMS** (ESI) calcd for $C_{40}H_{46}N_3O_5$ $[M]^+$ 648.3432, found 648.3440.

**General Method B: Conversion of carboxylic acids to NHS esters**

Representative Procedure for **BD Yellow$_{\mathbf{NHS}}$**:

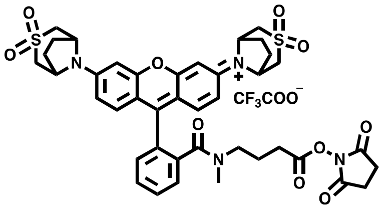


**BD Yellow (16)** (10 mg, 14 μmol, 1.0 eq.), TSTU (8.6 mg, 28 μmol, 2.0 eq.) and 2,6-Lutidine (15 mg, 140 μmol, 10.0 eq.) were dissolved in anhydrous DMF (1 mL). The mixture was stirred at room temperature for 4 h. The product was purified by RP-HPLC (eluent, a 20-min linear gradient, from 20% to 80% solvent B; flow rate, 5.0 mL/min; detection wavelength, 540 nm; eluent A (dd$H_2O$ containing 0.1% TFA (v/v)) and eluent B ($CH_3CN$)) and freeze-dried to yield **BD Yellow$_{\mathbf{NHS}}$** (8 mg, 80% yield) as a red solid.

**Analytical HPLC:** 97.6% purity (4.6 mm × 150 mm 5 μm C18 column; 2 μL injection; 5-100% $CH_3CN/H_2O$, linear-gradient, with constant 0.1% v/v TFA additive; 6 min run; 0.6 mL/min flow; ESI; positive ion mode; detection at 600 nm).

**HRMS** (ESI) calcd for $C_{41}H_{43}N_4O_{10}S_2$ $[M]^+$ 815.2415, found 815.2427.

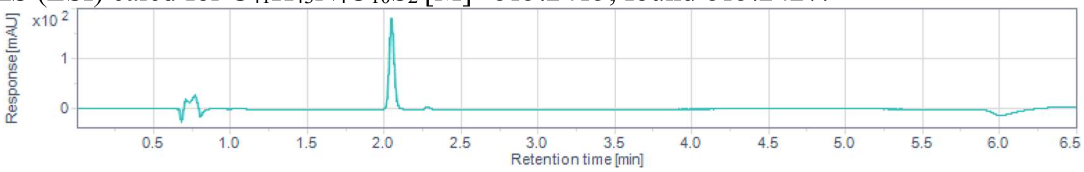


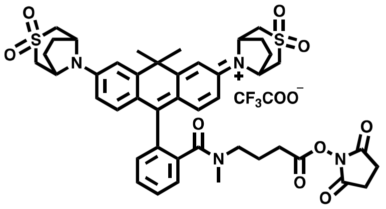


**BD Orange$_{\mathbf{NHS}}$**: The title compound (8.8 mg, 76%, purple solid) was prepared from **BD Orange (17)** according to general method B.

**Analytical HPLC:** 98.4% purity (4.6 mm × 150 mm 5 μm C18 column; 2 μL injection; 5-100% $CH_3CN/H_2O$, linear-gradient, with constant 0.1% v/v TFA additive; 6 min run; 0.6 mL/min flow; ESI; positive ion mode; detection at 600 nm).

**HRMS** (ESI) calcd for $C_{44}H_{49}N_4O_9S_2$ $[M]^+$ 841.2935, found 841.2962.

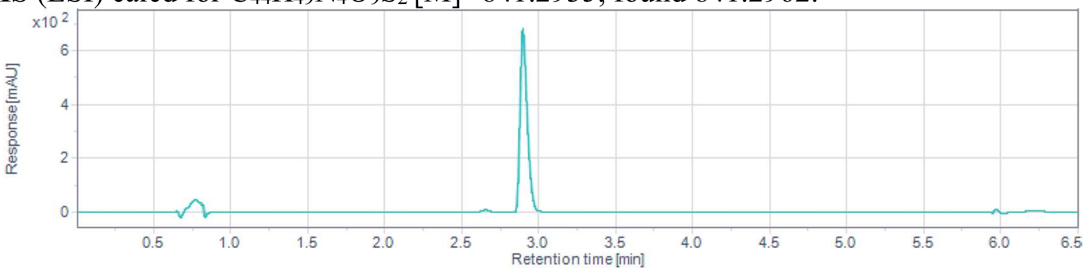

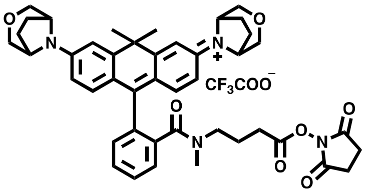

**BD Red$_{NHS}$**: The title compound (9.0 mg, 78%, blue solid) was prepared from **BD Red** according to general method B.

**Analytical HPLC:** 96.8% purity (4.6 mm × 150 mm 5 µm C18 column; 2 µL injection; 5-100% $CH_3CN/H_2O$, linear-gradient, with constant 0.1% v/v TFA additive; 6 min run; 0.6 mL/min flow; ESI; positive ion mode; detection at 630 nm).

**HRMS** (ESI) calcd for $C_{44}H_{49}N_4O_7$ $[M]^+$ 745.3596, found 745.3582.

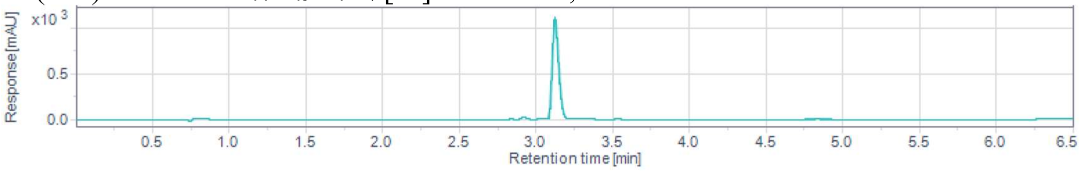


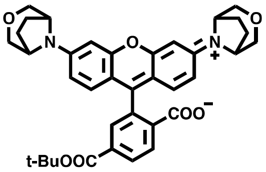

**BD 566$_{COOt\text{-}Bu}$: S11** is prepared according to a published protocol[18]. A vial was charged with ditriflate **S11** (100 mg, 144 µmol, 1.0 eq.), $Pd(OAc)_2$ (6.4 mg, 29 µmol, 0.2 eq.), BINAP (27 mg, 43 µmol, 0.3 eq.), $Cs_2CO_3$ (131 mg, 403 µmol, 2.8 eq.) and 3-oxa-8-azabicyclo[3.2.1]octane hydrochloride (90 mg, 605 µmol, 4.2 eq.). The vial was sealed and evacuated/backfilled with $N_2$ for three times. Toluene (2 mL) was added, and the reaction was flushed again with $N_2$ for three times. The reaction was stirred at 100 °C for 10 h. It was subsequently cooled to room temperature, diluted with MeOH, filtered and concentrated *in vacuo*. Purification of the residue by silica gel chromatography ($CH_2Cl_2$/MeOH (2M $NH_3$)=20/1, v/v) provided **BD 566$_{COOt\text{-}Bu}$** (48 mg, 54% yield) as a purple solid.

**$^1$H NMR** (400 MHz, $CDCl_3$) δ 8.19 (dd, $J$ = 8.0, 1.4 Hz, 1H), 8.03 (d, $J$ = 8.0 Hz, 1H), 7.80 – 7.75 (m, 1H), 6.63 – 6.53 (m, 4H), 6.48 – 6.41 (m, 2H), 4.09 – 4.05 (m, 4H), 3.92 – 3.84 (m, 5H), 3.56 – 3.49 (m, 4H), 2.16 – 1.96 (m, 8H), 1.55 (s, 9H).

**$^{13}$C NMR** (101 MHz, $CDCl_3$) δ 169.0, 164.5, 153.3, 149.2, 138.0, 130.8, 130.7, 129.6, 125.4, 125.4, 125.0, 124.9, 111.7, 107.9, 101.7, 82.5, 69.7, 69.6, 57.0, 56.9, 28.2, 27.0.

**HRMS** (ESI) calcd for $C_{37}H_{39}N_2O_7$ $[M+H]^+$ 623.2752, found 623.2764.

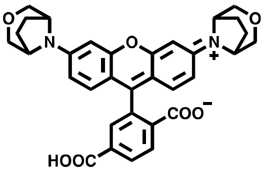

**BD 566$_{COOH}$**: **BD 566$_{COOt\text{-}Bu}$** (48 mg, 77 mmol) was dissolved in $CH_2Cl_2$ (1.5 mL), and trifluoroacetic acid (0.3 mL) was added. The reaction was stirred at room temperature for 2 h and then concentrated *in vacuo*. Purification of the residue by reverse phase HPLC (eluent, a 30-min linear gradient, from 30% to 95% solvent B; flow rate, 5.0 mL/min; detection wavelength, 560 nm; eluent A ($ddH_2O$ containing 0.1% TFA (v/v)) and eluent B ($CH_3CN$)) provided **BD 566$_{\mathbf{COOH}}$** (41 mg, 95% yield) as a purple solid.

**$^1$H NMR** (400 MHz, $CD_3OD$) δ 8.46 – 8.36 (m, 2H), 8.02 – 7.97 (m, 1H), 7.17 – 7.05 (m, 6H), 4.67 – 4.61 (m, 4H), 3.81 – 3.68 (m, 8H), 2.26 – 2.09 (m, 8H).

**$^{13}$C NMR** (101 MHz, $CD_3OD$) δ 167.7, 167.4, 159.7, 159.7, 154.7, 136.0, 135.9, 135.4, 132.9, 132.7, 132.4, 132.3, 116.9, 115.4, 99.0, 72.8, 59.3, 27.5.

**HRMS** (ESI) calcd for $C_{33}H_{31}N_2O_7$ $[M+H]^+$ 567.2126, found 567.2148.

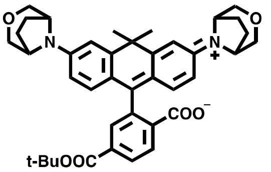


**BD 626$_{\mathbf{COOt\text{-}Bu}}$: S12** is prepared according to a published protocol[18]. A vial was charged with ditriflate **S12** (104 mg, 144 μmol, 1.0 eq.), $Pd(OAc)_2$ (6.4 mg, 29 μmol, 0.2 eq.), BINAP (27 mg, 43 μmol, 0.3 eq.), $Cs_2CO_3$ (131 mg, 403 μmol, 2.8 eq.) and 3-oxa-8-azabicyclo[3.2.1]octane hydrochloride (90 mg, 605 μmol, 4.2 eq.). The vial was sealed and evacuated/backfilled with $N_2$ for three times. Toluene (2 mL) was added, and the reaction was flushed again with $N_2$ for three times. The reaction was stirred at 100 °C for 6 h. It was subsequently cooled to room temperature, diluted with MeOH, filtered and concentrated *in vacuo*. Purification of the residue by silica gel chromatography ($CH_2Cl_2$/MeOH=40/1, v/v) provided **BD 626$_{\mathbf{COOt\text{-}Bu}}$** (42 mg, 45% yield) as a blue solid.

**$^1$H NMR** (400 MHz, $CDCl_3$) δ 8.15 (dd, $J$ = 8.0, 1.4 Hz, 1H), 8.02 (d, $J$ = 8.0 Hz, 1H), 7.66 – 7.61 (m, 1H), 6.96 (d, $J$ = 2.2 Hz, 2H), 6.60 – 6.51 (m, 4H), 4.14 – 4.03 (m, 4H), 3.95 – 3.83 (m, 4H), 3.58 – 3.47 (m, 4H), 2.13 – 1.95 (m, 8H), 1.82 (s, 3H), 1.73 (s, 3H), 1.54 (s, 9H).

**$^{13}$C NMR** (101 MHz, $CDCl_3$) δ 170.1, 164.5, 155.5, 147.9, 147.1, 137.9, 130.2, 130.1, 129.5, 125.1, 124.9, 120.6, 114.5, 112.4, 88.0, 82.5, 69.9, 69.8, 57.1, 57.0, 38.5, 35.4, 33.1, 28.2, 26.9.

**HRMS** (ESI) calcd for $C_{40}H_{45}N_2O_6$ $[M+H]^+$ 649.3272, found 649.3291.

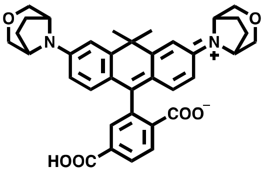


**BD 626$_{\mathbf{COOH}}$**:**BD 626$_{\mathbf{COOt\text{-}Bu}}$** (42 mg, 65 mmol) was dissolved in $CH_2Cl_2$ (1.5 mL), and trifluoroacetic acid (0.3 mL) was added. The reaction was stirred at room temperature for 2 h and then concentrated *in vacuo*. Purification of the residue by reverse phase HPLC (eluent, a 30-min linear gradient, from 30% to 95% solvent B; flow rate, 5.0 mL/min; detection wavelength, 630 nm; eluent A ($ddH_2O$ containing 0.1% TFA (v/v)) and eluent B ($CH_3CN$)) provided **BD 626$_{\mathbf{COOH}}$** (35 mg, 92% yield) as a blue solid.

**$^1$H NMR** (400 MHz, DMF-$d_7$) δ 8.31 (dd, $J$ = 8.0, 1.4 Hz, 1H), 8.21 (d, $J$ = 8.0 Hz, 1H), 7.72 – 7.66 (m, 1H), 7.36 – 7.28 (m, 2H), 6.83 (dd, $J$ = 8.9, 2.4 Hz, 2H), 6.69 (d, $J$ = 8.8 Hz, 2H), 4.50 – 4.27 (m, 4H), 3.83 – 3.74 (m, 4H), 3.58 – 3.46 (m, 4H), 2.07 – 1.94 (m, 8H), 1.92 (s, 3H), 1.80 (s, 3H).

**Analytical HPLC:** >99% purity (4.6 mm × 150 mm 5 μm C18 column; 2 μL injection; 5-100% $CH_3CN/H_2O$, linear-gradient, with constant 0.1% v/v TFA additive; 6 min run; 0.6 mL/min flow; ESI; positive ion mode; detection at 630 nm).
**HRMS** (ESI) calcd for $C_{36}H_{37}N_2O_6$ $[M+H]^+$ 593.2646, found 593.2641.

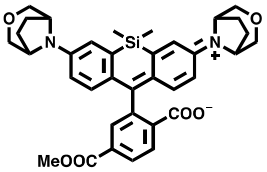


**BD 666$_{COOMe}$**: Compound **S9** (130 mg, 0.30 mmol, 1.0 eq.), 2-formyl-4-(methoxycarbonyl) benzoic acid (312 mg, 1.5 mmol, 5.0 eq.) and $CuBr_2$ (7 mg, 0.03 mmol, 0.1 eq.) were mixed in a sealable pressure tube. The tube was sealed tightly and heated at 140 °C for 4 h. After cooling to room temperature, the reaction mixture was dissolved in MeOH (5 mL), filtered, and concentrated *in vacuo*. The residue was purified by silica gel column chromatography (PE / EA = 10/1, v/v) to afford product **BD 666$_{COOMe}$** as a blue-green solid (97 mg, 52%).
**$^1$H NMR** (400 MHz, $CDCl_3$) δ 8.21 (dd, $J$ = 8.0, 1.3 Hz, 1H), 8.02 (d, $J$ = 8.0 Hz, 1H), 7.99 – 7.96 (m, 1H), 7.04 (d, $J$ = 2.7 Hz, 2H), 6.79 (d, $J$ = 8.8 Hz, 2H), 6.61 (dd, $J$ = 8.9, 2.8 Hz, 2H), 4.10 – 4.05 (m, 4H), 3.91 (s, 3H), 3.90 – 3.84 (m, 4H), 3.56 – 3.49 (m, 4H), 2.14 – 1.93 (m, 8H), 0.65 (s, 3H), 0.59 (s, 3H).
**$^{13}$C NMR** (101 MHz, $CDCl_3$) δ 169.9, 166.0, 154.6, 146.5, 137.4, 135.2, 133.0, 130.2, 130.1, 128.4, 126.0, 125.9, 120.0, 116.4, 91.7, 69.8, 69.7, 57.0, 52.8, 26.9, 26.9, 0.4, -1.2.
**HRMS** (ESI) calcd for $C_{36}H_{39}N_2O_6Si$ $[M+H]^+$ 623.2572, found 623.2560.

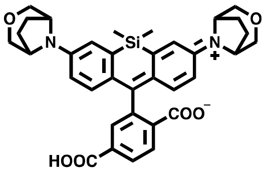


**BD 666$_{COOH}$**:To a solution of **BD666$_{COOMe}$** (97 mg, 0.16 mmol, 1.0 eq.) in THF/dd$H_2O$ (2:1, 2 mL in total) was added 1 N LiOH (800 μL, 0.80 mmol, 5.0 eq). The reaction was stirred at room temperature for 1 h. It was subsequently acidified with 1 N HCl (900 μL), diluted with water, and extracted with $CH_2Cl_2$ (3×). The organic extracts were combined and dried over $Na_2SO_4$, filtered, and concentrated *in vacuo*. The residue was purified by silica gel column chromatography ($CH_2Cl_2$ / MeOH = 25/1, v/v) to afford product **BD 666$_{COOH}$** as a blue-green solid (78 mg, 82%).
**$^1$H NMR** (400 MHz, DMSO-$d_6$) δ 8.17 (dd, $J$ = 7.9, 1.3 Hz, 1H), 8.07 (d, $J$ = 8.0 Hz, 1H), 7.79 – 7.75 (m, 1H), 7.21 (d, $J$ = 2.7 Hz, 2H), 6.77 (dd, $J$ = 8.9, 2.6 Hz, 2H), 6.68 (d, $J$ = 8.8 Hz, 2H), 4.37 – 4.08 (m, 4H), 3.69 – 3.61 (m, 4H), 3.47 – 3.39 (m, 4H), 1.98 – 1.80 (m, 8H), 0.64 (s, 3H), 0.54 (s, 3H).
**$^{13}$C NMR** (101 MHz, DMSO-$d_6$) δ 169.0, 166.1, 158.4, 158.1, 146.4, 136.4, 131.5, 130.2, 128.6, 127.6, 126.1, 124.7, 119.9, 116.4, 91.1, 68.9, 55.8, 26.4, 26.4, -0.1, -1.3.
**HRMS** (ESI) calcd for $C_{35}H_{37}N_2O_6Si$ $[M+H]^+$ 609.2415, found 609.2420.

**General Method C: Functionalization with HaloTag ligand**

Representative Procedure for **BD 566HTL**:

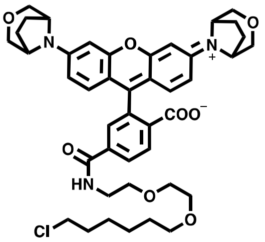

**BD 566COOH** (22 mg, 40 µmol, 1.0 eq.), HaloTag(O2)amine (13 mg, 60 µmol, 1.5 eq.) and BOP (21 mg, 48 µmol, 1.2 eq.) were dissolved in anhydrous DMF (2 mL). Then DIPEA (13 µL, 80 µmol, 2.0 eq.) was added and the mixture was stirred at room temperature overnight. Purification of the mixture by reverse phase HPLC (eluent, a 30-min linear gradient, from 30% to 95% solvent B; flow rate, 5.0 mL/min; detection wavelength, 560 nm; eluent A (dd$H_2O$ containing 0.1% TFA (v/v)) and eluent B ($CH_3CN$)) provided **BD 566HTL** (30 mg, 84% yield) as a purple solid.

**$^1$H NMR** (400 MHz, $CD_3OD$) δ 8.42 (d, *J* = 8.3 Hz, 1H), 8.22 (dd, *J* = 8.2, 1.8 Hz, 1H), 7.84 (d, *J* = 1.8 Hz, 1H), 7.19 – 7.05 (m, 6H), 4.68 – 4.59 (m, 4H), 3.80 – 3.70 (m, 8H), 3.70 – 3.53 (m, 8H), 3.52 (t, *J* = 6.6 Hz, 2H), 3.44 (t, *J* = 6.5 Hz, 2H), 2.28 – 2.05 (m, 8H), 1.76 – 1.67 (m, 2H), 1.55 – 1.46 (m, 2H), 1.44 – 1.27 (m, 4H).

**Analytical HPLC:** 98.6% purity (4.6 mm × 150 mm 5 µm C18 column; 2 µL injection; 5-100% $CH_3CN/H_2O$, linear-gradient, with constant 0.1% v/v TFA additive; 6 min run; 0.6 mL/min flow; ESI; positive ion mode; detection at 560 nm).

**HRMS** (ESI) calcd for $C_{43}H_{51}ClN_3O_8$ $[M+H]^+$ 772.3359, found 772.3383.

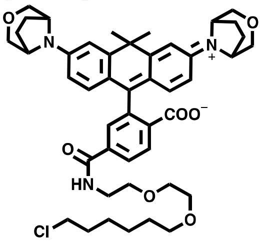

**BD 626HTL**: The title compound (28 mg, 87%, blue solid) was prepared from **BD 626COOH** according to general method C.

**$^1$H NMR** (400 MHz, $CD_3CN$) δ 8.27 (d, *J* = 8.2 Hz, 1H), 8.06 (dd, *J* = 8.2, 1.8 Hz, 1H), 7.66 (d, *J* = 1.7 Hz, 1H), 7.31 (t, *J* = 5.4 Hz, 1H), 7.18 (d, *J* = 2.4 Hz, 2H), 6.93 (d, *J* = 9.3 Hz, 2H), 6.71 (dd, *J* = 9.3, 2.4 Hz, 2H), 4.61 – 4.54 (m, 4H), 3.76 – 3.63 (m, 8H), 3.63 – 3.46 (m, 10H), 3.35 (t, *J* = 6.5 Hz, 2H), 2.17 – 2.01 (m, 8H), 1.82 (s, 3H), 1.72 (s, 3H), 1.70 – 1.65 (m, 2H), 1.51 – 1.20 (m, 6H).

**Analytical HPLC:** >99% purity (4.6 mm × 150 mm 5 µm C18 column; 2 µL injection; 5-100% $CH_3CN/H_2O$, linear-gradient, with constant 0.1% v/v TFA additive; 6 min run; 0.6 mL/min flow; ESI; positive ion mode; detection at 620 nm).

**HRMS** (ESI) calcd for $C_{46}H_{57}ClN_3O_7$ $[M+H]^+$ 798.3880, found 798.3872.

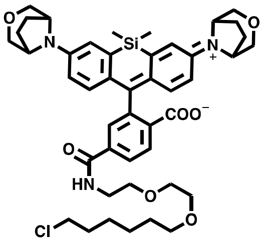


**BD 666HTL**: The title compound (25 mg, 78%, blue-green solid) was prepared from **BD 666COOH** according to general method C.

$^{1}$H NMR (400 MHz, $CD_3OD$) δ 8.29 (d, *J* = 8.2 Hz, 1H), 8.13 (dd, *J* = 8.2, 1.8 Hz, 1H), 7.74 (d, *J* = 1.7 Hz, 1H), 7.39 (d, *J* = 2.7 Hz, 2H), 6.95 (d, *J* = 9.4 Hz, 2H), 6.78 (dd, *J* = 9.5, 2.7 Hz, 2H), 4.75 – 4.54 (m, 4H), 3.79 – 3.55 (m, 16H), 3.51 (t, *J* = 6.6 Hz, 2H), 3.43 (t, *J* = 6.5 Hz, 2H), 2.26 – 2.01 (m, 8H), 1.76 – 1.65 (m, 2H), 1.51 (p, *J* = 6.8 Hz, 2H), 1.46 – 1.24 (m, 2H), 0.64 (s, 3H), 0.59 (s, 3H).

**Analytical HPLC:** 98.8% purity (4.6 mm × 150 mm 5 µm C18 column; 2 µL injection; 5-100% $CH_3CN/H_2O$, linear-gradient, with constant 0.1% v/v TFA additive; 6 min run; 0.6 mL/min flow; ESI; positive ion mode; detection at 660 nm).

**HRMS** (ESI) calcd for $C_{45}H_{57}ClN_3O_7Si$ $[M+H]^+$ 814.3649 found 814.3650.

### NMR spectra

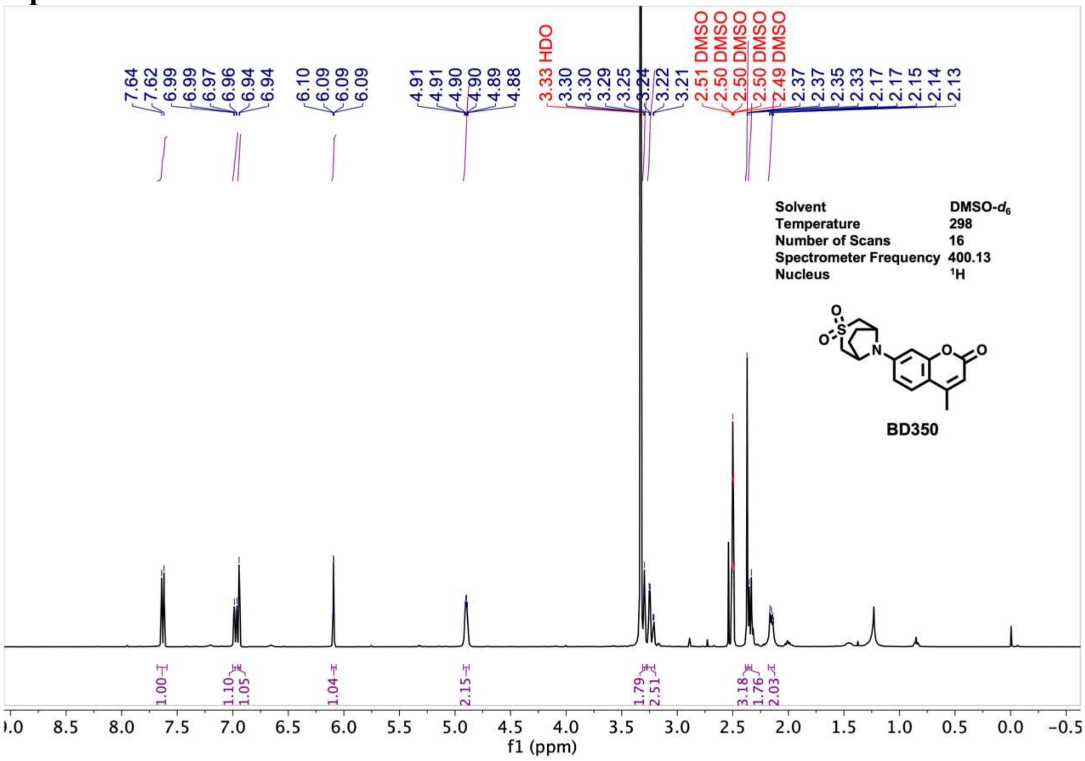

7.64
7.62
6.99
6.99
6.97
6.96
6.94
6.94
6.10
6.09
6.09
6.09
4.91
4.91
4.90
4.90
4.89
4.88
3.33 HDO
3.30
3.30
3.29
3.25
3.24
3.22
3.21
2.51 DMSO
2.50 DMSO
2.50 DMSO
2.50 DMSO
2.49 DMSO
2.37
2.37
2.35
2.33
2.17
2.17
2.15
2.14
2.13
Solvent DMSO-$d_6$
Temperature 298
Number of Scans 16
Spectrometer Frequency 400.13
Nucleus $^1$H
BD350
1.00
1.10
1.05
1.04
2.15
1.79
2.51
3.18
1.76
2.03
f1 (ppm)


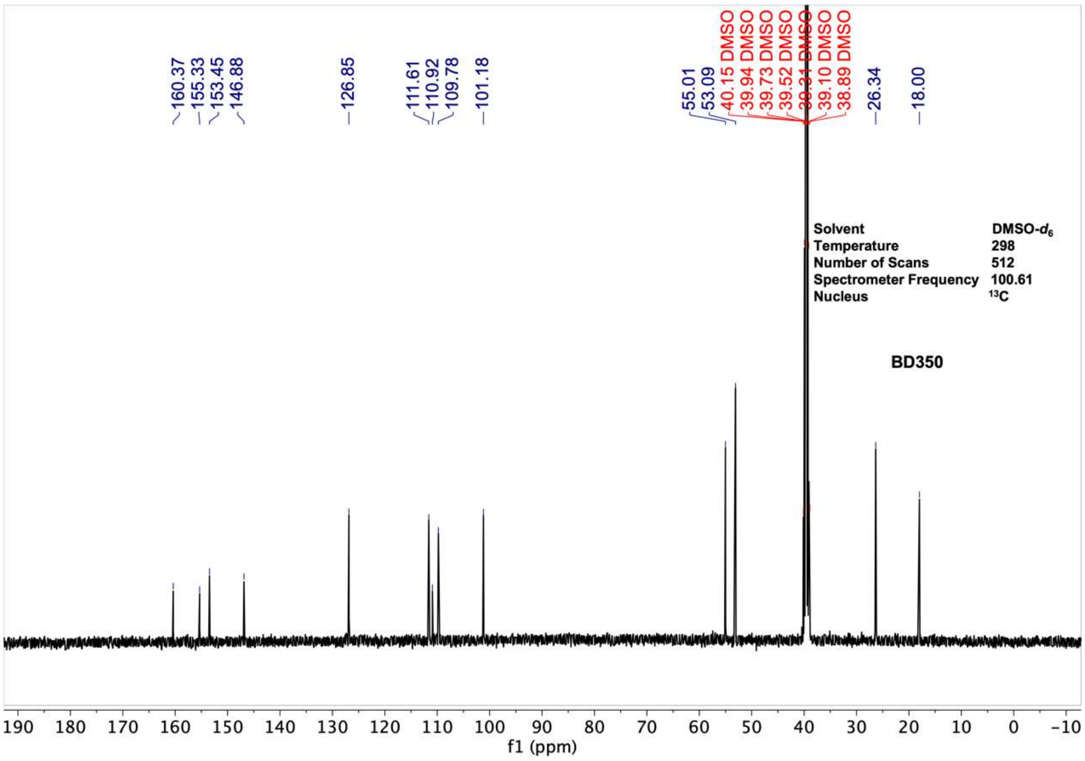

160.37
155.33
153.45
146.88
126.85
111.61
110.92
109.78
101.18
55.01
53.09
40.15 DMSO
39.94 DMSO
39.73 DMSO
39.52 DMSO
39.31 DMSO
39.10 DMSO
38.89 DMSO
26.34
18.00
Solvent DMSO-$d_6$
Temperature 298
Number of Scans 512
Spectrometer Frequency 100.61
Nucleus $^{13}$C
BD350
f1 (ppm)

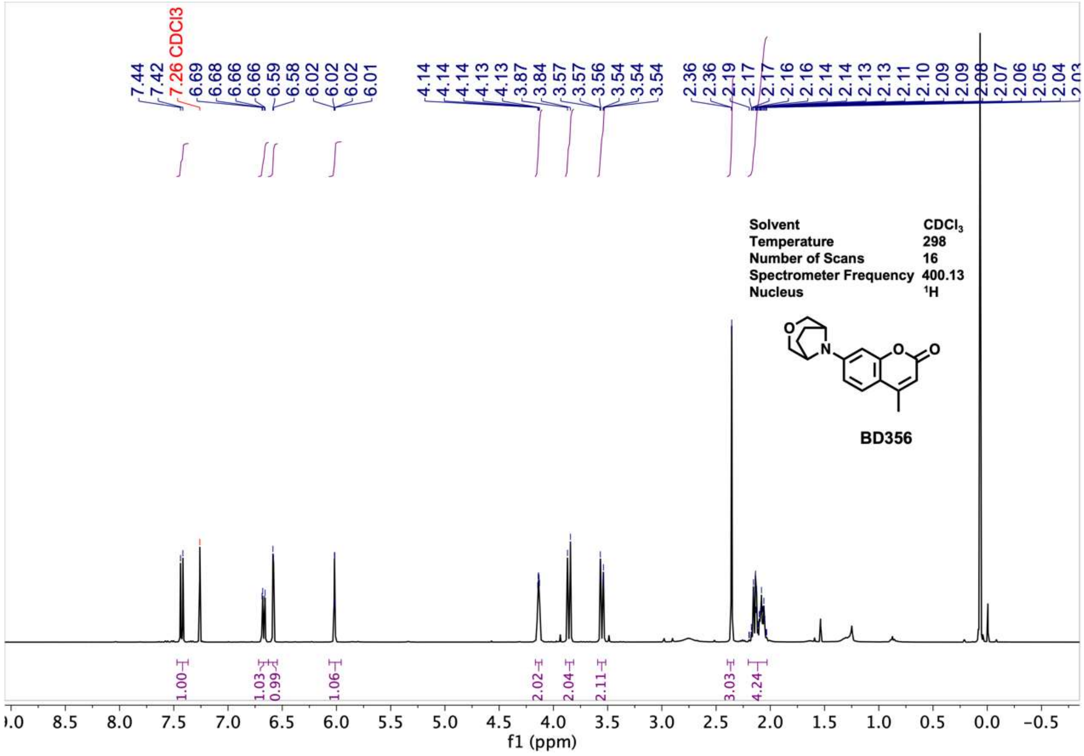
Solvent CDCl3
Temperature 298
Number of Scans 16
Spectrometer Frequency 400.13
Nucleus 1H
BD356
f1 (ppm)

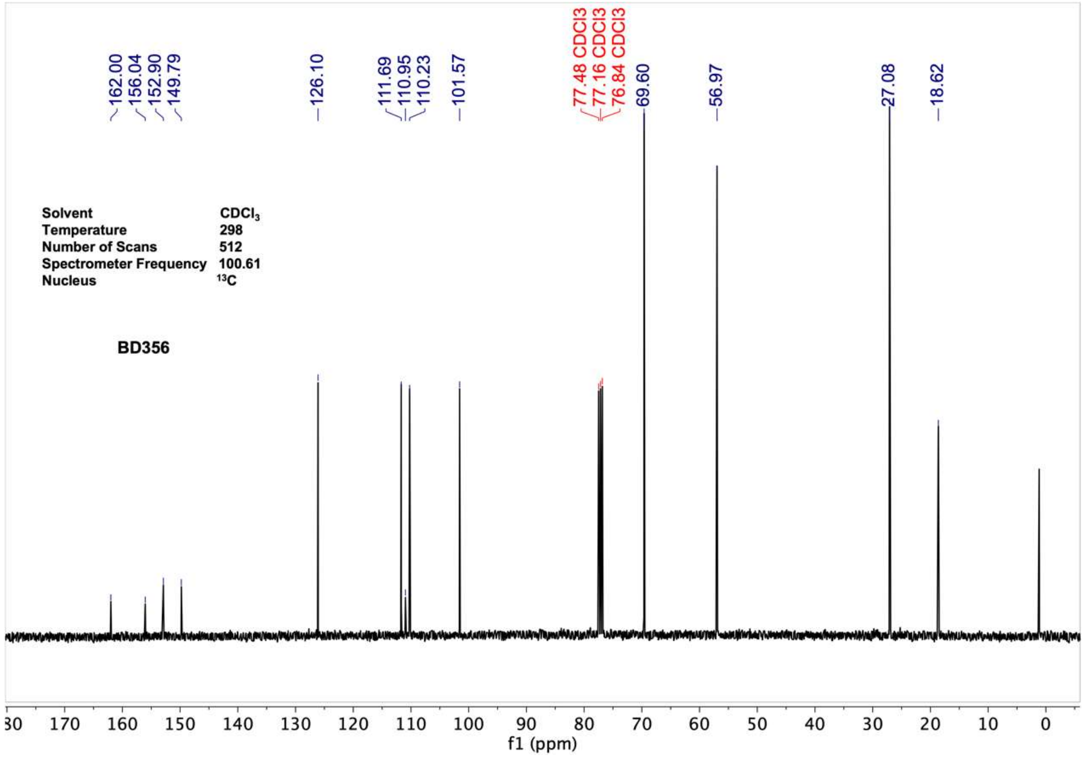
Solvent CDCl3
Temperature 298
Number of Scans 512
Spectrometer Frequency 100.61
Nucleus 13C
BD356
f1 (ppm)

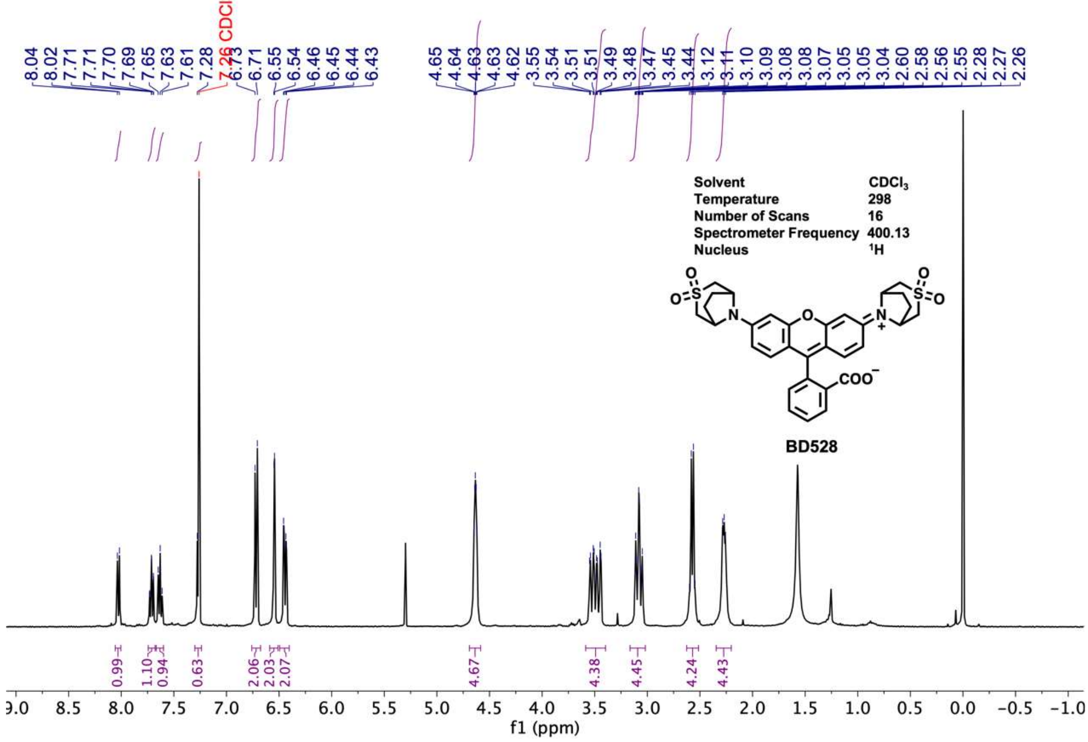
8.04
8.02
7.71
7.71
7.70
7.69
7.65
7.63
7.61
7.28
7.26 CDCl3
6.73
6.71
6.55
6.54
6.46
6.45
6.44
6.43
4.65
4.64
4.63
4.63
4.62
3.55
3.54
3.51
3.51
3.49
3.48
3.47
3.45
3.44
3.12
3.11
3.10
3.09
3.08
3.08
3.07
3.05
3.05
3.04
2.60
2.58
2.56
2.55
2.28
2.27
2.26
Solvent CDCl3
Temperature 298
Number of Scans 16
Spectrometer Frequency 400.13
Nucleus 1H
BD528
0.99
1.10
0.94
0.63
2.06
2.03
2.07
4.67
4.38
4.45
4.24
4.43
9.0 8.5 8.0 7.5 7.0 6.5 6.0 5.5 5.0 4.5 4.0 3.5 3.0 2.5 2.0 1.5 1.0 0.5 0.0 −0.5 −1.0
f1 (ppm)

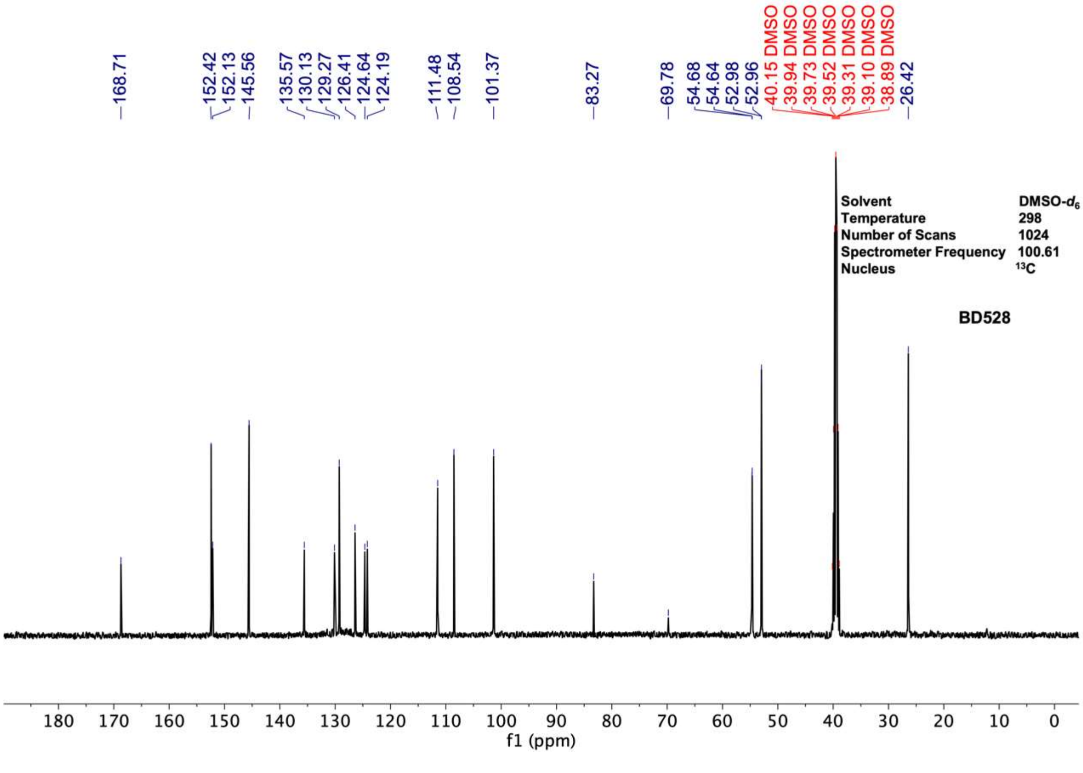
168.71
152.42
152.13
145.56
135.57
130.13
129.27
126.41
124.64
124.19
111.48
108.54
101.37
83.27
69.78
54.68
54.64
52.98
52.96
40.15 DMSO
39.94 DMSO
39.73 DMSO
39.52 DMSO
39.31 DMSO
39.10 DMSO
38.89 DMSO
26.42
Solvent DMSO-d6
Temperature 298
Number of Scans 1024
Spectrometer Frequency 100.61
Nucleus 13C
BD528
180 170 160 150 140 130 120 110 100 90 80 70 60 50 40 30 20 10 0
f1 (ppm)

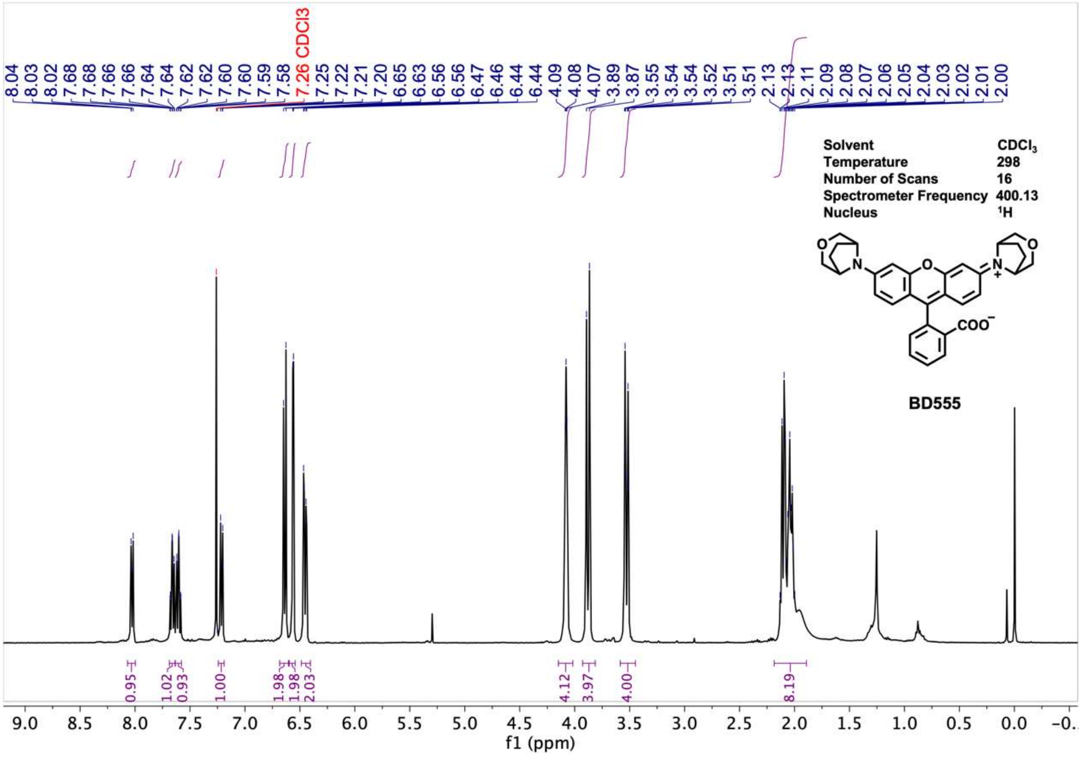
8.04
8.03
8.02
7.68
7.68
7.66
7.64
7.64
7.62
7.62
7.60
7.60
7.59
7.58
7.26 CDCl3
7.25
7.22
7.21
7.20
6.65
6.63
6.56
6.56
6.47
6.46
6.44
6.44
4.09
4.08
4.07
3.89
3.87
3.55
3.54
3.54
3.52
3.51
3.51
2.13
2.13
2.11
2.09
2.08
2.07
2.06
2.05
2.04
2.03
2.02
2.01
2.00
Solvent CDCl3
Temperature 298
Number of Scans 16
Spectrometer Frequency 400.13
Nucleus 1H
BD555
0.95
1.02
0.93
1.00
1.98
1.98
2.03
4.12
3.97
4.00
8.19
f1 (ppm)

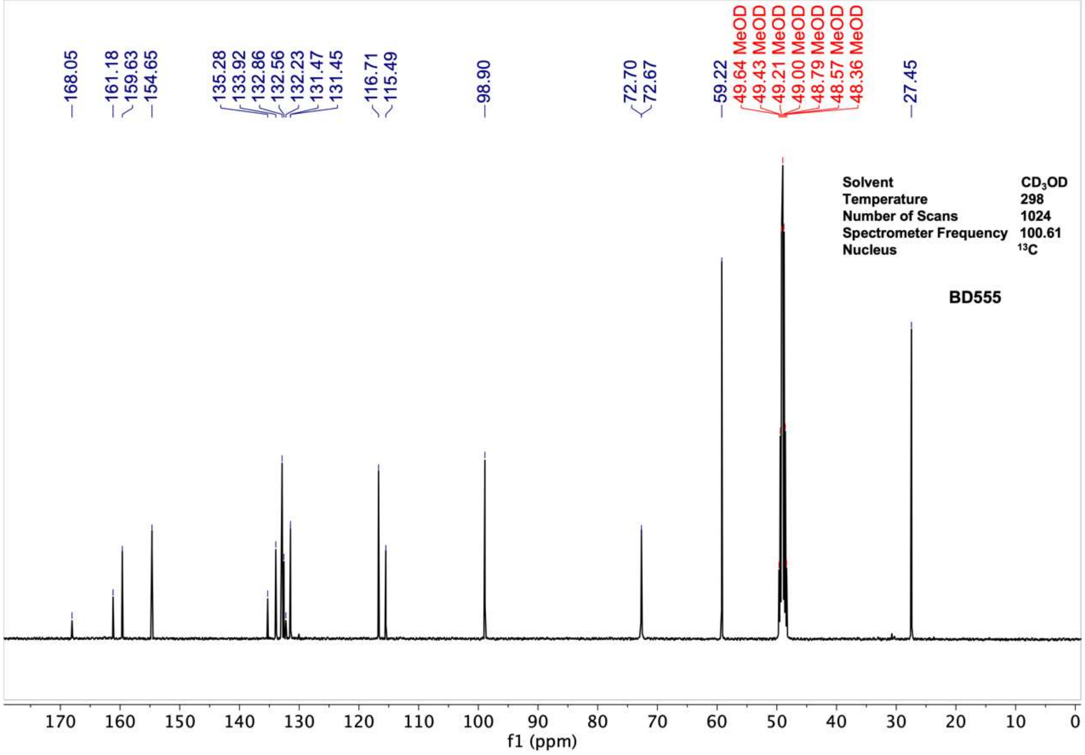
168.05
161.18
159.63
154.65
135.28
133.92
132.86
132.56
132.23
131.47
131.45
116.71
115.49
98.90
72.70
72.67
59.22
49.64 MeOD
49.43 MeOD
49.21 MeOD
49.00 MeOD
48.79 MeOD
48.57 MeOD
48.36 MeOD
27.45
Solvent CD3OD
Temperature 298
Number of Scans 1024
Spectrometer Frequency 100.61
Nucleus 13C
BD555
f1 (ppm)

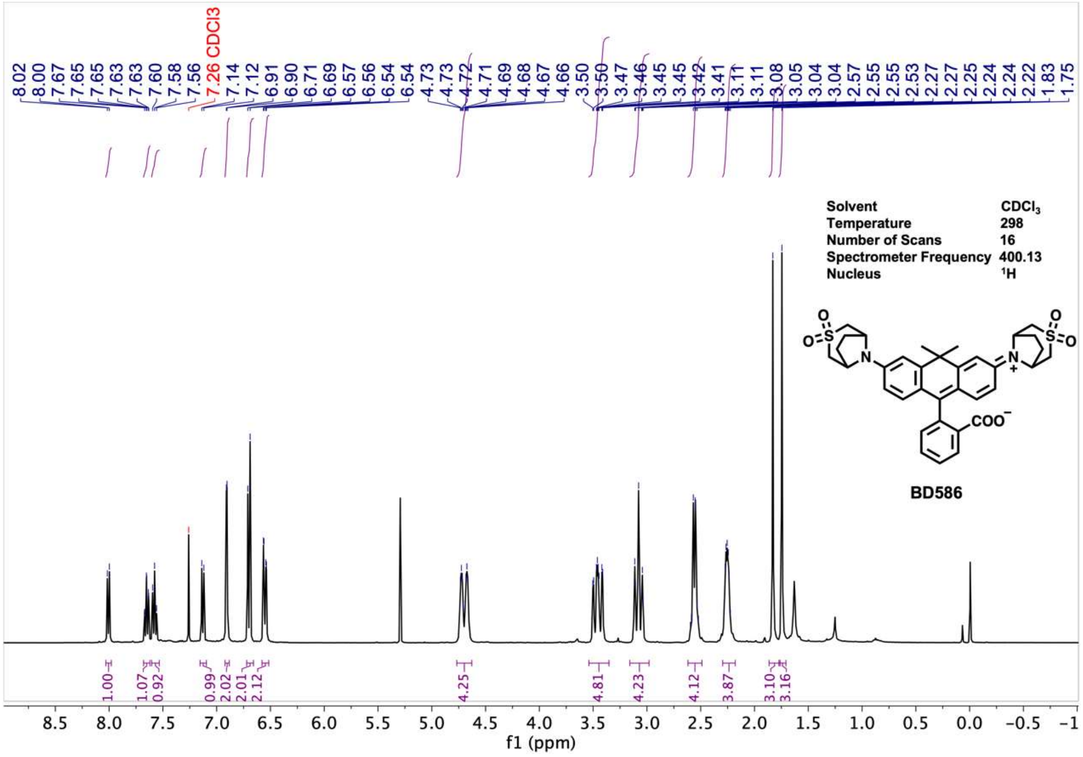

Solvent CDCl3
Temperature 298
Number of Scans 16
Spectrometer Frequency 400.13
Nucleus 1H
BD586
f1 (ppm)


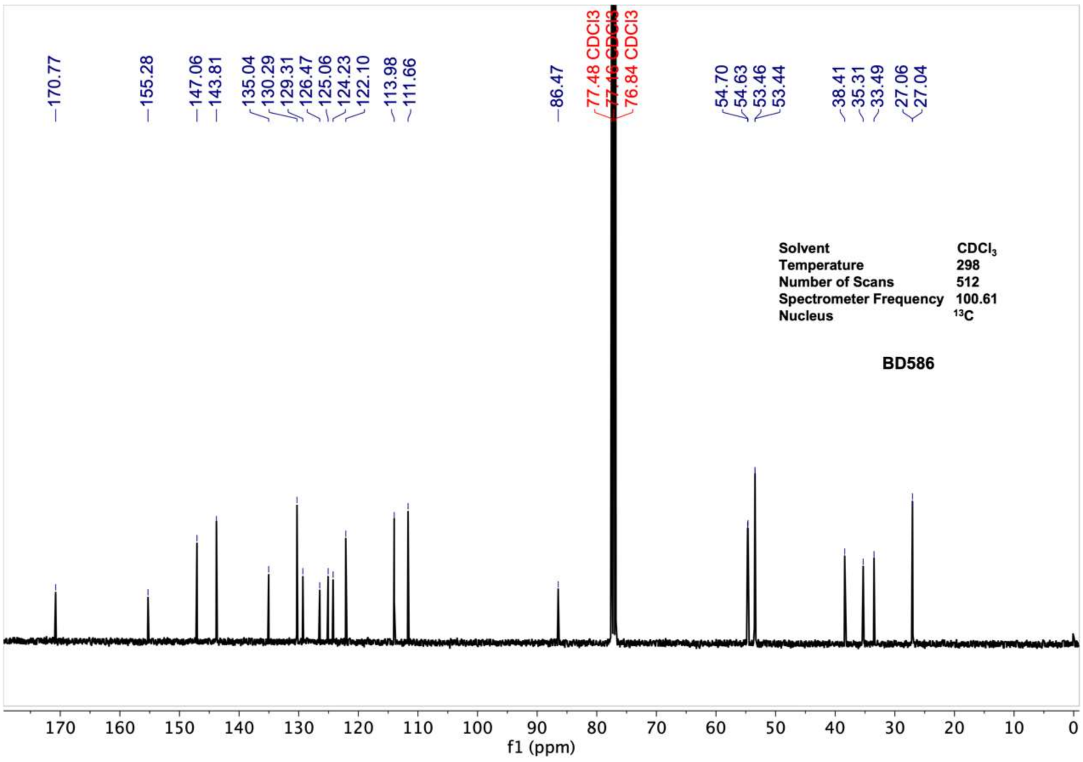

Solvent CDCl3
Temperature 298
Number of Scans 512
Spectrometer Frequency 100.61
Nucleus 13C
BD586
f1 (ppm)

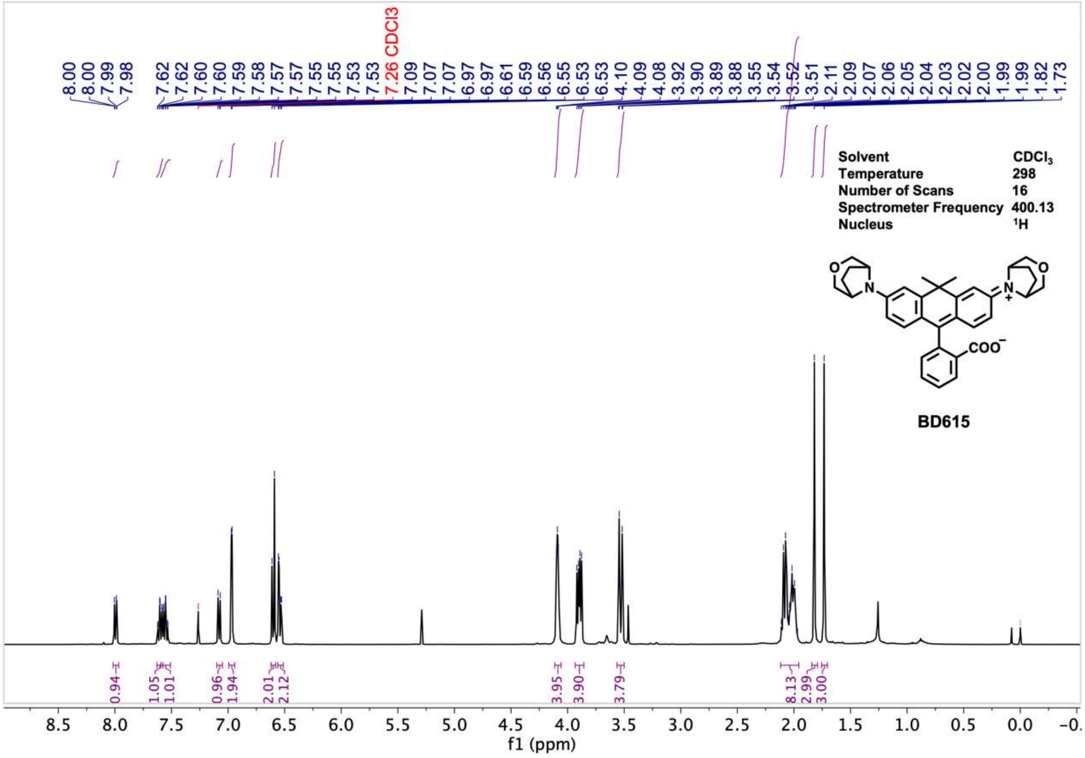
8.00
8.00
7.99
7.98
7.62
7.62
7.60
7.60
7.59
7.58
7.57
7.57
7.55
7.55
7.53
7.53
7.26 CDCl3
7.09
7.07
7.07
6.97
6.97
6.61
6.59
6.56
6.55
6.53
6.53
4.10
4.09
4.08
3.92
3.90
3.89
3.88
3.55
3.54
3.52
3.51
2.11
2.09
2.07
2.06
2.05
2.04
2.03
2.02
2.00
1.99
1.99
1.82
1.73
Solvent CDCl$_3$
Temperature 298
Number of Scans 16
Spectrometer Frequency 400.13
Nucleus $^1$H
COO$^-$
N$^+$
BD615
0.94
1.05
1.01
0.96
1.94
2.01
2.12
3.95
3.90
3.79
8.13
2.99
3.00
f1 (ppm)

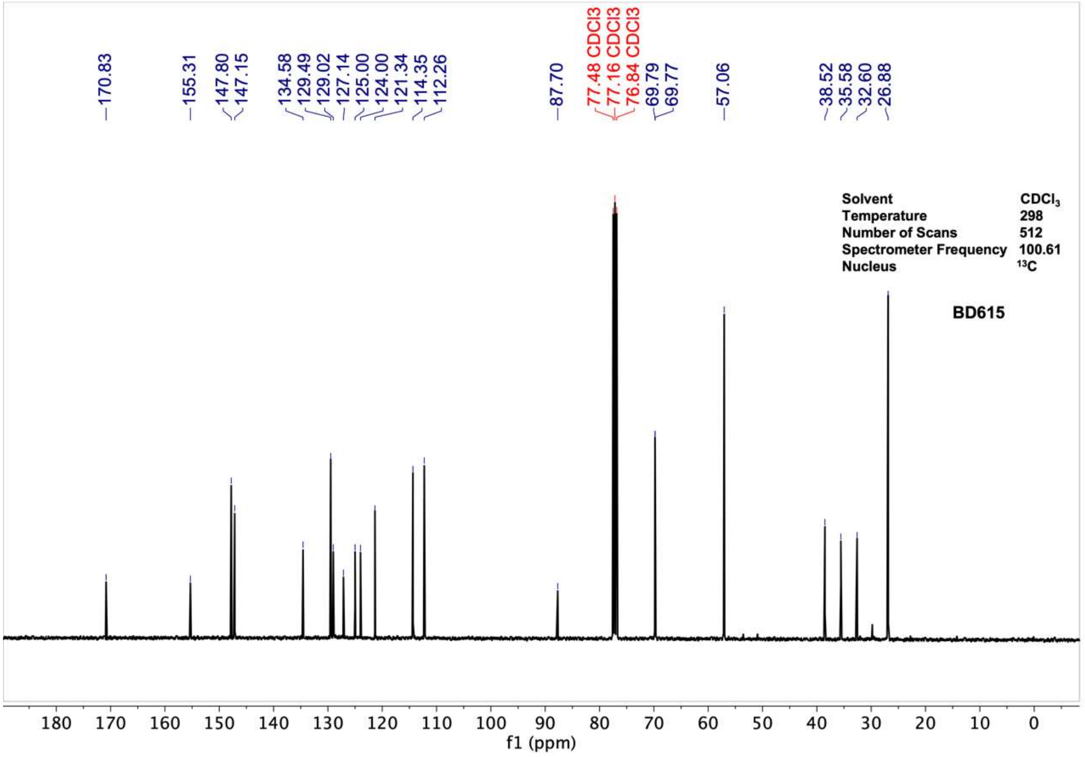
170.83
155.31
147.80
147.15
134.58
129.49
129.02
127.14
125.00
124.00
121.34
114.35
112.26
87.70
77.48 CDCl3
77.16 CDCl3
76.84 CDCl3
69.79
69.77
57.06
38.52
35.58
32.60
26.88
Solvent CDCl$_3$
Temperature 298
Number of Scans 512
Spectrometer Frequency 100.61
Nucleus $^{13}$C
BD615
f1 (ppm)

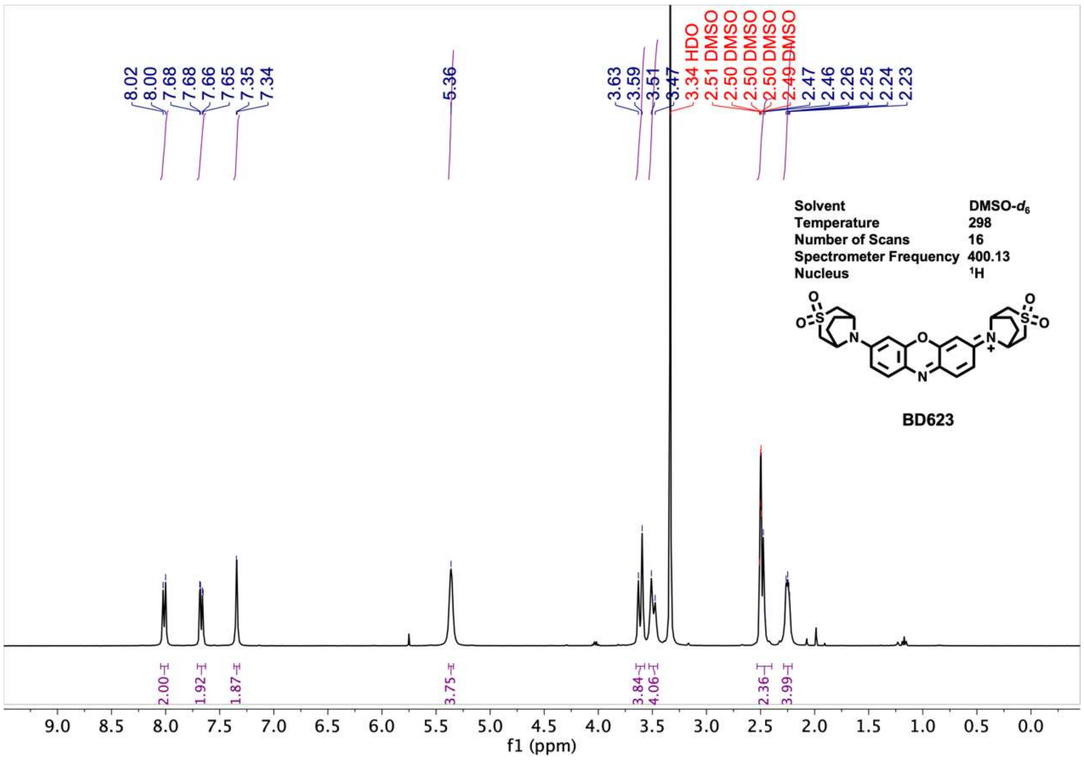
8.02
8.00
7.68
7.68
7.66
7.65
7.35
7.34
5.36
3.63
3.59
3.51
3.47
3.34 HDO
2.51 DMSO
2.50 DMSO
2.50 DMSO
2.50 DMSO
2.49 DMSO
2.47
2.46
2.26
2.25
2.24
2.23
Solvent DMSO-$d_6$
Temperature 298
Number of Scans 16
Spectrometer Frequency 400.13
Nucleus $^1H$
BD623
2.00
1.92
1.87
3.75
3.84
4.06
2.36
3.99
f1 (ppm)

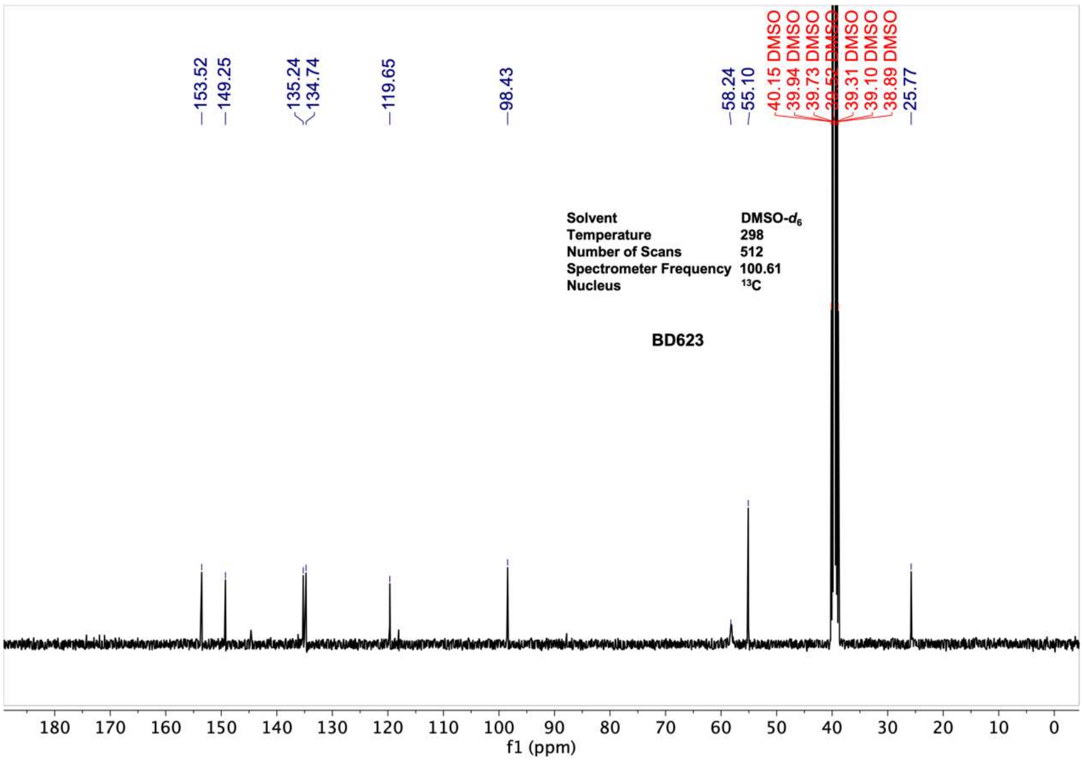
153.52
149.25
135.24
134.74
119.65
98.43
58.24
55.10
40.15 DMSO
39.94 DMSO
39.73 DMSO
39.52 DMSO
39.31 DMSO
39.10 DMSO
38.89 DMSO
25.77
Solvent DMSO-$d_6$
Temperature 298
Number of Scans 512
Spectrometer Frequency 100.61
Nucleus $^{13}C$
BD623
f1 (ppm)

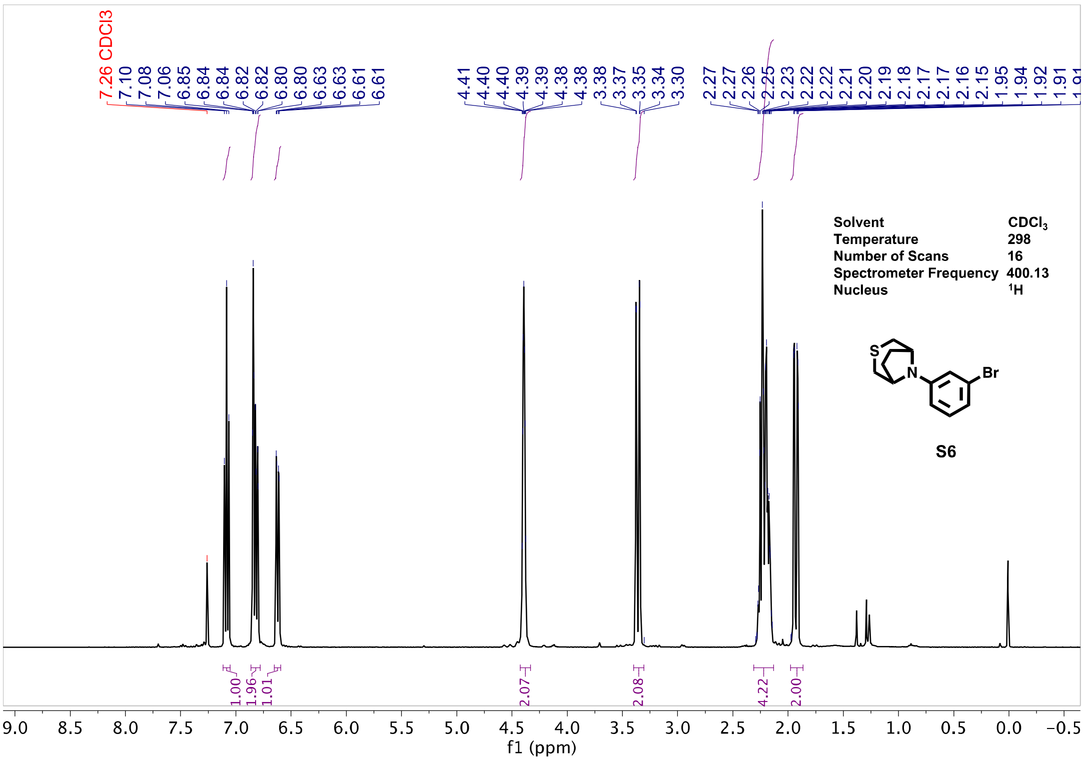
Solvent CDCl3
Temperature 298
Number of Scans 16
Spectrometer Frequency 400.13
Nucleus 1H
S6
f1 (ppm)

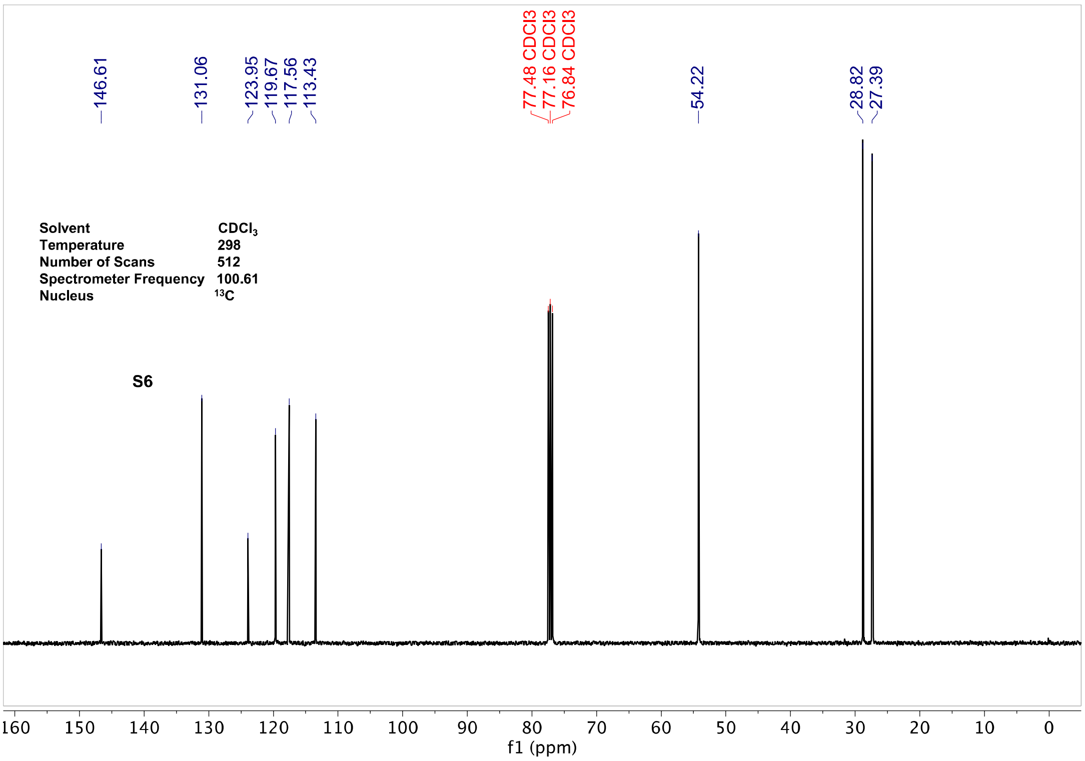
Solvent CDCl3
Temperature 298
Number of Scans 512
Spectrometer Frequency 100.61
Nucleus 13C
S6
146.61
131.06
123.95
119.67
117.56
113.43
77.48 CDCl3
77.16 CDCl3
76.84 CDCl3
54.22
28.82
27.39
f1 (ppm)

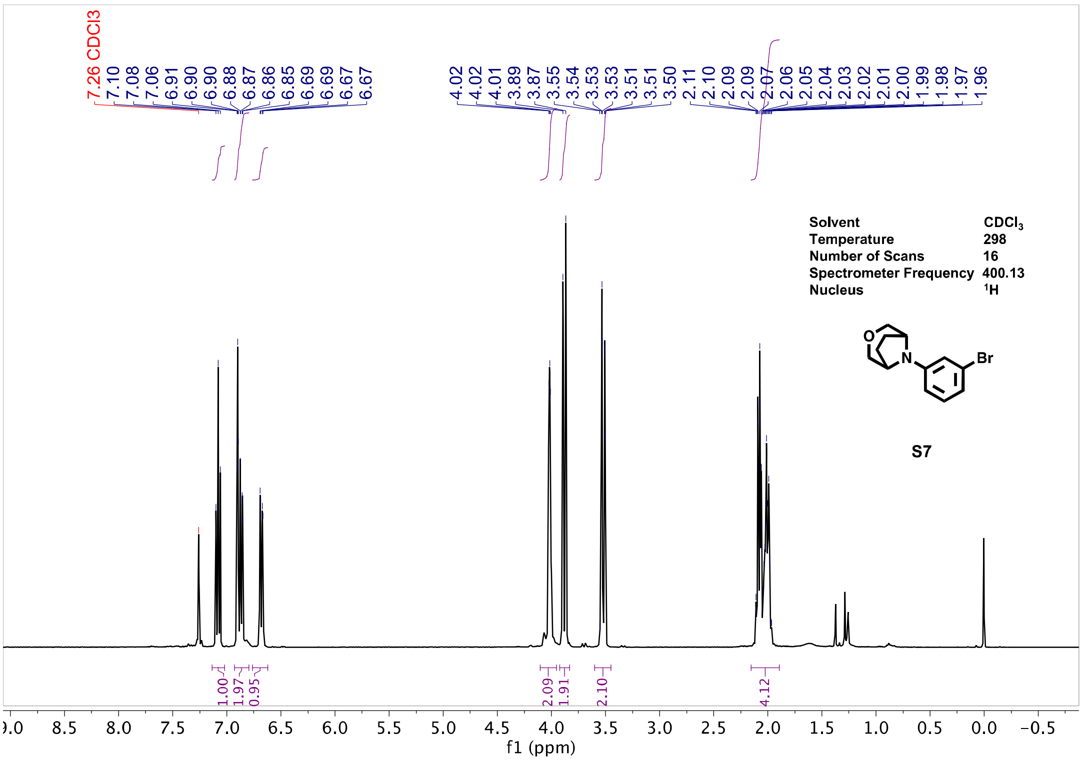
7.26 CDCl3
7.10
7.08
7.06
6.91
6.90
6.90
6.88
6.87
6.86
6.85
6.69
6.69
6.67
6.67
4.02
4.02
4.01
3.89
3.87
3.55
3.54
3.53
3.53
3.51
3.51
3.50
2.11
2.10
2.09
2.09
2.07
2.06
2.05
2.04
2.03
2.02
2.01
2.00
1.99
1.98
1.97
1.96
Solvent CDCl3
Temperature 298
Number of Scans 16
Spectrometer Frequency 400.13
Nucleus 1H
O
N
Br
S7
1.00
1.97
0.95
2.09
1.91
2.10
4.12
f1 (ppm)

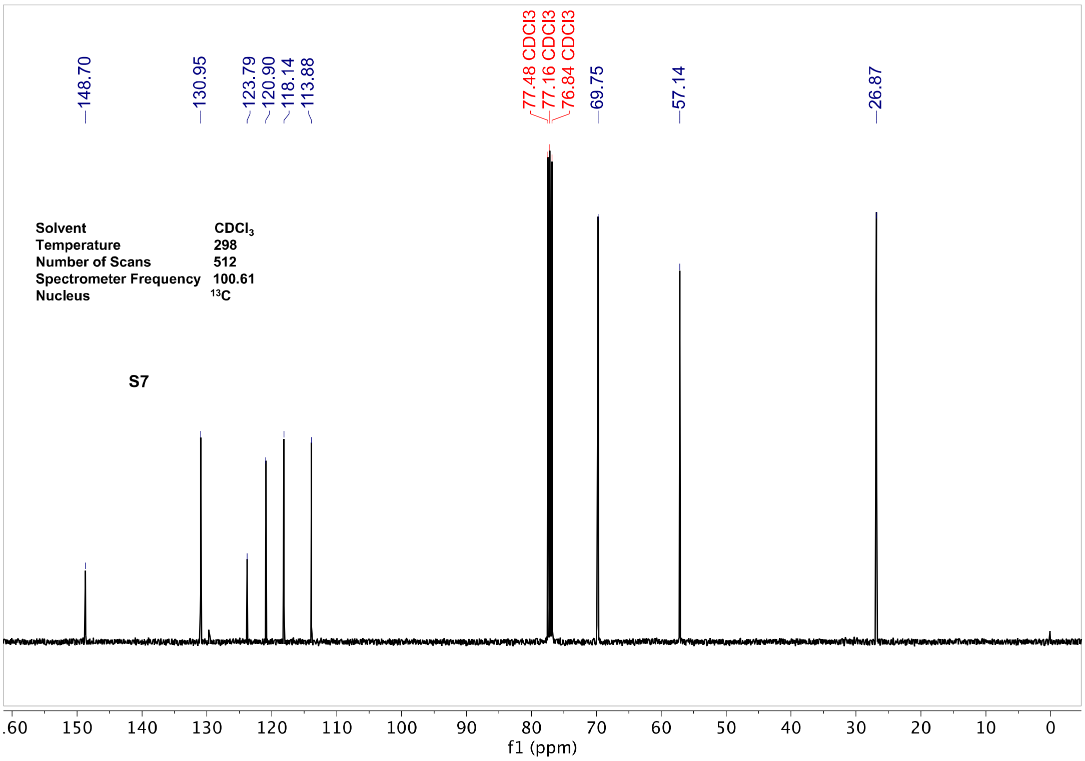
148.70
130.95
123.79
120.90
118.14
113.88
77.48 CDCl3
77.16 CDCl3
76.84 CDCl3
69.75
57.14
26.87
Solvent CDCl3
Temperature 298
Number of Scans 512
Spectrometer Frequency 100.61
Nucleus 13C
S7
f1 (ppm)

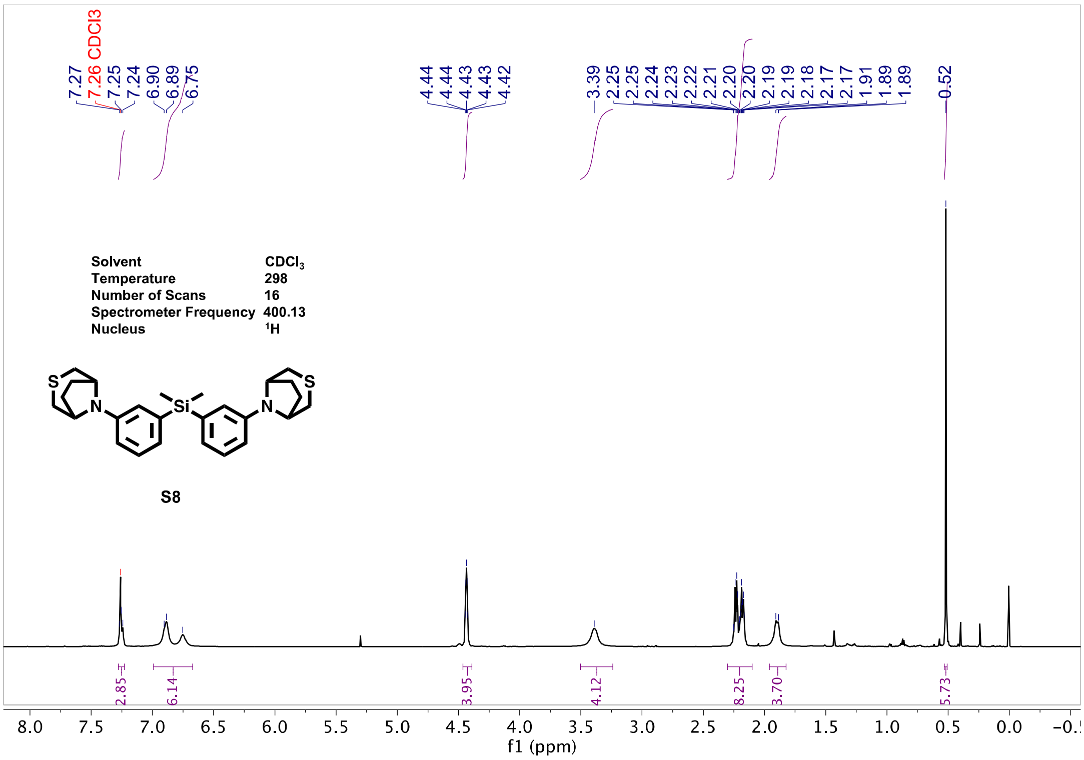
Solvent CDCl3
Temperature 298
Number of Scans 16
Spectrometer Frequency 400.13
Nucleus 1H
S8
f1 (ppm)

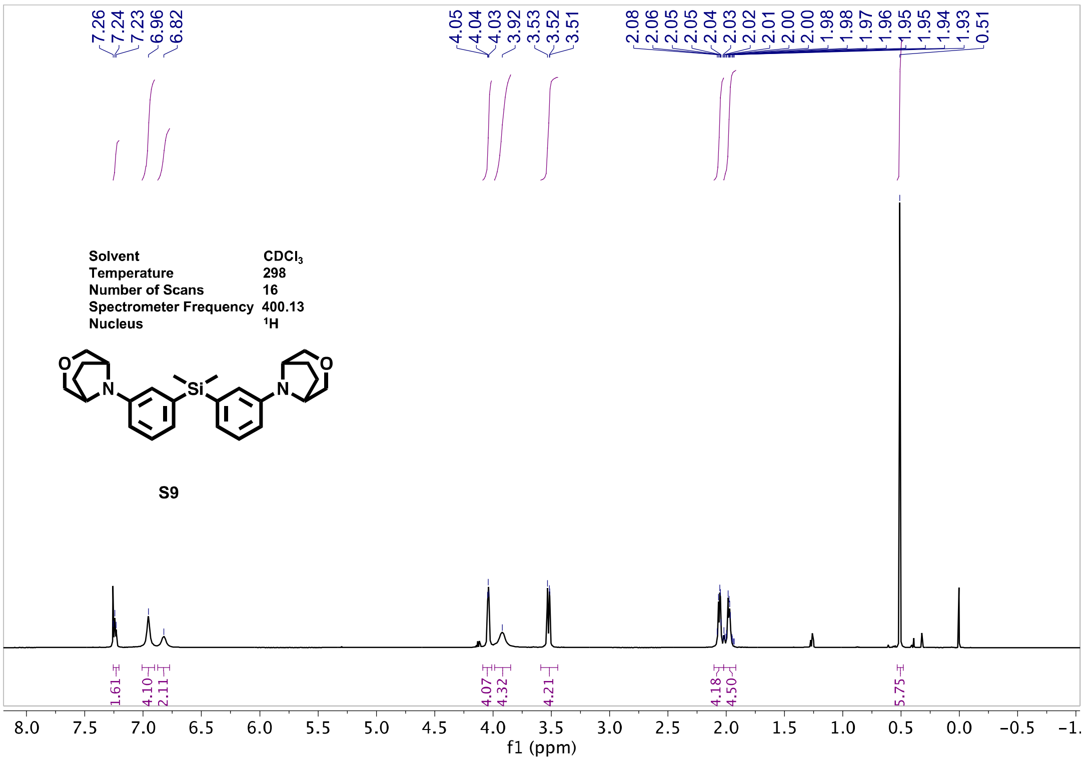
Solvent CDCl3
Temperature 298
Number of Scans 16
Spectrometer Frequency 400.13
Nucleus 1H
S9
f1 (ppm)

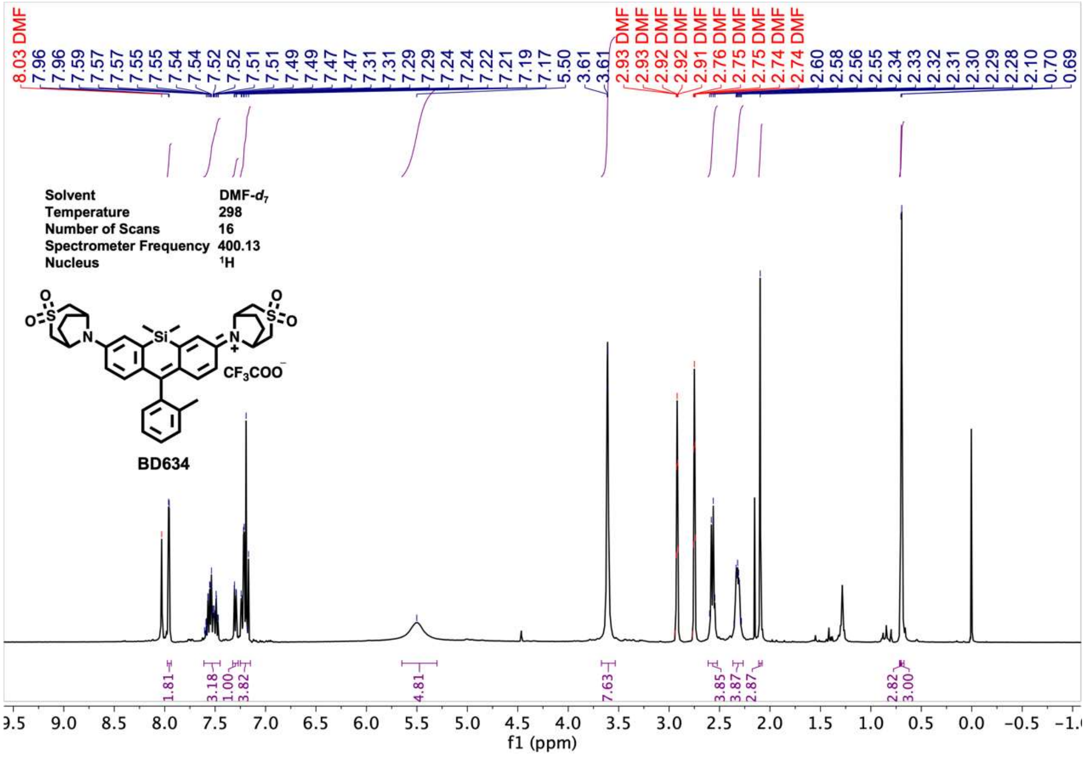
Solvent DMF-$d_7$
Temperature 298
Number of Scans 16
Spectrometer Frequency 400.13
Nucleus $^1H$
$CF_3COO^-$
BD634
f1 (ppm)

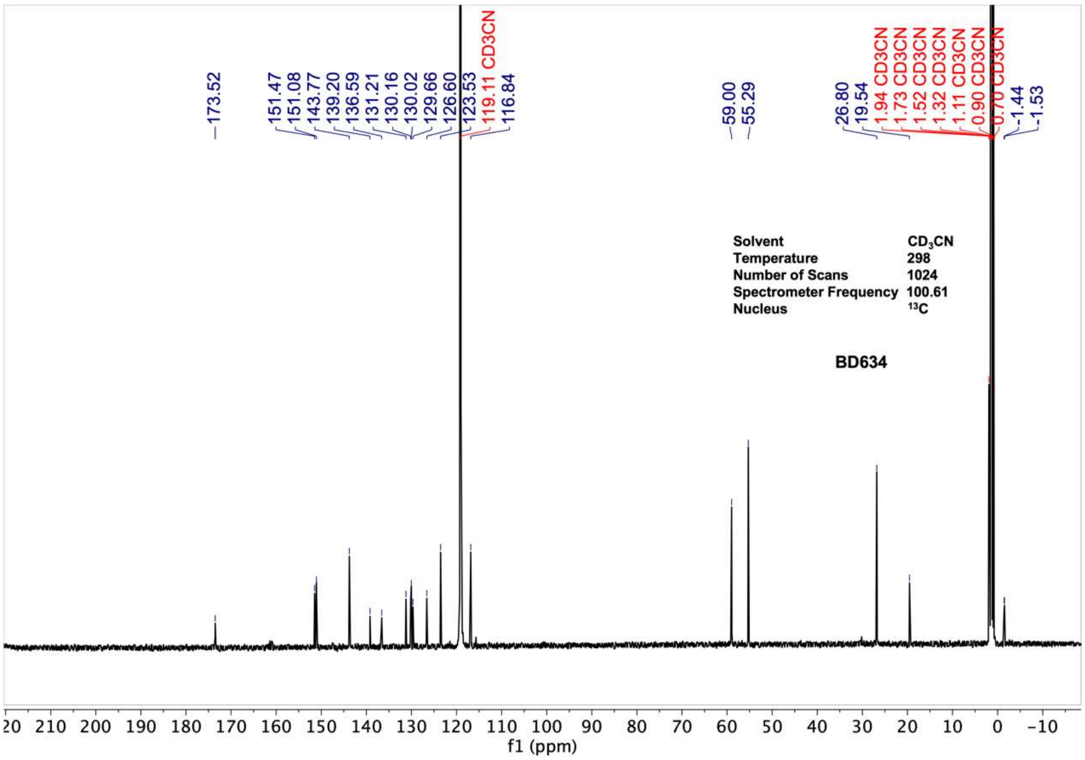
Solvent $CD_3CN$
Temperature 298
Number of Scans 1024
Spectrometer Frequency 100.61
Nucleus $^{13}C$
BD634
f1 (ppm)

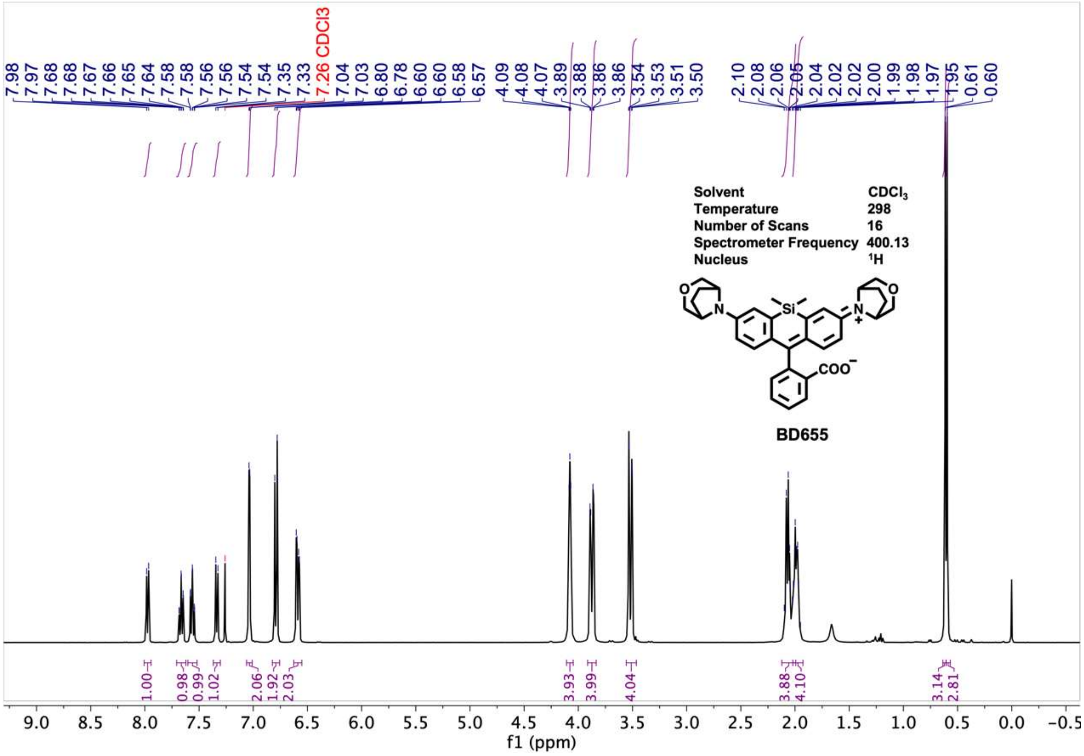
7.98 7.97 7.68 7.68 7.67 7.66 7.65 7.64 7.58 7.58 7.56 7.56 7.54 7.54 7.35 7.33
7.26 CDCl3
7.04 7.03 6.80 6.78 6.60 6.60 6.58 6.57 4.09 4.08 4.07 3.89 3.88 3.86 3.86 3.54 3.53 3.51 3.50 2.10 2.08 2.06 2.05 2.04 2.02 2.02 2.00 1.99 1.98 1.97 1.95 0.61 0.60
Solvent CDCl3
Temperature 298
Number of Scans 16
Spectrometer Frequency 400.13
Nucleus 1H
BD655
1.00 0.98 0.99 1.02 2.06 1.92 2.03 3.93 3.99 4.04 3.88 4.10 3.14 2.81
f1 (ppm)

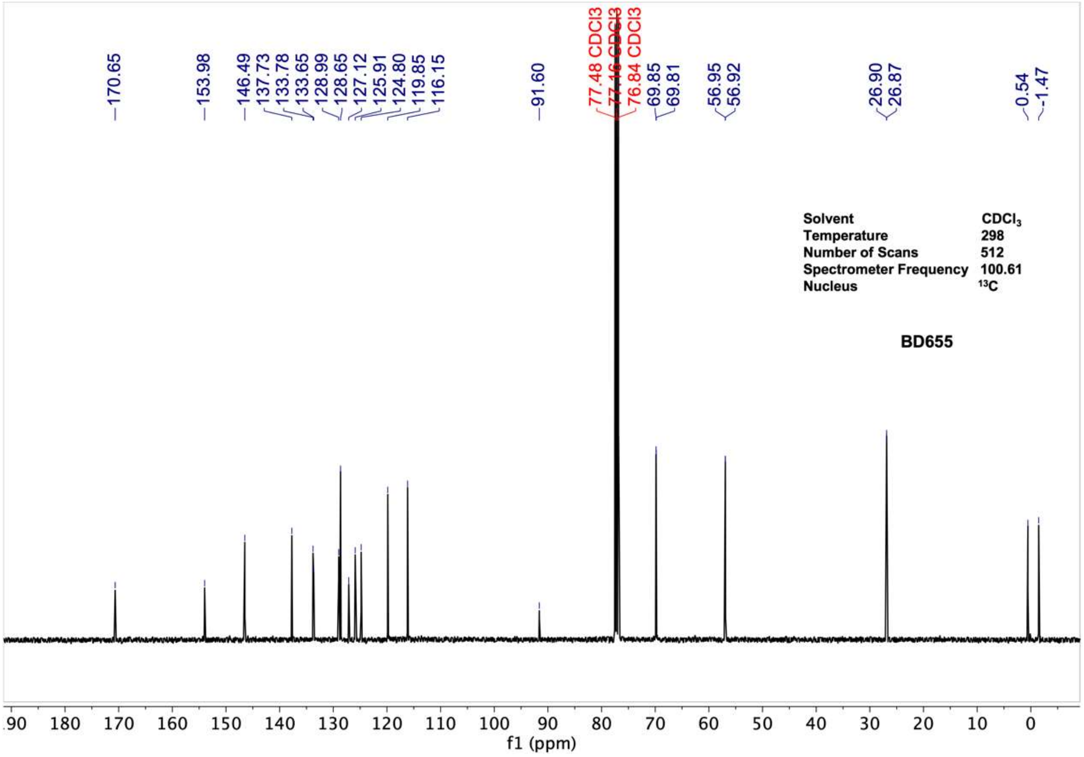
170.65 153.98 146.49 137.73 133.78 133.65 128.99 128.65 127.12 125.91 124.80 119.85 116.15 91.60
77.48 CDCl3
77.16 CDCl3
76.84 CDCl3
69.85 69.81 56.95 56.92 26.90 26.87 0.54 -1.47
Solvent CDCl3
Temperature 298
Number of Scans 512
Spectrometer Frequency 100.61
Nucleus 13C
BD655
f1 (ppm)

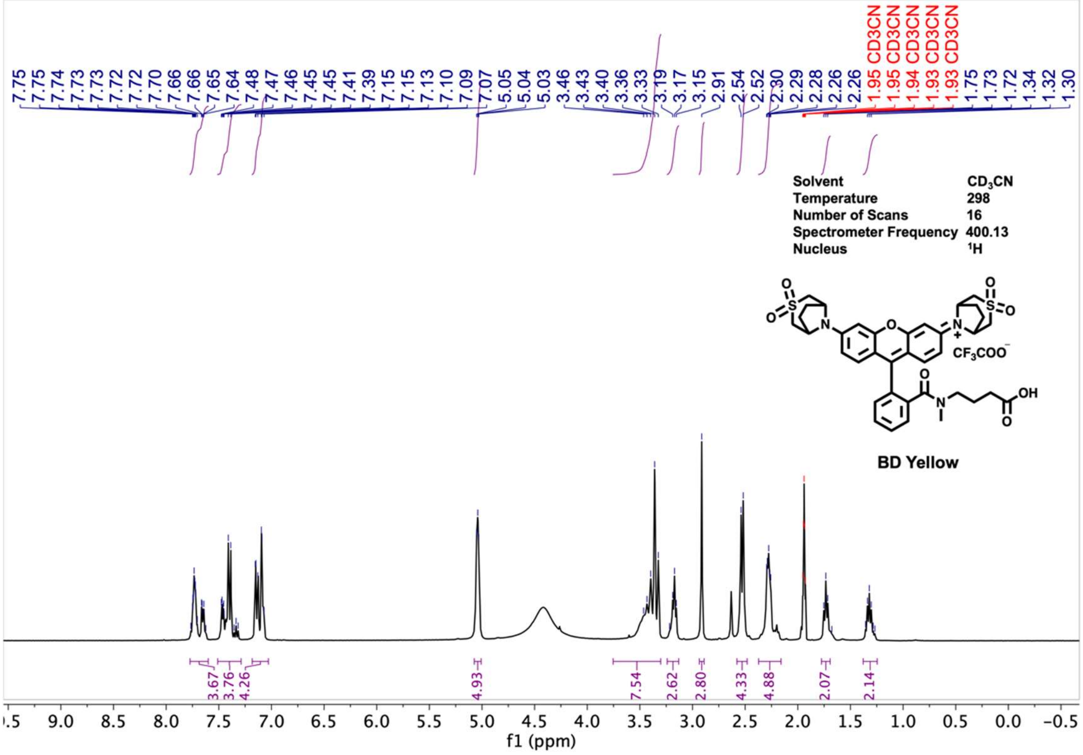
7.75 7.75 7.74 7.73 7.73 7.72 7.72 7.70 7.66 7.66 7.65 7.64 7.48 7.47 7.46 7.45 7.45 7.41 7.39 7.15 7.15 7.13 7.10 7.09 7.07 5.05 5.04 5.03 3.46 3.43 3.40 3.36 3.33 3.19 3.17 3.15 2.91 2.54 2.52 2.30 2.29 2.28 2.26 2.26
1.95 CD3CN 1.95 CD3CN 1.94 CD3CN 1.93 CD3CN 1.93 CD3CN
1.75 1.73 1.72 1.34 1.32 1.30
Solvent CD3CN
Temperature 298
Number of Scans 16
Spectrometer Frequency 400.13
Nucleus 1H
CF3COO−
OH
BD Yellow
3.67 3.76 4.26 4.93 7.54 2.62 2.80 4.33 4.88 2.07 2.14
f1 (ppm)

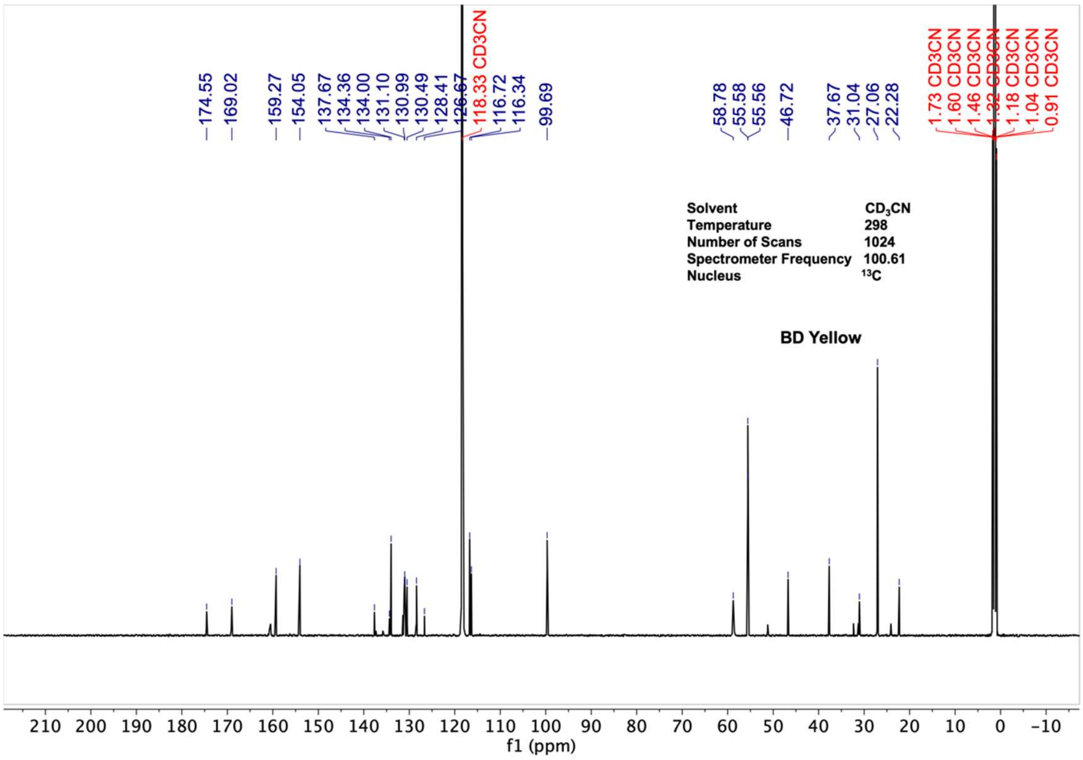
174.55 169.02 159.27 154.05 137.67 134.36 134.00 131.10 130.99 130.49 128.41 126.67
118.33 CD3CN
116.72 116.34 99.69 58.78 55.58 55.56 46.72 37.67 31.04 27.06 22.28
1.73 CD3CN 1.60 CD3CN 1.46 CD3CN 1.32 CD3CN 1.18 CD3CN 1.04 CD3CN 0.91 CD3CN
Solvent CD3CN
Temperature 298
Number of Scans 1024
Spectrometer Frequency 100.61
Nucleus 13C
BD Yellow
f1 (ppm)

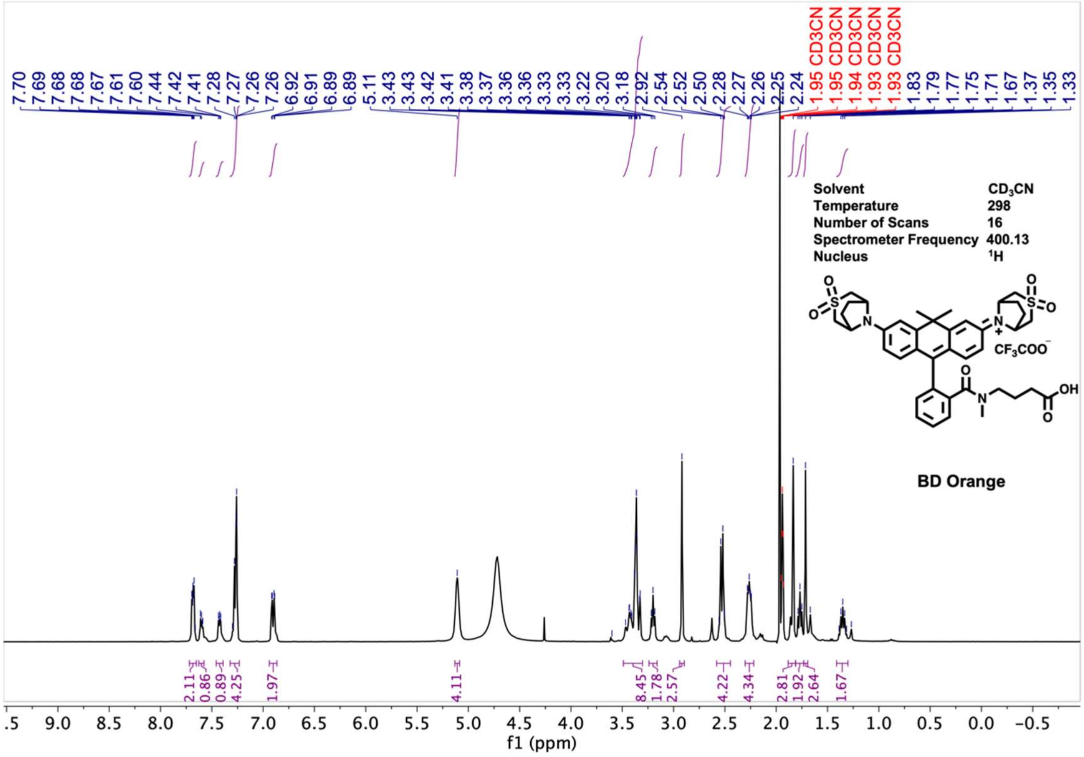

Solvent CD$_3$CN
Temperature 298
Number of Scans 16
Spectrometer Frequency 400.13
Nucleus $^1$H
CF$_3$COO$^-$
BD Orange
f1 (ppm)


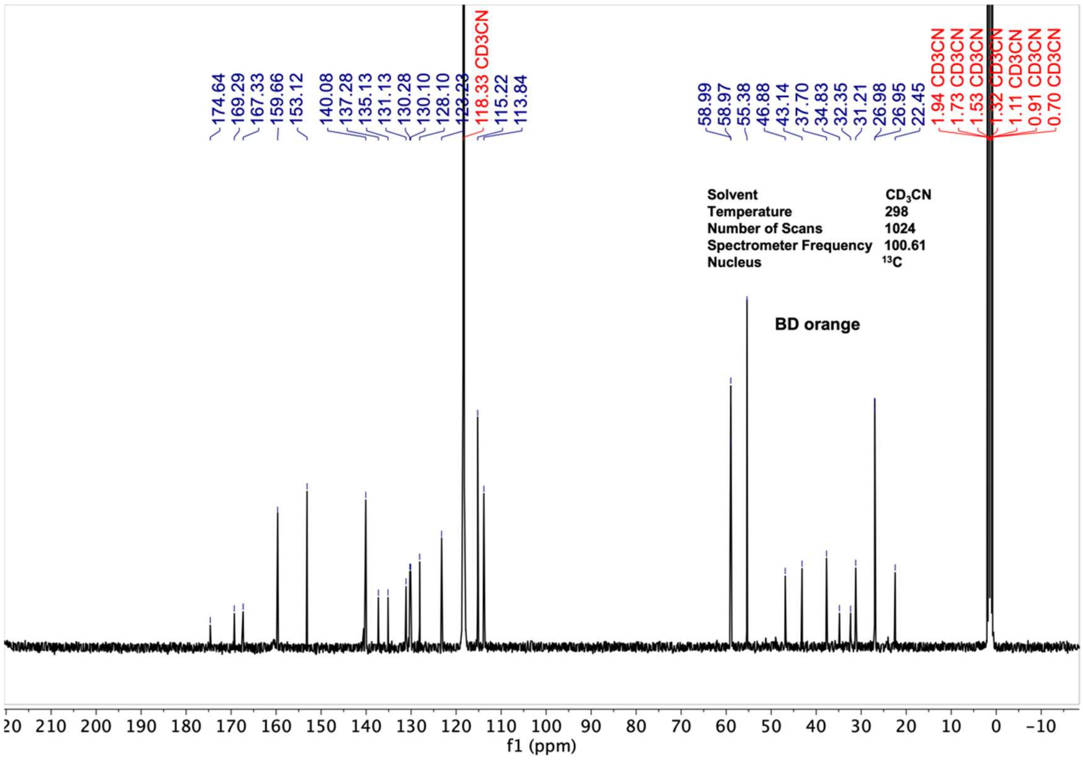

Solvent CD$_3$CN
Temperature 298
Number of Scans 1024
Spectrometer Frequency 100.61
Nucleus $^{13}$C
BD orange
f1 (ppm)

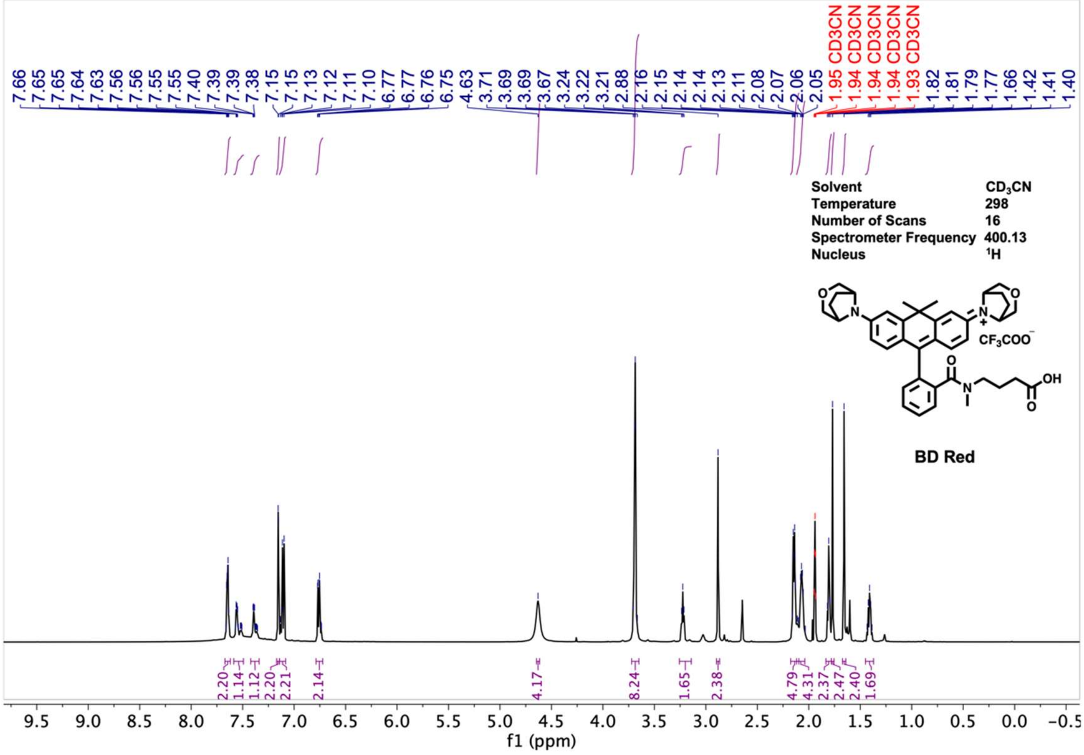
7.66 7.65 7.65 7.64 7.63 7.56 7.56 7.55 7.55 7.40 7.39 7.39 7.38 7.15 7.15 7.13 7.12 7.11 7.10 6.77 6.77 6.76 6.75 4.63 3.71 3.69 3.69 3.67 3.24 3.22 3.21 2.88 2.16 2.15 2.14 2.14 2.13 2.11 2.08 2.07 2.06 2.05
1.95 CD3CN 1.94 CD3CN 1.94 CD3CN 1.94 CD3CN 1.93 CD3CN
1.82 1.81 1.79 1.77 1.66 1.42 1.41 1.40
Solvent CD3CN
Temperature 298
Number of Scans 16
Spectrometer Frequency 400.13
Nucleus 1H
CF3COO−
OH
BD Red
2.20 1.14 1.12 2.20 2.21 2.14 4.17 8.24 1.65 2.38 4.79 4.31 2.37 2.47 2.40 1.69
9.5 9.0 8.5 8.0 7.5 7.0 6.5 6.0 5.5 5.0 4.5 4.0 3.5 3.0 2.5 2.0 1.5 1.0 0.5 0.0 −0.5
f1 (ppm)

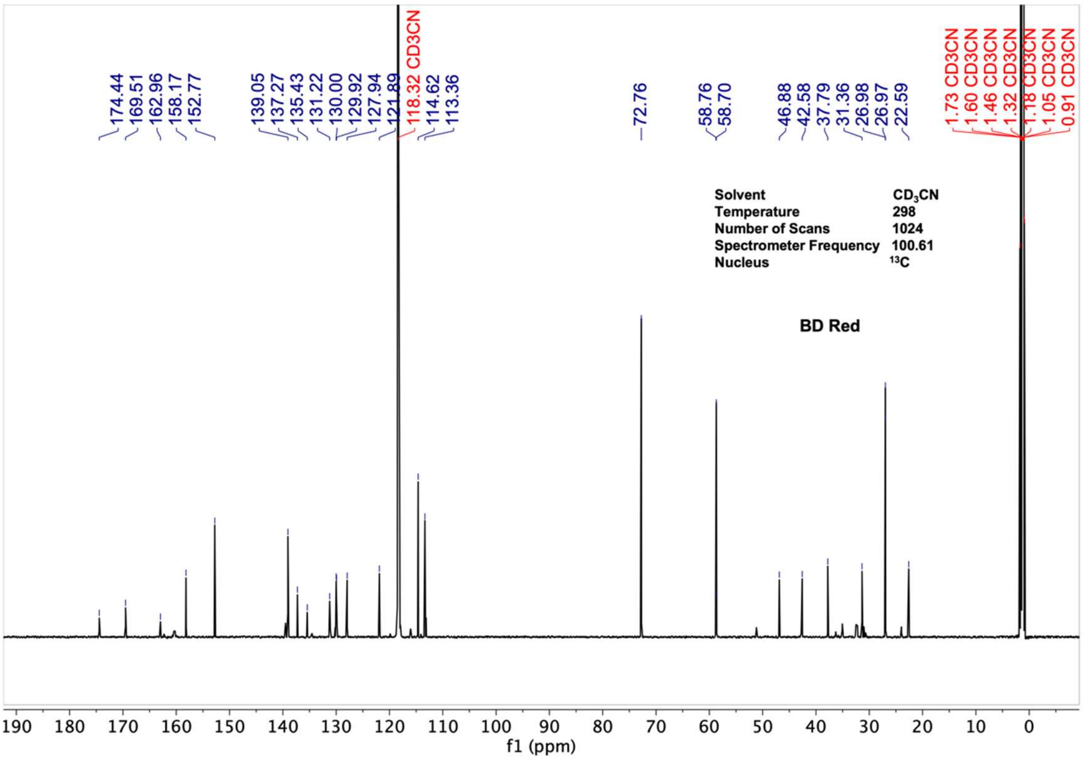
174.44 169.51 162.96 158.17 152.77 139.05 137.27 135.43 131.22 130.00 129.92 127.94 121.89
118.32 CD3CN
114.62 113.36 72.76 58.76 58.70 46.88 42.58 37.79 31.36 26.98 26.97 22.59
1.73 CD3CN 1.60 CD3CN 1.46 CD3CN 1.32 CD3CN 1.18 CD3CN 1.05 CD3CN 0.91 CD3CN
Solvent CD3CN
Temperature 298
Number of Scans 1024
Spectrometer Frequency 100.61
Nucleus 13C
BD Red
190 180 170 160 150 140 130 120 110 100 90 80 70 60 50 40 30 20 10 0
f1 (ppm)

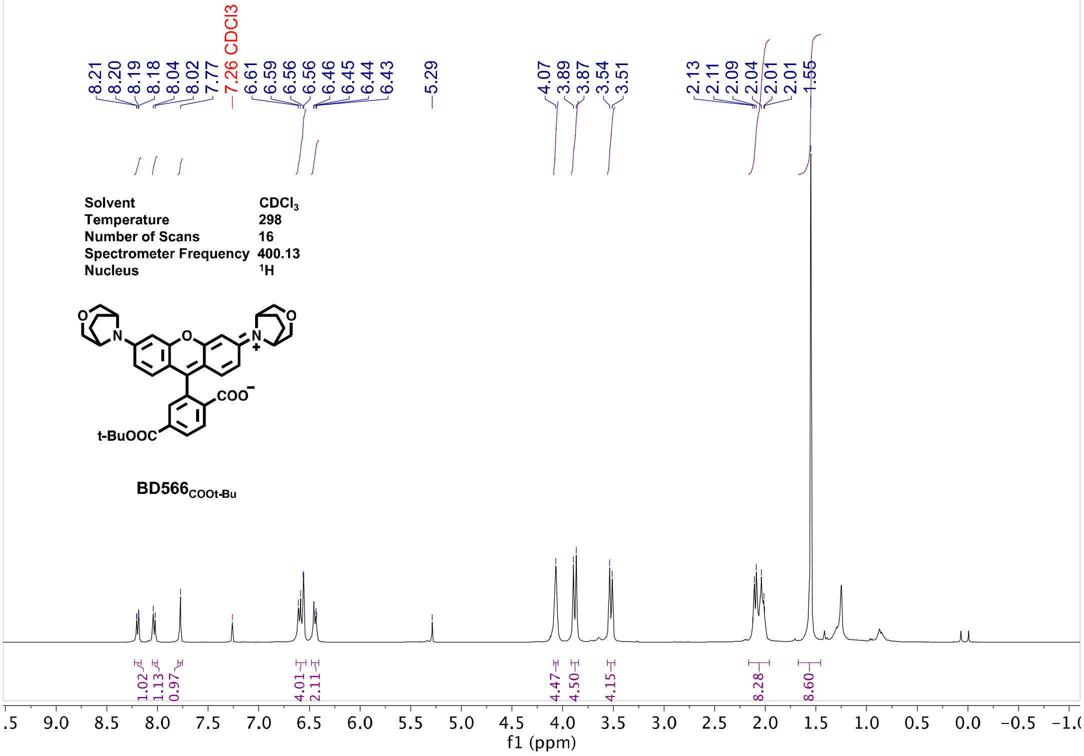

8.21
8.20
8.19
8.18
8.04
8.02
7.77
7.26 CDCl3
6.61
6.59
6.56
6.56
6.46
6.45
6.44
6.43
5.29
4.07
3.89
3.87
3.54
3.51
2.13
2.11
2.09
2.04
2.01
2.01
1.55
Solvent CDCl3
Temperature 298
Number of Scans 16
Spectrometer Frequency 400.13
Nucleus 1H
COO−
t-BuOOC
BD566COOt-Bu
1.02
1.13
0.97
4.01
2.11
4.47
4.50
4.15
8.28
8.60
f1 (ppm)


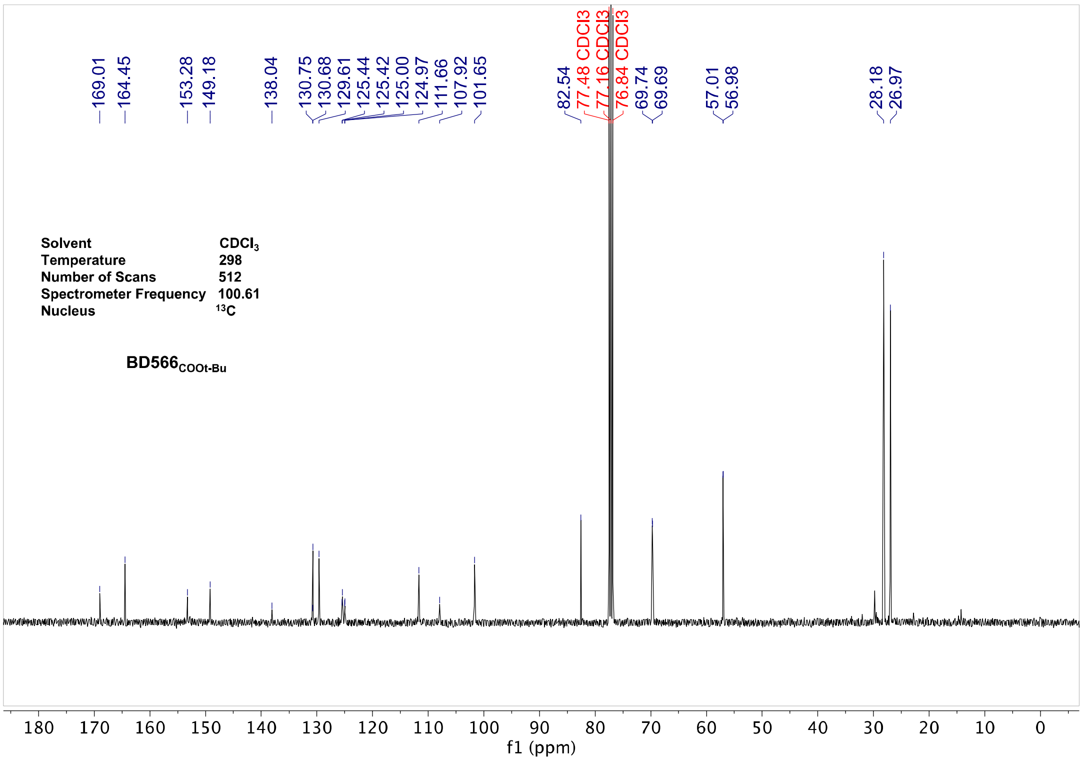

169.01
164.45
153.28
149.18
138.04
130.75
130.68
129.61
125.44
125.42
125.00
124.97
111.66
107.92
101.65
82.54
77.48 CDCl3
77.16 CDCl3
76.84 CDCl3
69.74
69.69
57.01
56.98
28.18
26.97
Solvent CDCl3
Temperature 298
Number of Scans 512
Spectrometer Frequency 100.61
Nucleus 13C
BD566COOt-Bu
f1 (ppm)

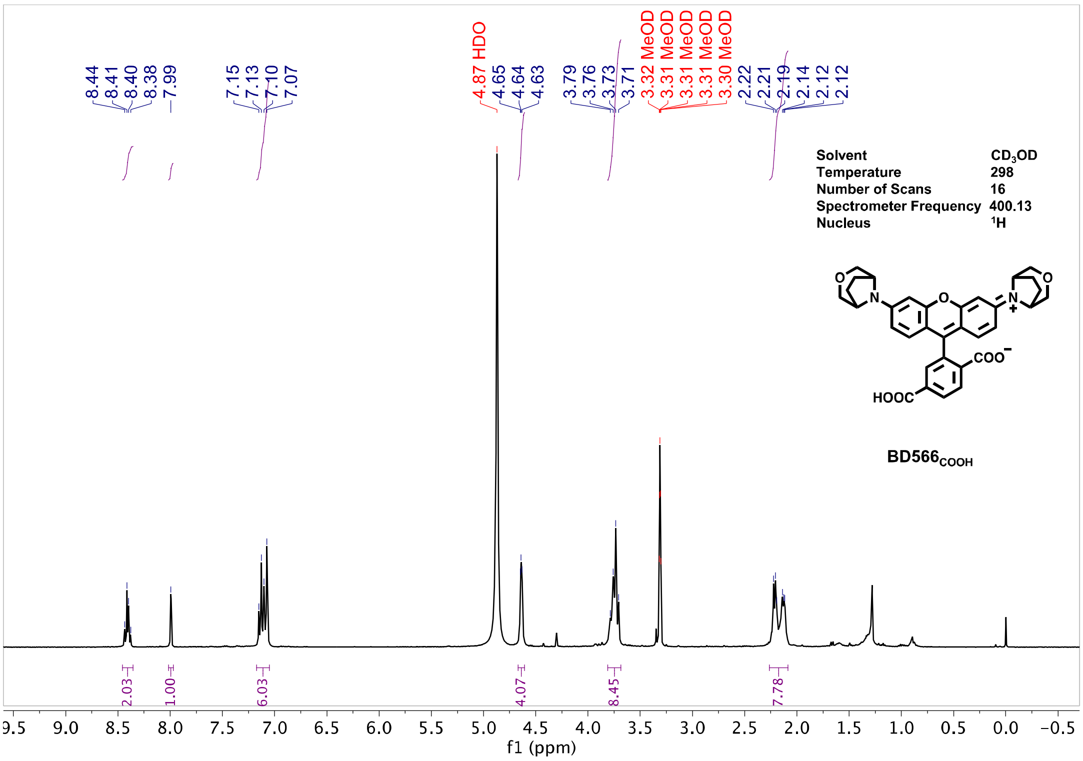
8.44 8.41 8.40 8.38 7.99 7.15 7.13 7.10 7.07 4.87 HDO 4.65 4.64 4.63 3.79 3.76 3.73 3.71 3.32 MeOD 3.31 MeOD 3.31 MeOD 3.31 MeOD 3.30 MeOD 2.22 2.21 2.19 2.14 2.12 2.12
Solvent CD3OD
Temperature 298
Number of Scans 16
Spectrometer Frequency 400.13
Nucleus 1H
HOOC
COO
BD566COOH
2.03 1.00 6.03 4.07 8.45 7.78
f1 (ppm)

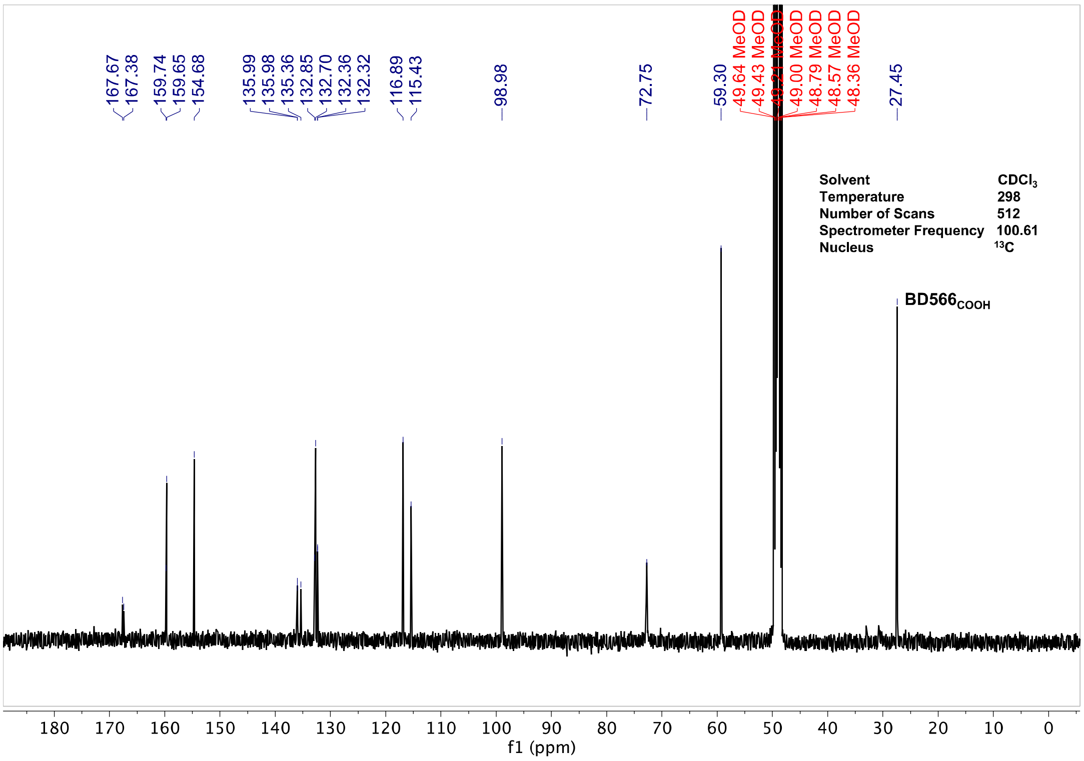
167.67 167.38 159.74 159.65 154.68 135.99 135.98 135.36 132.85 132.70 132.36 132.32 116.89 115.43 98.98 72.75 59.30 49.64 MeOD 49.43 MeOD 49.21 MeOD 49.00 MeOD 48.79 MeOD 48.57 MeOD 48.36 MeOD 27.45
Solvent CDCl3
Temperature 298
Number of Scans 512
Spectrometer Frequency 100.61
Nucleus 13C
BD566COOH
f1 (ppm)

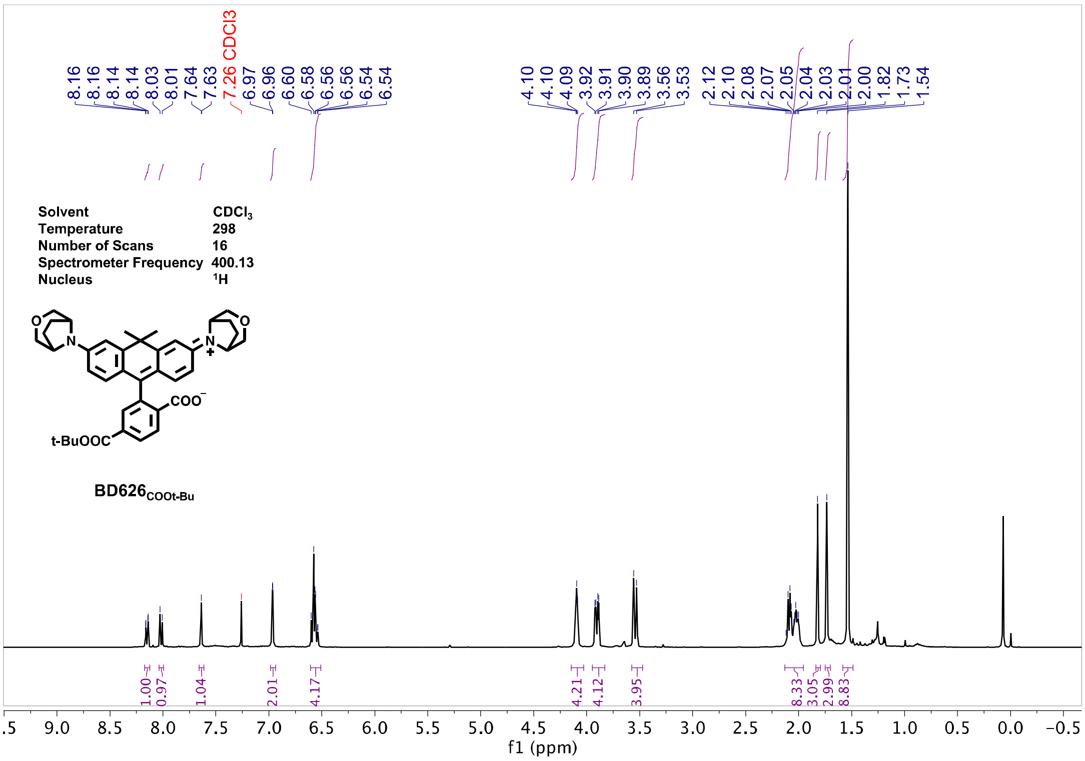
8.16 8.16 8.14 8.14 8.03 8.01 7.64 7.63 7.26 CDCl3 6.97 6.96 6.60 6.58 6.56 6.56 6.54 6.54
4.10 4.10 4.09 3.92 3.91 3.90 3.89 3.56 3.53 2.12 2.10 2.08 2.07 2.05 2.04 2.03 2.01 2.00 1.82 1.73 1.54
Solvent CDCl3
Temperature 298
Number of Scans 16
Spectrometer Frequency 400.13
Nucleus 1H
O
N
N
O
COO−
t-BuOOC
BD626COOt-Bu
1.00 0.97 1.04 2.01 4.17 4.21 4.12 3.95 8.33 3.05 2.99 8.83
f1 (ppm)

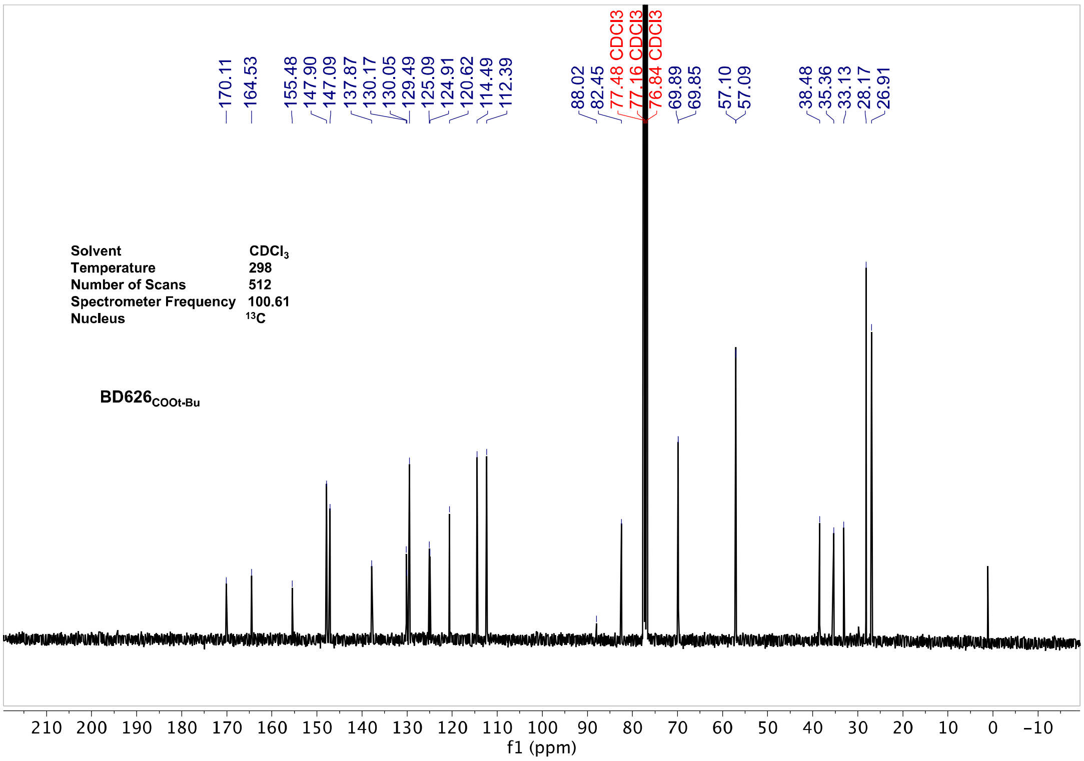
170.11 164.53 155.48 147.90 147.09 137.87 130.17 130.05 129.49 125.09 124.91 120.62 114.49 112.39
88.02 82.45 77.48 CDCl3 77.16 CDCl3 76.84 CDCl3 69.89 69.85 57.10 57.09 38.48 35.36 33.13 28.17 26.91
Solvent CDCl3
Temperature 298
Number of Scans 512
Spectrometer Frequency 100.61
Nucleus 13C
BD626COOt-Bu
f1 (ppm)

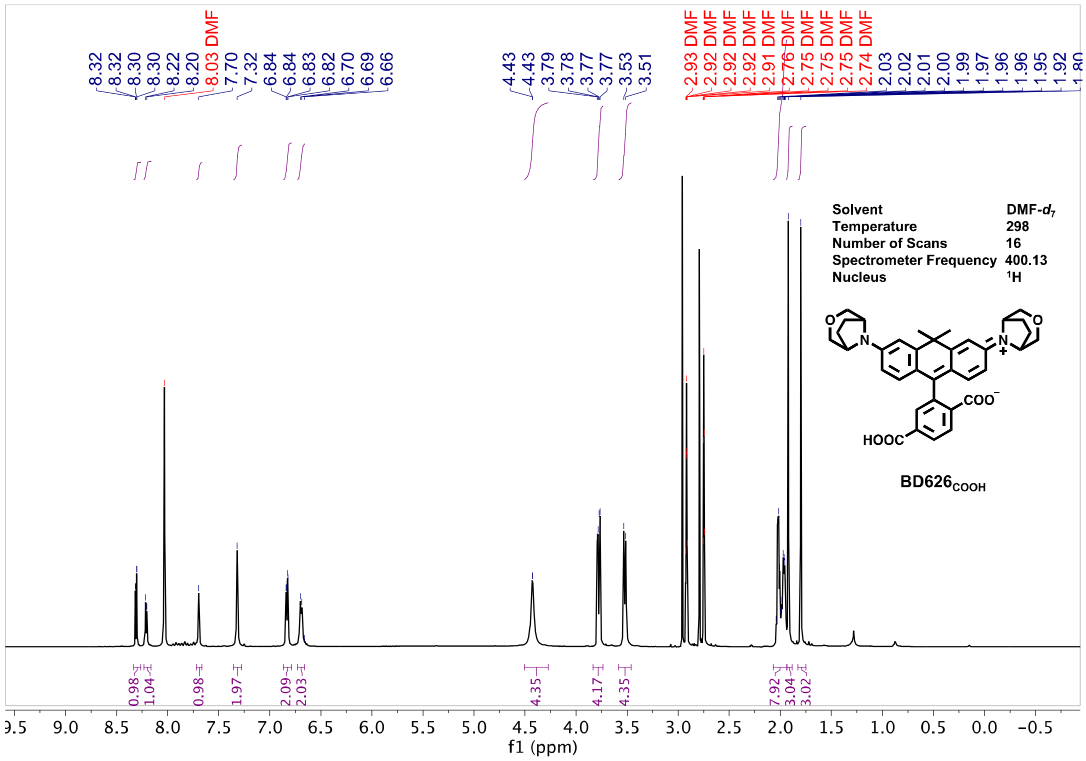
Solvent DMF-$d_7$
Temperature 298
Number of Scans 16
Spectrometer Frequency 400.13
Nucleus $^1$H
BD626$_{COOH}$
f1 (ppm)

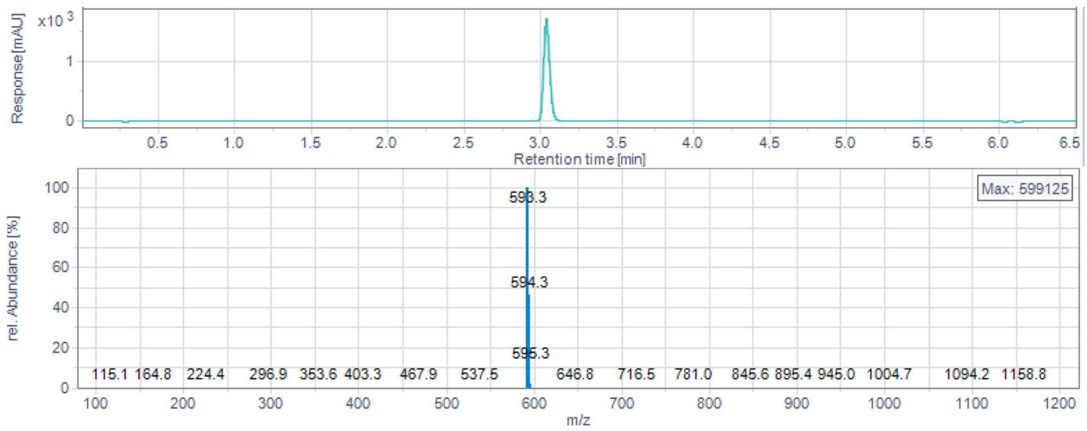
Response [mAU]
Retention time [min]
rel. Abundance [%]
Max: 599125
593.3
594.3
595.3
m/z

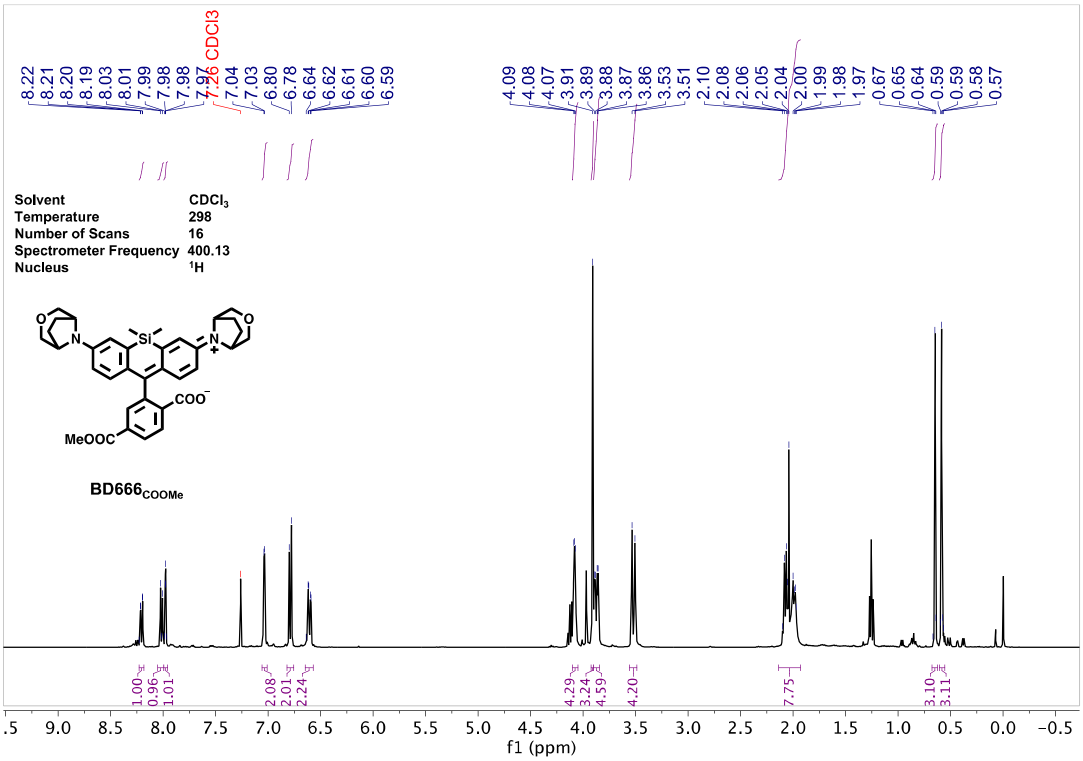

8.22
8.21
8.20
8.19
8.03
8.01
7.99
7.98
7.98
7.97
7.26 CDCl3
7.04
7.03
6.80
6.78
6.64
6.62
6.61
6.60
6.59
4.09
4.08
4.07
3.91
3.89
3.88
3.87
3.86
3.53
3.51
2.10
2.08
2.06
2.05
2.04
2.00
1.99
1.98
1.97
0.67
0.65
0.64
0.59
0.59
0.58
0.57
Solvent CDCl3
Temperature 298
Number of Scans 16
Spectrometer Frequency 400.13
Nucleus 1H
MeOOC
COO−
BD666COOMe
1.00
0.96
1.01
2.08
2.01
2.24
4.29
3.24
4.59
4.20
7.75
3.10
3.11
f1 (ppm)


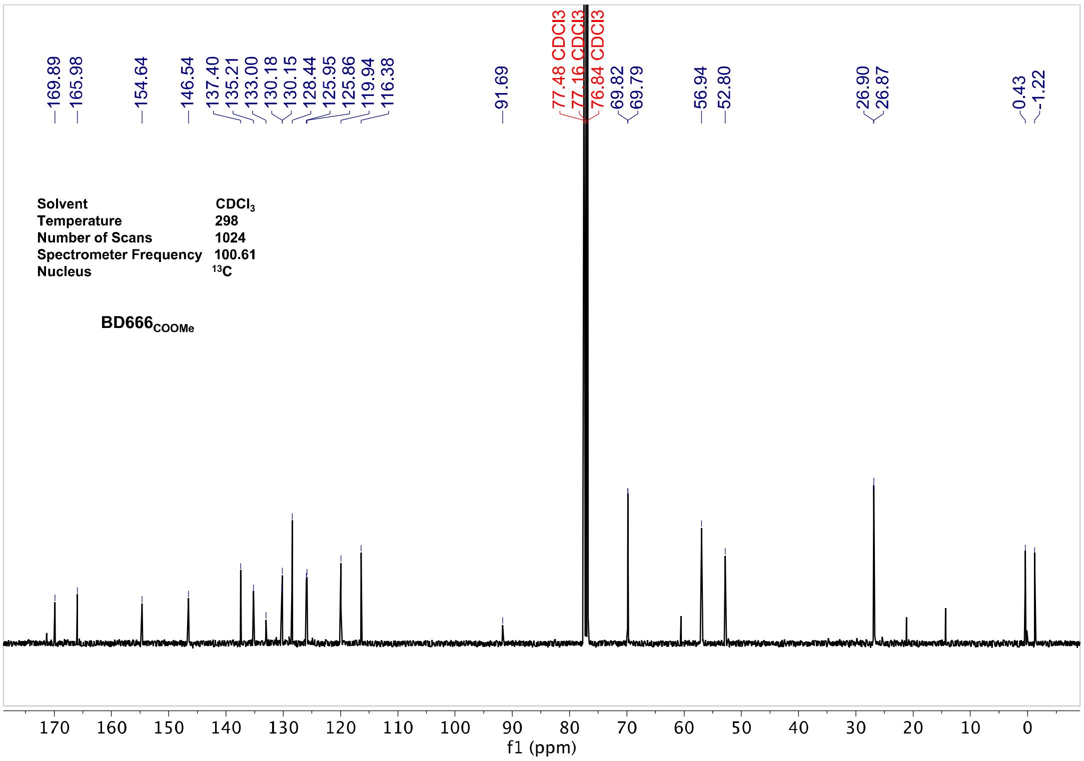

169.89
165.98
154.64
146.54
137.40
135.21
133.00
130.18
130.15
128.44
125.95
125.86
119.94
116.38
91.69
77.48 CDCl3
77.16 CDCl3
76.84 CDCl3
69.82
69.79
56.94
52.80
26.90
26.87
0.43
-1.22
Solvent CDCl3
Temperature 298
Number of Scans 1024
Spectrometer Frequency 100.61
Nucleus 13C
BD666COOMe
f1 (ppm)

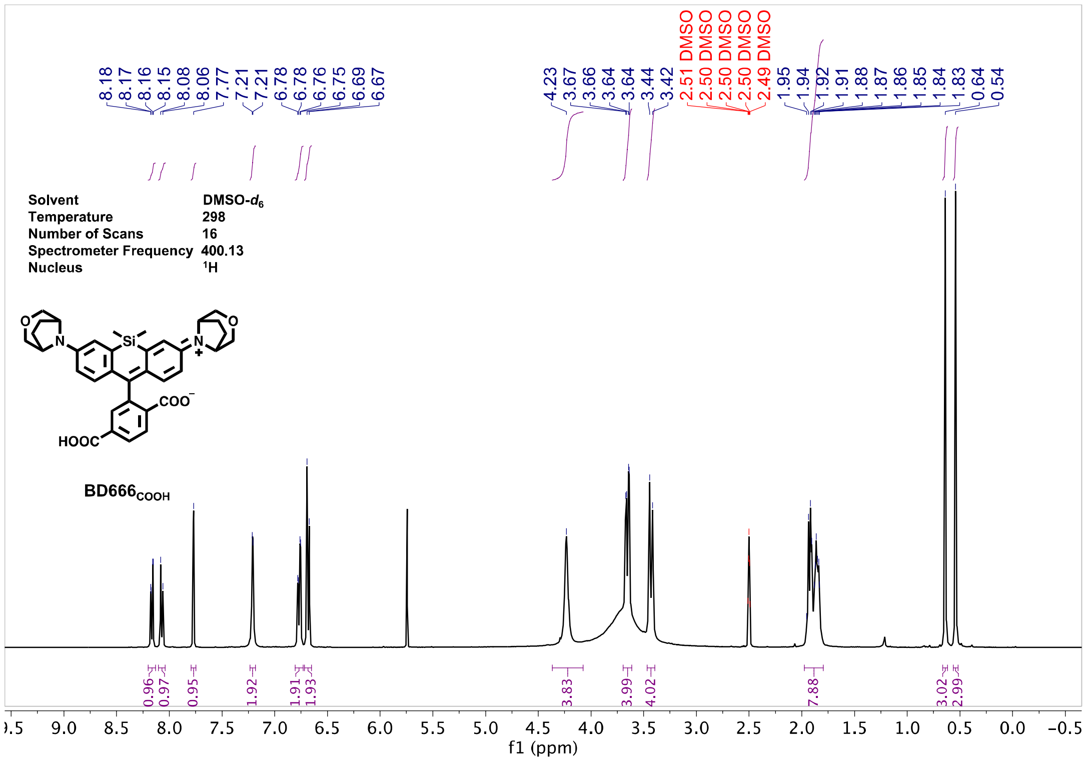
Solvent DMSO-$d_6$
Temperature 298
Number of Scans 16
Spectrometer Frequency 400.13
Nucleus $^1$H
BD666$_{COOH}$
f1 (ppm)

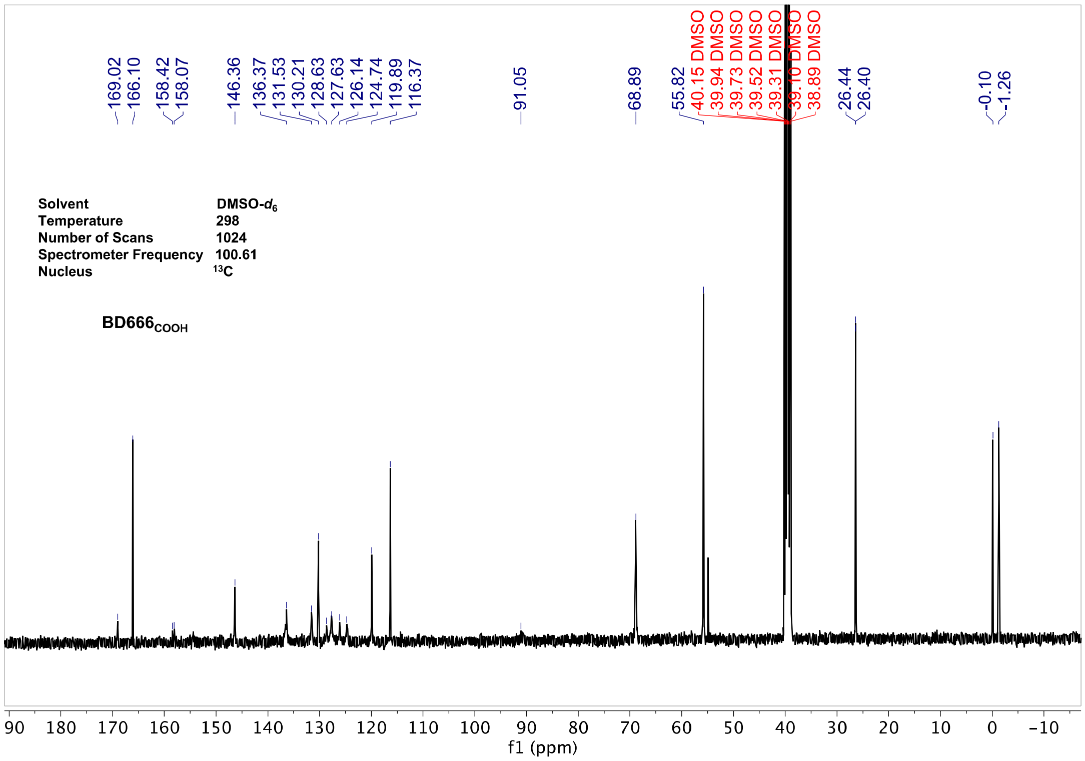
Solvent DMSO-$d_6$
Temperature 298
Number of Scans 1024
Spectrometer Frequency 100.61
Nucleus $^{13}$C
BD666$_{COOH}$
f1 (ppm)

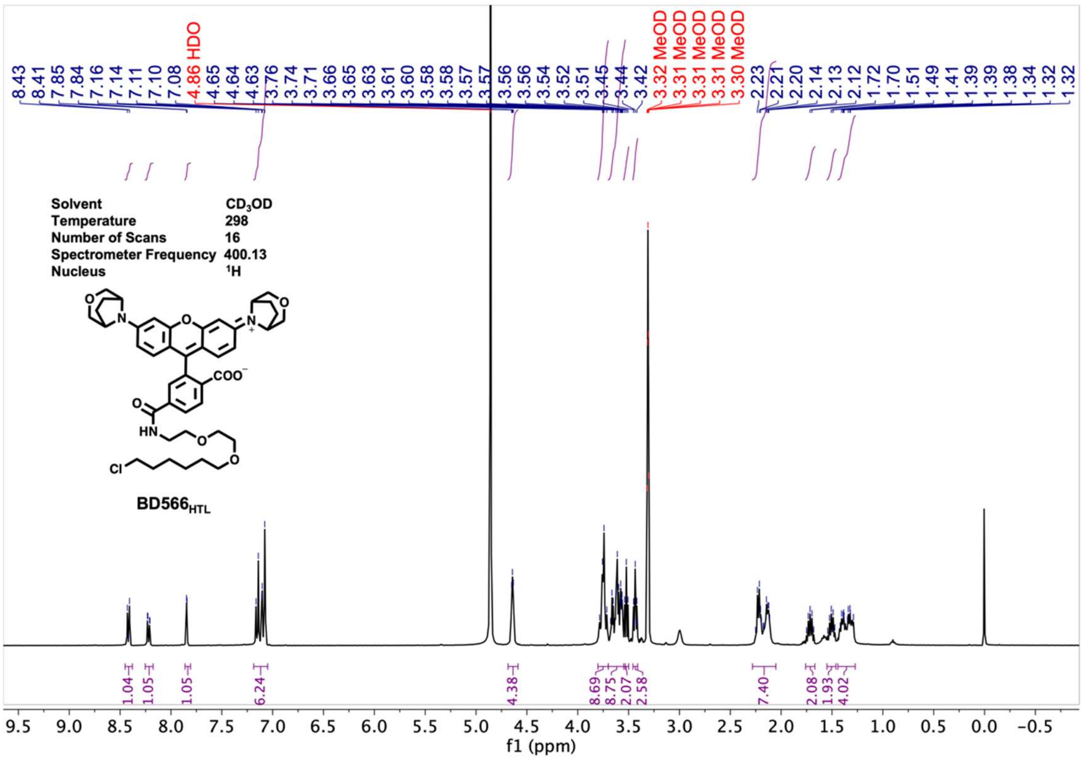

8.43 8.41 7.85 7.84 7.16 7.14 7.11 7.10 7.08 4.86 HDO 4.65 4.64 4.63 3.76 3.74 3.71 3.66 3.65 3.63 3.61 3.60 3.58 3.58 3.57 3.57 3.56 3.56 3.54 3.52 3.51 3.45 3.44 3.42 3.32 MeOD 3.31 MeOD 3.31 MeOD 3.31 MeOD 3.30 MeOD 2.23 2.21 2.20 2.14 2.13 2.12 1.72 1.70 1.51 1.49 1.41 1.39 1.38 1.34 1.32 1.32
Solvent CD3OD
Temperature 298
Number of Scans 16
Spectrometer Frequency 400.13
Nucleus 1H
COO
HN
O
Cl
N
BD566HTL
1.04 1.05 1.05 6.24 4.38 8.69 8.75 2.07 2.58 7.40 2.08 1.93 4.02
f1 (ppm)


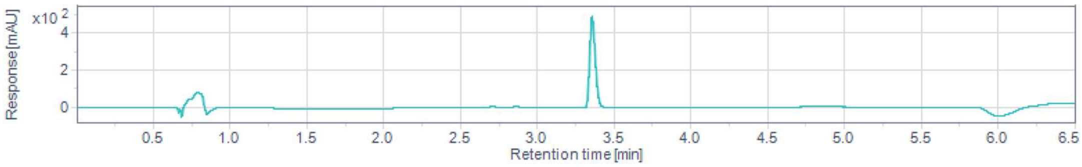

Response [mAU]
x10 2
Retention time [min]


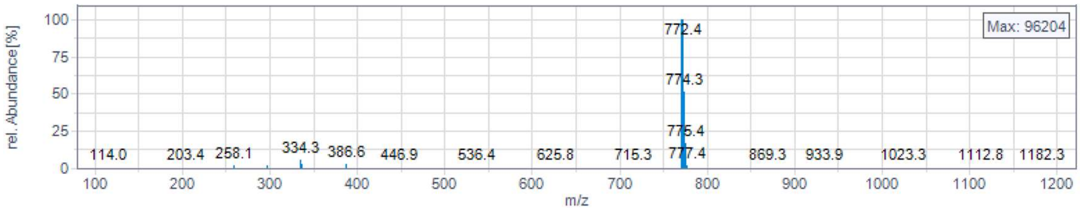

rel. Abundance [%]
Max: 96204
772.4
774.3
775.4
777.4
114.0 203.4 258.1 334.3 386.6 446.9 536.4 625.8 715.3 869.3 933.9 1023.3 1112.8 1182.3
m/z

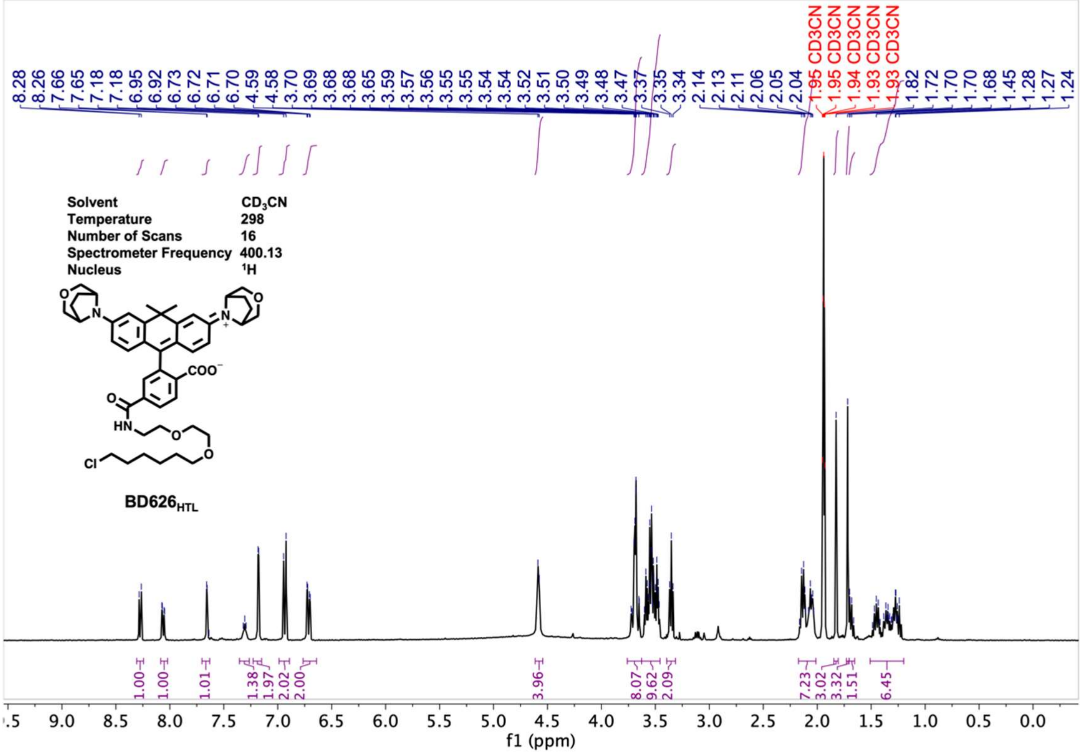

Solvent CD3CN
Temperature 298
Number of Scans 16
Spectrometer Frequency 400.13
Nucleus 1H
BD626HTL
f1 (ppm)


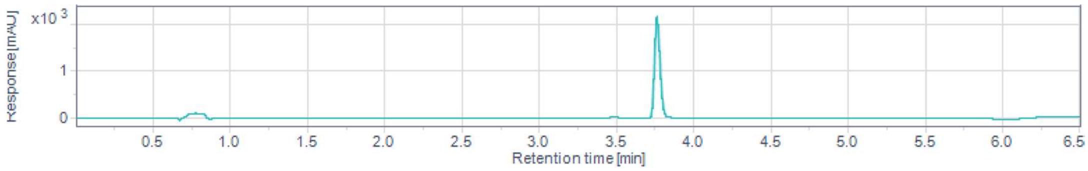

Response [mAU]
x10 3
Retention time [min]


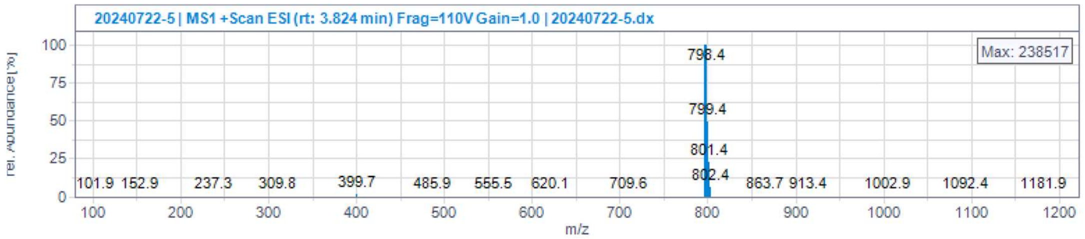

20240722-5 | MS1 +Scan ESI (rt: 3.824 min) Frag=110V Gain=1.0 | 20240722-5.dx
Max: 238517
798.4
799.4
801.4
802.4
Rel. Abundance [%]
m/z

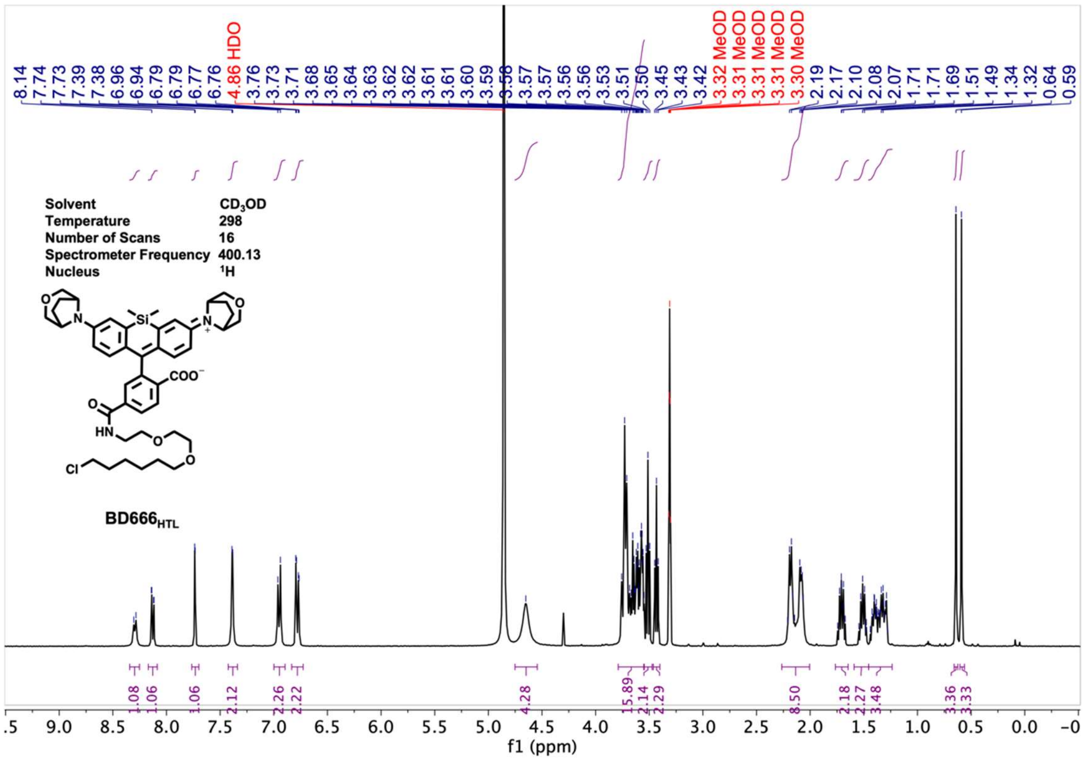
8.14 7.74 7.73 7.39 7.38 6.96 6.94 6.79 6.79 6.77 6.76 4.86 HDO 3.76 3.73 3.71 3.68 3.65 3.64 3.63 3.62 3.62 3.61 3.61 3.60 3.59 3.58 3.57 3.57 3.56 3.56 3.53 3.51 3.50 3.45 3.43 3.42 3.32 MeOD 3.31 MeOD 3.31 MeOD 3.31 MeOD 3.30 MeOD 2.19 2.17 2.10 2.08 2.07 1.71 1.71 1.69 1.51 1.49 1.34 1.32 0.64 0.59
Solvent CD3OD
Temperature 298
Number of Scans 16
Spectrometer Frequency 400.13
Nucleus 1H
Si
N
O
COO−
HN
Cl
BD666HTL
1.08 1.06 1.06 2.12 2.26 2.22 4.28 15.89 2.14 2.29 8.50 2.18 2.27 3.48 3.36 3.33
f1 (ppm)

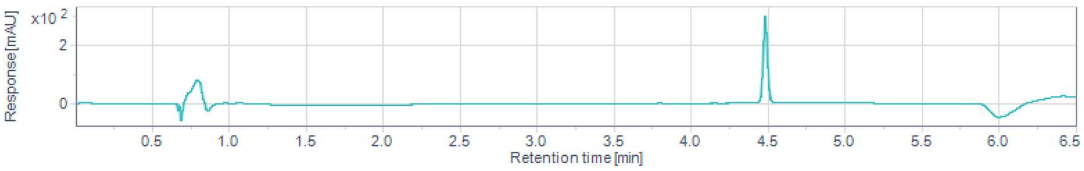
x10 2
Response [mAU]
Retention time [min]

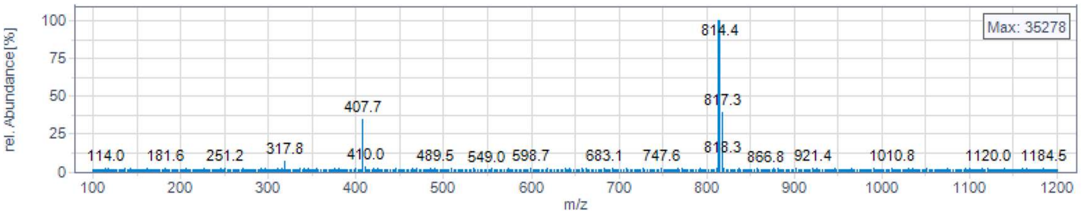
rel. Abundance [%]
Max: 35278
814.4
817.3
407.7
114.0 181.6 251.2 317.8 410.0 489.5 549.0 598.7 683.1 747.6 818.3 866.8 921.4 1010.8 1120.0 1184.5
m/z
